# Evolutionary prediction for new echolocators

**DOI:** 10.1101/2023.09.13.556757

**Authors:** Qi Liu, Qin-Yang Hua, Rui Sun, Shui-Wang He, Meng-Cheng Li, Lu-Ye Shi, Peng Chen, Yuan-Shuo Ma, Qin Zhang, Xue-Long Jiang, Yong Wang, Peng Shi

## Abstract

It was suggested over 60 years ago that echolocation is widespread in animals. Although echolocation has been found to evolve independently in several mammalian lineages, this hypothesis remains largely untested due to the difficulty of determining the presence of echolocation. To address this issue, we combined molecular convergence analyses from 190 high-quality mammalian genomes with machine learning to predict potential new mammalian echolocators. Our model predicted three promising lineages of echolocating mammals. Behavioral experiments confirmed that the gracile shrew mole (*Uropsilus gracilis*), the highest- ranking species among predicted echolocators, as well as almost all shrew moles (Uropsilinae), are capable of echolocation through the use of ultrasonic pulses. In contrast to most bats that use laryngeal echolocation, the characteristics of calls, the morphology of the stylohyal bone, and the results of tongue ligation experiments all suggest that shrew moles produce ultrasonic clicks with their tongues for echolocation. Finally, we estimated at least 20% of all living mammalian orders with echolocation ability, thus empirically supporting Griffin’s hypothesis that echolocation is widespread among animals. Our findings not only provide evidence that three novel lineages of echolocating mammals, but also demonstrate that phylogenetically replicated phenotypes can be predicted through genetic convergence.

**One sentence summary:** Shrew moles are capable of echolocation.

## Main Text

Since coined the term ‘echolocation’ through the observation of bats using their sounds and hearing to detect obstacles in 1938 (*1*), Griffin further hypothesized that echolocation could be widespread among animals(*2*), likely due to the adaptation of many species to vision-ineffective environments with morphological characteristics related to echolocation, such as reduced eyes, enlarged outer ear, and high-frequency calls. Echolocation has been found to have evolved across multiple lineages of vertebrates, including birds (*3*), bats, whales, shrews (*4*), tenrecs (*5*), and soft-furred tree mice (*6*). In particular, echolocation has independently evolved at least five times in mammals, with its origin potentially stemming from the last common ancestors of bats and toothed whales (*7, 8*), making it a noteworthy example of the evolutionary repeatability of phenotypes. Nevertheless, the validity of Griffin’s hypothesis lacks systematic examination. This is due to the complexity of echolocation as an orientation behavior, which makes it difficult to ascertain whether an animal uses echolocation based on morphological characters, detailed field observation in various natural habitats, or behavioral experiments across thousands of mammalian species.

Accumulating evidence supports the notion that adaptive morphology, physiology, and ethology often converge owing to the molecular repeatability of convergent amino acid substitutions under the pressure of natural selection (*9–11*). In particular, increasing evidence indicates that echolocating mammalian species or groups have undergone adaptively convergent amino acid replacements in a number of genes (*12–17*). A comparison of genome sequences revealed a substantial molecular convergence in hearing-related proteins among echolocating bats, toothed whales, and soft-furred tree mice (*6*). Furthermore, it is strongly suggested that molecular repeatability is closely associated with the origins and functions of the convergent phenotype of echolocation among mammalian echolocators, which motivates us to combine molecular convergences among echolocating mammalian species with machine learning for predicting novel echolocating lineages among hundreds of mammals with available genomic sequences.

### Machine learning aided new echolocator detection

To identify the molecular convergences that can efficiently differentiate echolocating and nonecholocating mammals, a total of 9,554 one-to-one orthologous proteins were collected across 93 mammalian species (**Fig. S1; Tables S1, S2**), including 35 echolocating mammals from four separate orders: one rodential soft-furred tree mouse (*Typhlomys cinereus*), one tenrecid lesser hedgehog tenrec (*Echinops telfairi*), 14 insectivorous bats, and 19 toothed whales. Ancestral amino acid sequences were inferred using the maximum likelihood method (*18*) based on the phylogeny of these species, and convergent amino acid substitutions among the echolocating lineages were identified. As echolocation is suggested to have originated in the last common ancestor (LCA) of bats and the LCA of toothed whales, respectively (*7, 8*), convergent amino acid substitutions were counted not only among living echolocating species from four lineages, but also among the soft-furred tree mouse, the lesser hedgehog tenrec, the LCA of bats, and the LCA of toothed whales. In total, 497 convergent amino acid substitutions were identified. To ensure the relevance of these molecular convergences to echolocation, a convergent amino acid substitution was excluded if it occured in the 53 non-nocturnal or known nonecholocating mammals (**Table S2**), as the non-nocturnal species mainly use vision to explore environments and thus are not expected to possess the ability of echolocation. Finally, 14 convergent amino acid substitutions with high confidence among known echolocating mammalian lineages were obtained (**Table S3**), and these sites were found to be effective in differentiating echolocating and nonecholocating mammals (**Fig. 1A; Fig. S2**).

**Fig. 1.**
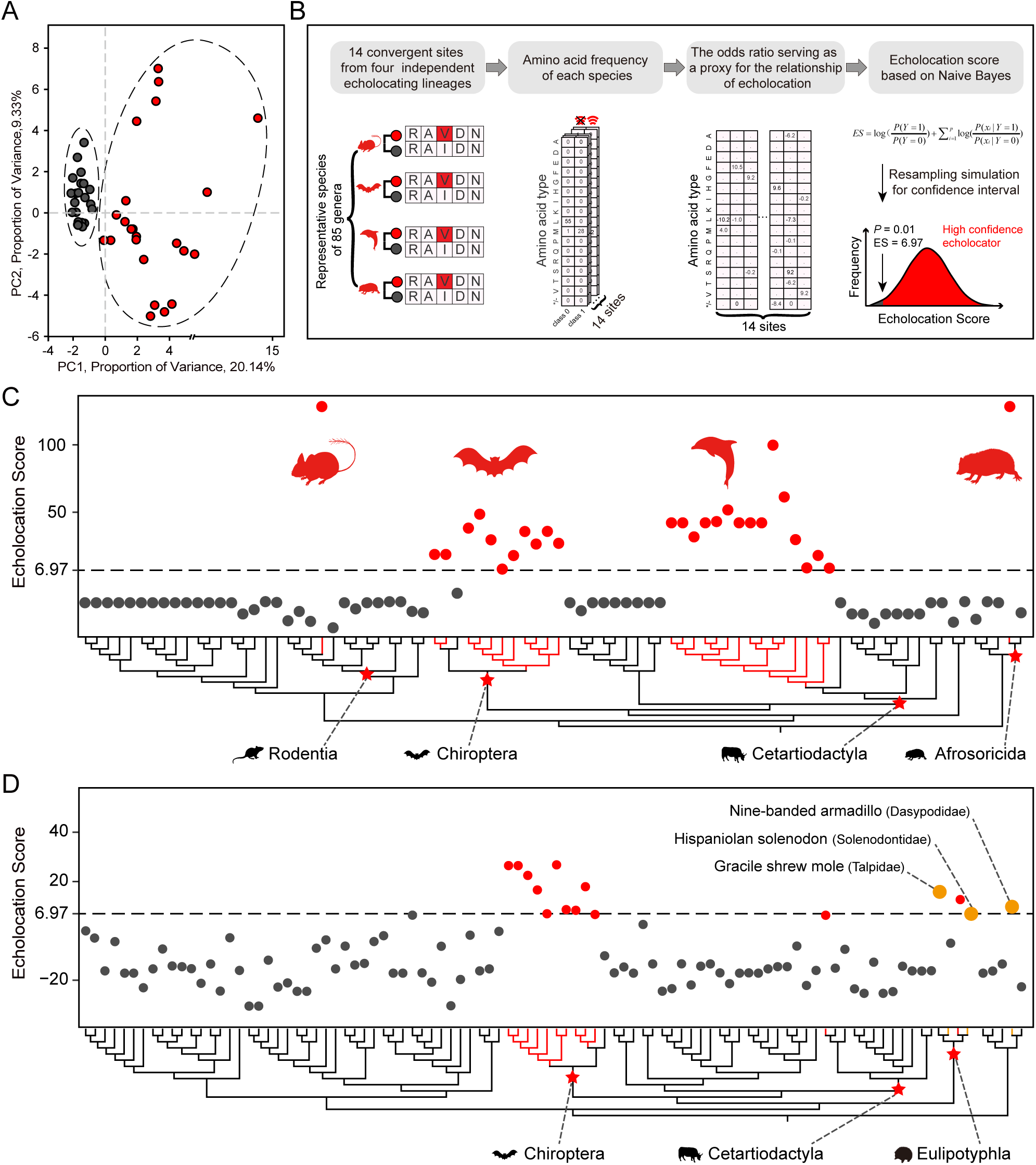
Combine molecular convergences with machine learning to predict potential new mammalian echolocators. (**A**) Principal component analysis used 14 amino acid substitutions from known echolocating mammals can differentiate the echolocating and nonecholocating mammals. (B) Schematic diagram shows workflow of machine learning and algorithm of Echolocation scores (ES) (see details in materials and methods). (**C**) Echolocation scores with stricter cutoff ES > 6.97 can stand out all echolocating species from training dataset based on machine learning. (**D**) Combine machine learning with molecular convergences predict three putative echolocating mammalian species. The red dots represent echolocators, black dots represent non-echolocators, pink dots represent putative echolocators, red pentagrams mark orders possess the ability of echolocation up to now. Convergent amino acid substitutions across representative echolocating and non-echolocating mammalian genera are shown in **Table S13**, ES score of each species are shown in **Table S4**.

To properly model the polygenic architecture and determine whether a particular mammalian species is an echolocator, we evaluated 5 different machine learning algorithms and selected Naïve Bayes to generate an echolocation score (ES) by integrating genomic features of 14 convergent amino acid substitutions across 85 representative echolocating and non-echolocating mammalian species for each genus (**Fig 1B; Fig. S3**). The prediction of echolocator (labeled Y = 1 if echolocator, and 0 otherwise) for each species was based on the calculation of the posterior odds of echolocation given the presence of genomic features *x*_1_, *x*_2_, …, *x*_*n*_ (n=14) as follows using the Bayes rule:

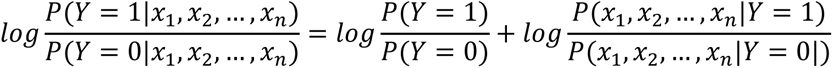

*P*(*Y* = 1|*x*_1_, *x*_2_, …, *x*_*n*_) is the probability that the mammal is an echolocator given these features. P(Y=1)/P(Y=0) is the prior odds. *P*(*x*_1_, *x*_2_, …, *x*_*n*_|*Y* = 1) / *P*(*x*_1_, *x*_2_, …, *x*_*n*_|*Y* = 0) is the likeli-hood ratio for the combined features. Specifically, we first constructed a type frequency table for each of 14 convergent amino acid substitutions that emerged in echolocating and nonecholocating groups, respectively (**Fig. S4**). We then calculated the likelihood ratio of the type frequency for each amino acid along the concatenated convergent sites. The likelihood ratio of the *i*-th amino acid was determined by dividing the number of one certain amino acid *x_i_* relative to the total number of amino acids in the echolocating group (Y=1) by the number of the amino acid *x_i_* relative to the total number of amino acids in the nonecholocating group (Y=0).

A mammal species is predicted to be an echolocator if the posterior odds calculated is greater than a pre-established cutoff. Assuming that genomic features are conditionally independent given echolocation, the likelihood ratio of the combined features (ES value) is equal to the product of the likelihood ratios for individual features. Upon calculation, it was found that the ES was greater than 0 for all 28 echolocating mammals and less than 0 for the remaining 57 nonecholocating mammals in the training dataset (**Fig 1C; Table S4**). We pursued a stricter cutoff by randomly selecting an amino acid from each of 14 convergent sites among 28 echolocating mammals to create a simulated dataset and calculate its ES. This procedure was repeated 10,000 times to obtain a null distribution of ES values from the simulated datasets. According to the statistical test, the probability that a species with ES > 6.97 is not an echolocator is less than 1% (**Fig. 1B; Fig. S5**). To screen new echolocating mammalian species, we focused on 97 mammalian species, of which 74 species with relatively high-quality genomic data from the Zoonomia Project (**Table S5**) (*19*) and 23 were nocturnal species from the database OrthoMaM (**Table S6**) (*20*). Notably, this dataset included 12 known echolocating mammals, such as echolocating bats, toothed whales, and the European shrew (*Sorex araneus*), which were used as positive controls. By calculating ES values of these species, we found that the ES values of 9 out of 12 known echolocating mammals were larger than the cutoff 6.97 and those of the remaining 3 echolocating mammals were close to the cutoff (**Fig. 1D**; **Table S7**), further indicating the validity of the machine learning model. In addition to these echolocating mammals, two species had ES values greater than 6.97, the gracile shrew mole (*Uropsilus gracilis*) with ES = 15.88 and the nine-banded armadillo (*Dasypus novemcinctus*) with ES = 9.88, as well as one species with ES close to the cutoff, the solenodon (*Solenodon paradoxus*) with ES = 6.88 (**Fig. 1D**; **Table S7**), suggesting that these species may be potential new echolocators in mammals.

### Echolocation in shrew moles

As the highest-ranking species according to ES, the gracile shrew mole possesses additional features associated with echolocation, such as nocturnal habit, reduced eyes, and large external ears, which further suggests that this species may use echolocation (*21*). It is noted that all eight recognized species of shrew moles (subfamily Uropsilinae) share similar ecological and morphological traits (**Fig. 2A, 2B**) (*21–23*), leading to the expectation that echolocation may be present across all shrew moles if the gracile shrew mole is an echolocator. To assess this hypothesis, we initially recorded vocalizations of four species of shrew moles while they moved freely in the dark. We found that all four species produced short, broadband ultrasonic vocalizations (USVs) consisting of different types of pulse groups, including singles, dyads, and triads (**Fig. 2C**). The acoustic variables of the USVs were largely similar among these species, with pulse duration of ∼0.8 ms, frequency ranging from ∼32 to ∼101 kHz, bandwidth of ∼69 kHz, and peak frequency of ∼56 kHz (**Fig. 2D, 2E**; **Table S8**). These regularly emitted USVs, as in echolocating mammals, suggest important biological implications and that all shrew moles have evolved echolocation.

**Fig. 2.**
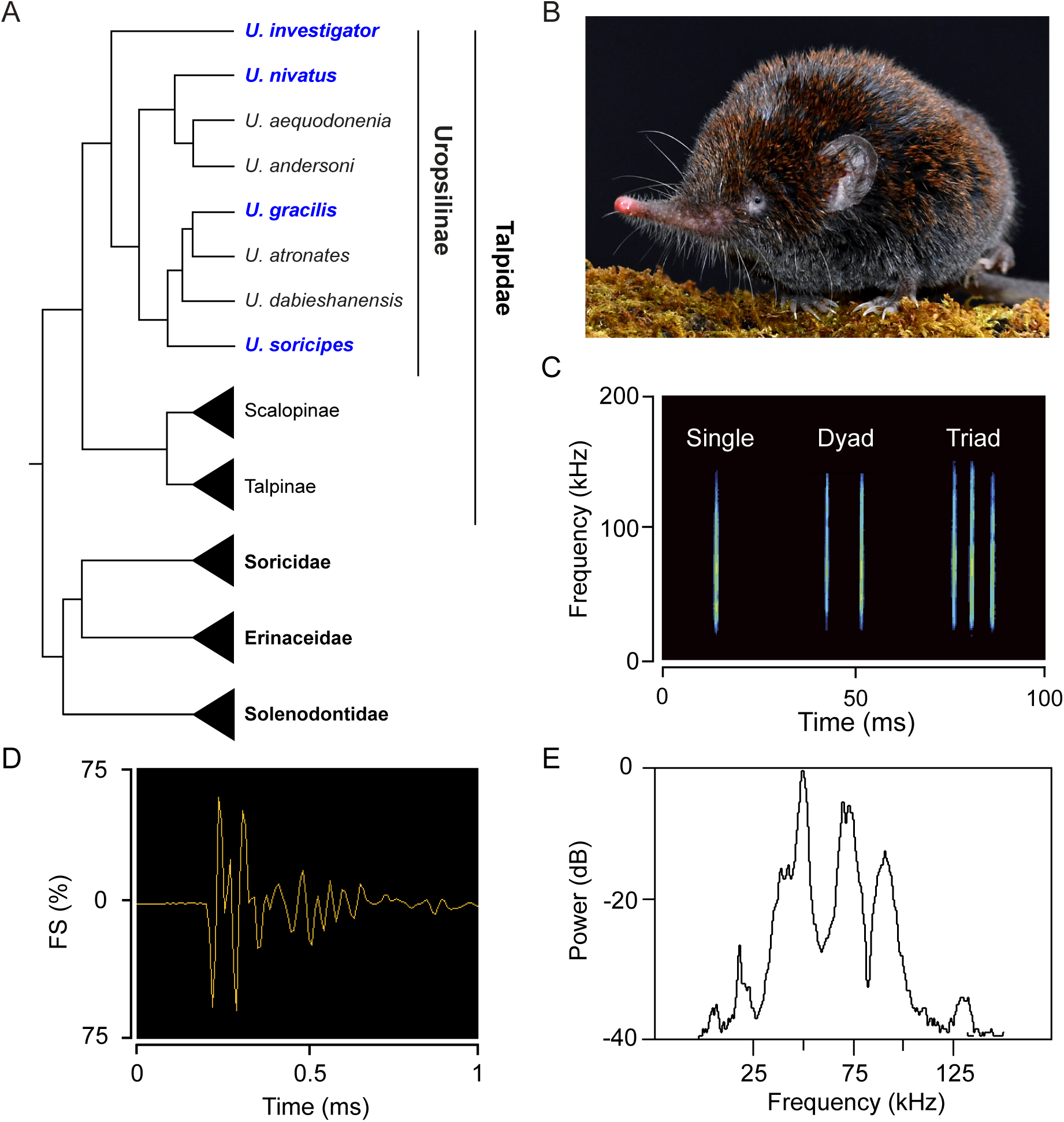
Morphological traits and acoustic characteristics of the shrew moles. (**A**) There are eight recognized shrew mole species (genus *Uropsilus*) in the family Talpidae, and four representative species (purple) of this genus were investigated in this study. (**B**) Morphology of an adult gracile shrew mole (*U. gracilis*). (**C**) Gracile shrew mole produce various types of pulse groups. Spectrum (**D**) and waveform (**E**) of single ultrasonic pulse from the gracile shrew mole.

To experimentally determine whether shrew moles possess the ability of echolocation, we performed a series of behavioral experiments with the disc-platform apparatus that was previously used to test echolocation in terrestrial mammals such as shrews (*4, 24*), tenrecs (*5*) and soft-furred mice (*6*). First, a disc-small disc circle was built as a control, which mainly consisted of a raised central disk surrounded by a circle of eight small discs (**Fig. 3A**). The central disc was divided into eight equal sections, each of which corresponded to one of the small discs. A microphone was randomly placed in one of the small discs prior to each trial in order to record the sonic pulses. If an animal was placed on the central disk to explore, it was expected to spend a comparable amount of time and emit a similar number of pulses in each sector. Subsequently, the small-disc circle was replaced with a platform that was connected to a reward box by a ramp, and the position of the platform was randomly determined before each trial. If echolocation is utilized by the animal, then it should: (i) increase its exploration time and emit more sonic pulses in the sector over the platform, thus descending to the platform; (ii) lose its preference for the over-platform sector and fail to land on the platform when its ears were plugged; and (iii) regain its preference for the over platform sector when the blockages were removed or when plastic tubes that enable hearing were inserted. Four species of shrew moles were used as representatives, namely the gracile shrew mole (*U. gracilis*), the inquisitive shrew mole (*U. investigator*), the snow mountain shrew mole (*U. nivatus*), and the Chinese shrew mole (*U. soricipes*), which cover the main clades of shrew moles (**Fig. 2A**).

**Fig. 3.**
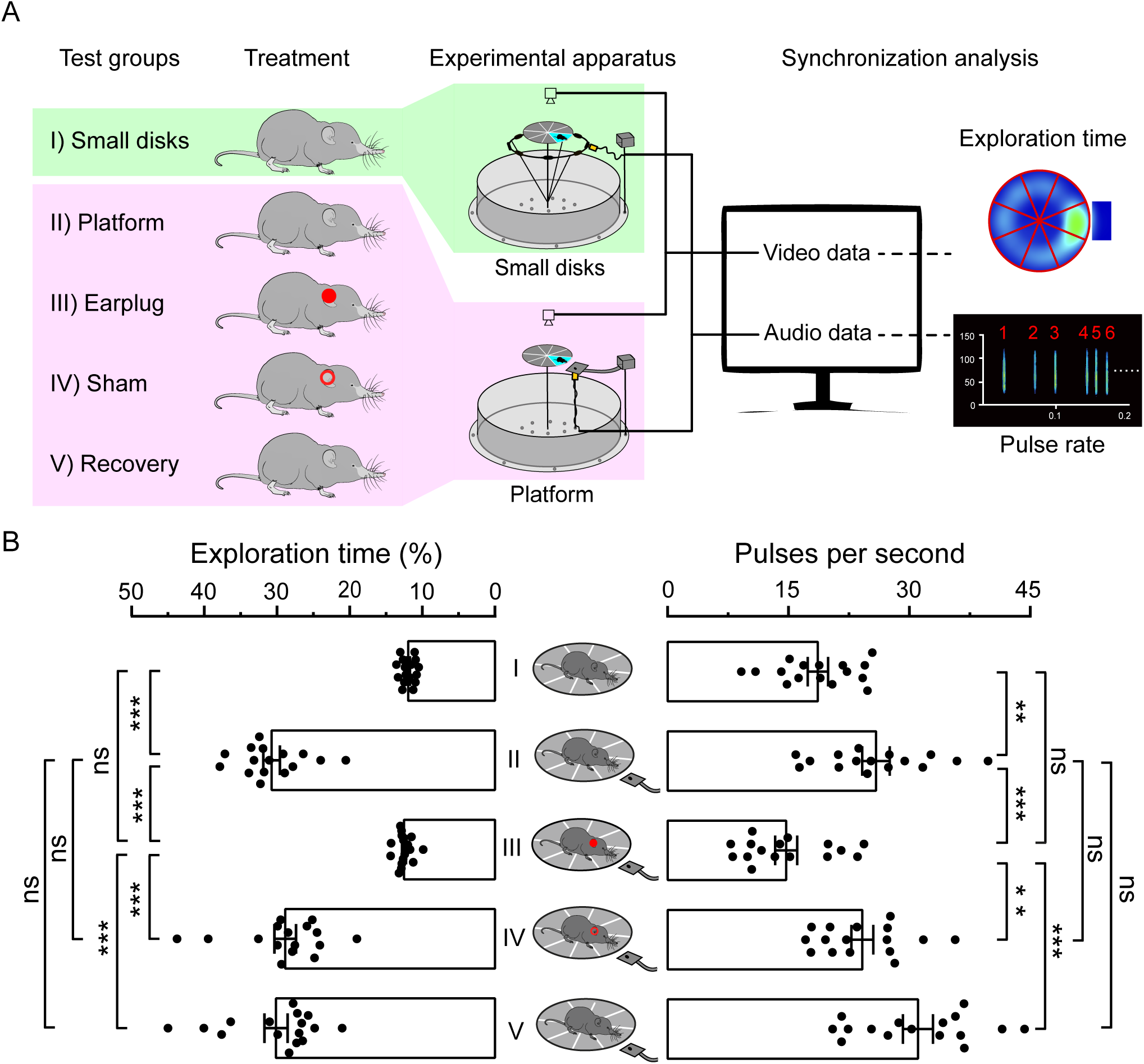
Behavioral experiments for testing echolocation in the shrew moles. (A) Schematic diagram of the behavioral experiments with shrew moles (see details in materials and methods). (**B**) Comparisons of the percentage of exploration time and pulse rates in the sector corresponding to the ultrasonic probe when the gracile shrew moles explored the central disc under various experimental treatments. All data were obtained from *U. gracilis*. Each dot represents the mean of the measurements of each individual (N = 16). The bar denotes the mean value across all individuals, and the error bars represent SEM. *P* values are from two-tailed paired Student’s *t* tests and were corrected with the Bonferroni method. ** *p* < 0.01; *** *p* < 0.001. ns, not significant.

When the gracile shrew moles were placed in the disc-small disc circle apparatus, they spent a similar amount of time (11.96 ± 0.24 % of the total exploration time; N = 16 animals) and produced a similar number of ultrasonic pulses per second within each sector of the central disc in darkness (18.64 ± 1.26; N = 16 animals; **Fig. 3B**). By comparison, when in the disc-platform apparatus, the gracile shrew moles spent more time and emitted more ultrasonic pulses in the sector over the platform (*P* < 0.01; two tailed paired Student’s *t* tests; N = 16 animals; **Fig. 3B**), successfully detecting and dropping down to the lower platform from the central disc. However, the preference of the gracile shrew moles for the over-platform sector was lost when their ears were blocked by earplugs, performing similarly in the control tests (**Fig.3B**). Following the removal or replacement of the earplugs with hollow plastic tubes, the gracile shrew moles regained the preference for the over-platform sector, spending significantly more exploration time and producing more USVs (*P* < 0.01; **Fig. 3B**).

To further verify the use of echolocation by the gracile shrew moles, we compared the preference for the over-platform sector with its adjacent sectors. If the animals echolocate, it was expected that they would spend more time in the over-platform sector than its adjacent sectors. Indeed, we found that the gracile shrew moles spent significantly more exploration time in the over-platform sector (30.75 ± 1.13 %) than in its adjacent sectors (11.79 ± 0.61%; *P* < 0.001; **Fig. S7**). Furthermore, when the ears were occluded, the gracile shrew moles lost their preference for the over-platform sector in comparison with its adjacent and were unable to locate the platform beneath the central disc (*P* > 0.05; **Fig. S7**). Upon the removal of the earplugs, the gracile shrew moles regained the preference for the over-platform sector (*P* < 0.01; Fig. S3). It is noteworth that the other three species of shrew moles, *U. investigator*, *U. nivatus*, and *U. soricipes*, exhibited very similar performance in the aforementioned behavioral experiments (**Figs. S8, S9**, and **S10**; **Tables S9, S10** and **S11**). These findings suggest that the shrew moles can locate targets by emitting USVs and hearing, strongly indicating that shrew moles are a novel lineage of echolocating mammals.

### Shrew moles produce biosonar clicks using their tongues

During exploring environments, the shrew moles consistently produce ultrasonic pulses of brief duration (∼0.8 ms) and wide bandwidth (∼69 kHz) (**Table S8**), which were significantly shorter and broader than those of frequency-modulated (FM) and constant-frequency (CF) echolocation signals of insectivorous bats (*P* < 0.01; **Fig. 4A, 4B**). Furthermore, the duration and bandwidth of shrew moles’ echolocation signals were largely comparable to click-like echolocation signals of cave-dwelling rousette bats, toothed whales, and tenrecs (*P* > 0.05; **Fig. 4B**; **Tables S8** and **S12**). These similar features to click-like echolocation calls suggest that the shrew moles produce biosonar clicks to sense the environments (*25*).

**Fig. 4.**
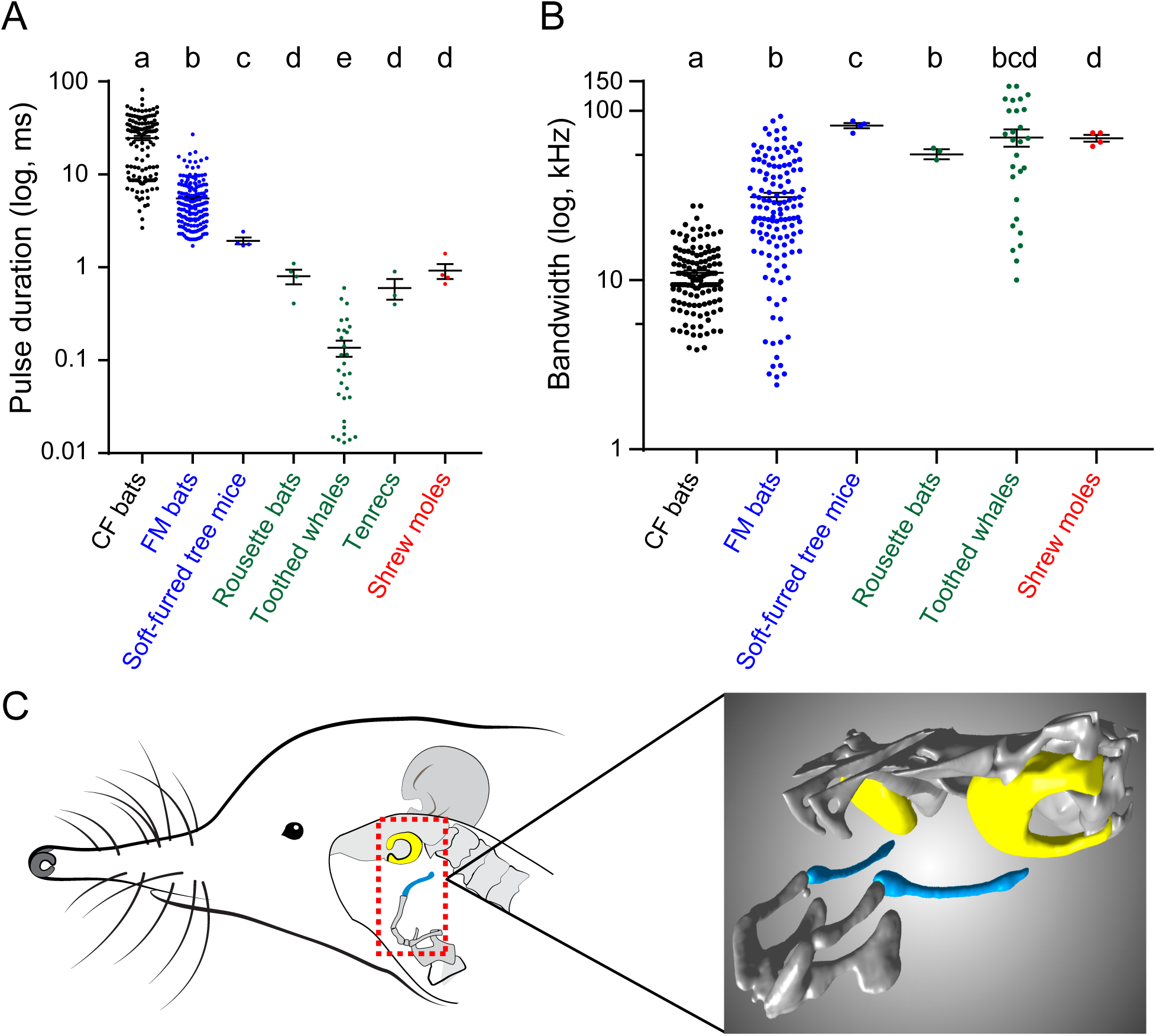
Signal features and morphology of vocal organ indicate that the shrew moles generate click-like signals for echolocation. Comparisons of the duration (**A**) and bandwidth (**B**) of different types of pulses in echolocating mammals. Each dot represents a value of one echolocating species. The raw data points are shown in **Tables S8** and **S12**. The bar denotes the mean value across all individuals, and the error bars represent SEM. *P* values are from Mann‒ Whitney U tests, different letters indicate significant differences, *p* < 0.05. (**C**) Stylohyal and tympanic bones were separated in *U. gracilis*. Left diagram showing the spatial positions of the stylohyal and tympanic bones in a gracile shrew mole. Blue and yellow colors in the dotted box highlight stylohyal and tympanic bones, respectively. Right diagram showing the stylohyal bone separated from the tympanic bone in the gracile shrew mole.

Next, we sought to uncover which organ was used to generate click-like echolocation signals in shrew moles. Unlike insectivorous bats and soft-furred tree mice producing tonal echolocation signals in the larynx (*6, 26*), the rousette bats and tenrecs generate echolocation clicks by the tongue (*5, 25*) and toothed whales by the phonic lips (*27*). Thus, no known echolocating mammals use the larynx to produce click-like echolocation signals. To further corroborate this in the shrew moles, we employed micro-computed tomography to scan the proximal articulation of the stylohyal bone with the tympanic bone, which is an anatomic feature of laryngeally echolocating mammals (*26*). The results demonstrated that the proximal end of the stylohyal bone did not contact the tympanic bone in shrew moles (*U.gracilis*, *U. nivatus*, and *U. investigator*) (**Fig. 4C**; **Fig. S11**), as in rousette bats (*26*).

To verify that the shrew moles generate click-like signals by tongue for echolocation, we conducted a serial of behavioral experiments while controlling the tongue movements. The rousette bat (*Rousettus leschenaultii*) and the soft-furred tree mouse (*T*. *daloushanensis*) were used as the positive and negative controls, respectively. First, we ligated the tongue of the rousette bat and found that these bats failed to produce any click-like signals while flying in the dark. When the ligation was removed, the rousette bats regained the ability to generate the click-like pulses (15.44 ± 0.45 per second), which was largely consistent with that before the treatment (15.12 ± 0.46) (*P* > 0.05; **Fig. 5A**). By contrast, when the tongue of the soft-furred mouse was ligated, the animal still produced echolocation pulses, albeit with a reduced pulse rate (**Fig. 5B**). These results strongly suggest that ligation was an effective method to assess whether the shrew moles use their tongues to generate click-like signals. Upon ligation of the shrew moles’ tongues, no click-like signals were produced. When the ligation was removed, the shrew moles again vocalized with a pulse rate similar to that prior to the treatment (*P* > 0.05; **Fig. 5C**).

**Fig. 5.**
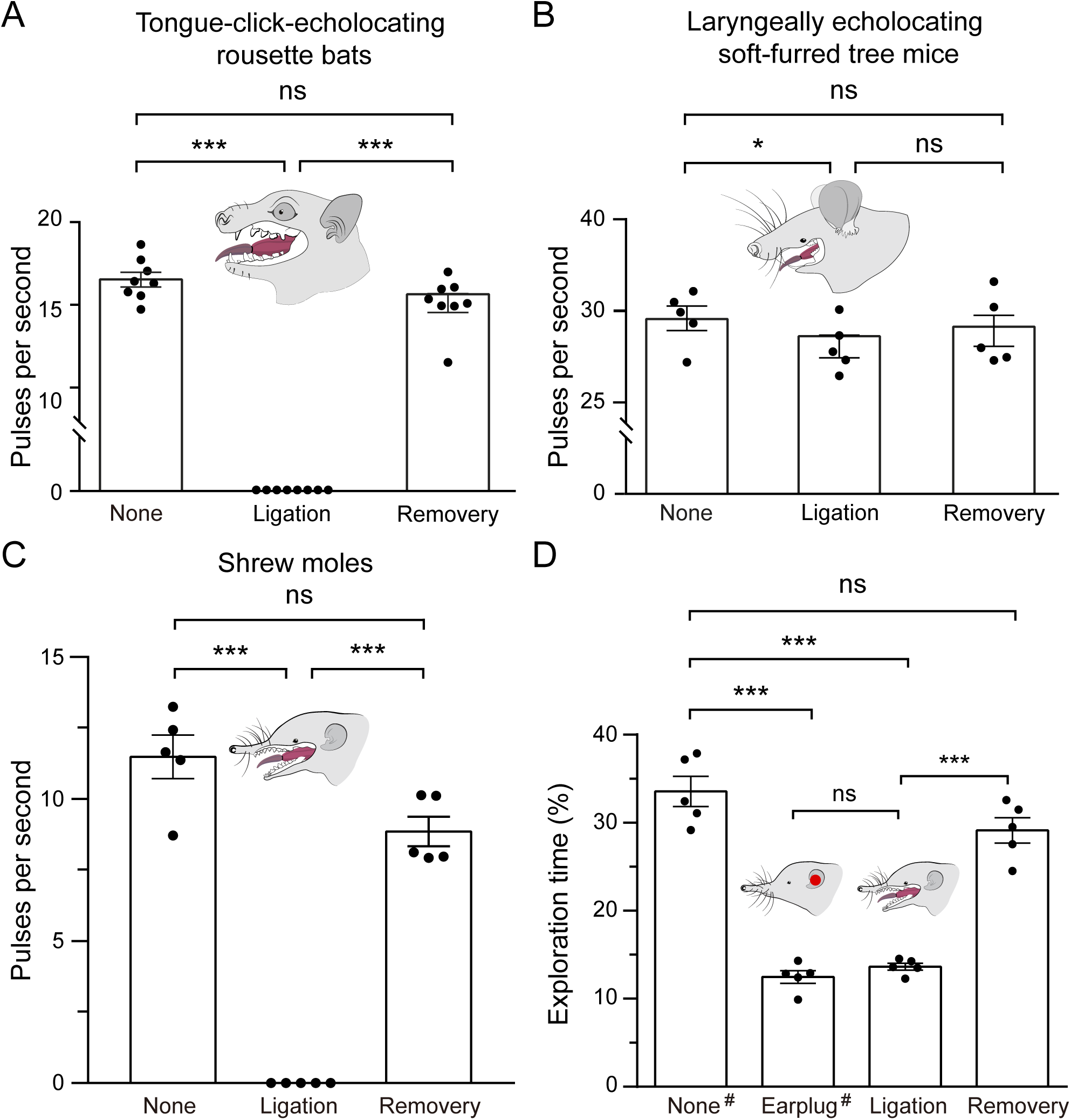
Shrew moles produce echolocation clicks with the tongue. Comparisons of pulse rates when echolocating mammals were tied a knot on the front of the tongue, (A) tongue-click-echolocating bats (*Rousettus leschenaultii*, N = 8), (**B**) laryngeally echolocating soft-furred mice (*Typhlomys daloushanensis*, N = 5), (**C**) gracile shrew moles (*U. gracilis*, N = 5). (**D**) Comparisons of the percentage of exploration time in the sector corresponding to the ultrasonic probe when the gracile shrew moles were exploring on the central disc under various experimental treatments, platform and earplug groups reference to figure 1, ligation group animals tongue was tied a knot, recovery group remove knot. Each dot represents the mean of the measurements of each individual. The bar denotes the mean value across all individuals, and the error bars represent SEM. *P* values are from two-tailed paired Student’s t tests and were corrected with the Bonferroni method. * *p* < 0.05; *** *p* < 0.001. ns, not significant.

To determine the implication of the tongue clicks of the shrew moles for echolocation, we performed behavioral experiments in the disc-platform apparatus. We found that when the tongue was ligated, the shrew moles failed to produce echolocation pulses on the over-platform sector, lost their preference for the sector, and were unable to detect the platform, as did when their ears were blocked (*P* > 0.05; **Fig. 5D**). Following the removal of the ligation, the shrew moles regained the capacity to produce click-like signals, and significantly increased the amount of time spent exploring the over-platform sector, succeeding in landing the platform (*P* < 0.001; **Fig. 5D**). These results further confirmed that the shrew moles produce click-like signals through the use of their tongues for echolocation, thus indicating a new lineage of echolocating mammals with tongue clicks.

## Discussion

In this study, we developed a machine learning model based on molecular convergences among echolocating mammals and trained it using a test dataset to screen for unknown echolocating species among 97 mammals with available high-quality genomic data. Three additional mammalian species, the gracile shrew mole, the nine-banded armadillo, and the solenodon, were identified as potential echolocators. We employed echolocation-related behavioral experiments (*6, 24*) to verify that not only the gracile shrew mole, but likely all shrew moles (Uropsilinae) have evolved echolocation.

It has been suggested that echolocation has independently evolved at least six times in mammals. Unfortunately, due to the unavailability of living animals, we were unable to perform the same behavioral experiments for the nine-banded armadillo and the solenodon. However, interestingly, the solenodon (family Solenodontidae) has been described as producing shrew-like sounds, which suggests that this species may be an echolocator (*28*), thus providing some support for our results. If it is true, combining with the echolocating shrews (family Soricidae) (*4*), there are three out of four families in the order Eulipotyphla that involve echolocating species, suggesting that order Eulipotyphla is the third major echolocating group of mammals in addition to the order Chiroptera and the infraorder Cetacea, and possess more complicated evolutionary trajectories of echolocation than the other two orders. If the nine-banded armadillo is confirmed to be an echolocator, the number of mammalian orders involving echolocating species will increase to six, accounting for approximately 20% of all living mammalian orders, thus empirically supporting Griffin’s hypothesis that echolocation is widespread among animals. As more mammalian genomes are available in the future, we will obtain more accurate number of how many animals possess echolocation abilities by using our developed pipeline.

Our behavioral experiments demonstrated that all four species of shrew moles examined use echolocation. As these species were selected from the main evolutionary clades of shrew moles, particularly *U. investigator* from the basal clade of shrew moles (**Fig. 2A**), it is reasonable to infer that echolocation evolved in the last common ancestors of shrew moles. This raises the question of the origin of echolocation in shrew moles. The shrew moles belong to the subfamily Uropsilinae, which is one of three subfamilies of the family Talpidae and is located at the basal position (*29*). It has been observed that the majority of species in the other two subfamilies Scalopinae and Talpinae inhabit fully fossorial, semi-fossorial, and semi-aquatic habitats, suggesting the scalopin and talpin moles may not use echolocation. This implies that echolocation either evolved in the last common ancestor of shrew moles or originated in the last common ancestor of Talpidae and was subsequently lost in the evolution of the scalopin and talpin moles. Two potential ideas may be proposed to address this question. If the fossils of the last common ancestor of the talpids could be discovered, one idea would be to examine the echolocation-related features, such as cochlear width, to determine whether they are the same in the echolocating shrew moles and whether there is evidence for the structural reduction in the scalopin and talpin moles, which would suggest a single gain of echolocation in the ancestor of the talpids and loss in the scalopin and talpin moles. The other idea would be to investigate molecular convergences in the genes related to echolocation between the branch of the last common ancestor of the talpids and known echolocating mammals. A significantly larger number of convergences in these genes would suggest that echolocation originated on the ancestral branch of the talpids (*7, 8*).

The use of machine learning methods to construct predictive models of biological traits based on molecular characteristics has been demonstrated to be effective, resulting in increased throughputs and reduced costs for subsequent research (*30, 31*). Our machine learning model allows us to identify new echolocating mammals on a large scale with only a limited number of convergent amino acids among known echolocating mammals. There are several reasons for selecting these sites as a training set. First, these convergent sites occurred not by chance and can effectively differentiate echolocating and nonecholocating mammalian species (**Fig. 1A**). Second, the experimental evidence has suggested that the adaptive convergent amino acids largely contributed to the functional convergence related to echolocation among echolocating mammals (*7, 8, 17, 32*). Nevertheless, we acknowledge that the caveat of the small number of convergent sites could potentially distort the predictive accuracy of our approach, which could be further enhanced as additional molecular elements or genes related to mammalian echolocation are incorporated to the training dataset.

A fundamental question in evolutionary genetics is the extent to which adaptively convergent phenotypes are caused by convergent changes at the molecular sequence level (*9, 33*). This raises the question of whether adaptative molecular convergence allows for the genetic prediction of similar phenotypes adapted to similar environmental challenges in independent lineages. Our approach and findings, which focus on mammalian echolocation, have made a beneficial exploration of these questions and have paved the way for determining the presence of other repeatable phenotypes in new mammalian lineages through the detection of adaptative molecular convergence.

## Acknowledgements

We gratefully acknowledge Chang-Zhe Pu, Ming-Jing Pu, Kang Luo and Jing Luo for their assistance in the field. We thank the Jinfoshan National Nature Reserve, the Qinling Mountain National Nature Reserve, the Yulong Snow Mountain Provincial Nature Reserve, and the Gaoligong Mountain National Nature Reserve for approving and facilitating our fieldwork.

## Author contributions

Conceptualization: PS

Methodology: PS, XLJ, YW, QL, QYH, RS

Investigation: QL, QYH, RS, SWH, MCL, LYS, YSM, QZ

Visualization: QL, QYH, RS, PC Funding acquisition: PS, XLJ, YW

Project administration: PS Supervision: PS, XLJ, YW

Writing – original draft: PS, QL, QYH, YW, RS

## Data and materials availability

All data are available in the main text or the supplementary materials.

## Supplementary Materials

### Materials and methods

#### Identification of molecular convergences

The high-quality genomic data was collected from the OrthoMam (v10c) database (*20*), GenBank (*34*) and previous studies (*6, 19, 35*), including 35 high-quality echolocating mammalian genomes from four orders: one rodential soft-furred tree mouse (*Typhlomys cinereus*), one tenrecid lesser hedgehog tenrec (*Echinops telfairi*), 14 insectivorous bats, and 19 toothed whales. To largely reduce the bias of the unequal number of closely related species against the subsequent evolutionary analyses, we chose a representative species for each non-echolocating mammalian genus in the dataset. Finally, a total of 93 mammalian species with high-quality genomic data were retained in our dataset (**Tables S1,** and **S2**) for the subsequent evolutionary analyses.

We obtained a total of 9,554 one-to-one orthologous protein-coding genes across 93 species using Inparanoid v4.1 (*36*). PRANK (*37*) was used to align each of these protein-coding genes. To identify convergent amino acid substitutions for echolocation, we inferred ancestral amino acid sequences for each interior node of the phylogeny of these species (**Fig. S1**) using the maximum likelihood method built in PAML (*18*). To ensure the reliability of the results, only ancestral amino acids with posterior probabilities ≥0.95 were considered. On the one hand, given that echolocation originated in the last common ancestor (LCA) of bats and the LCA of toothed whales (*7, 8*), we identified convergent amino acid substitutions among the soft-furred tree mouse, the lesser hedgehog tenrec, the LCA of bats, and the LCA of toothed whales. On the other hand, echolocation displays remarkable diversity in ultrasonic characteristics, behavioral modes, anatomic features of auditory organs, and so on across different species of echolocating bats and toothed whales (*25, 38*). To largely include the amino acid substitutions related to echolocation, we also identified convergent sites among the soft-furred tree mouse, the lesser hedgehog tenrec, any of 14 echolocating bats, and any of 19 toothed whales. By combining the two datasets above, we identified a total of 497 convergent amino acid substitutions that are putatively related to mammalian echolocation.

To further ensure the relevance of these molecular convergences to echolocation, we refined these candidate convergent sites according to the flowing criteria. A convergent amino acid substitution was removed if it (i) occurred in the closest nonecholocating species or on the ancestral branches of non-echolocators; (ii) occurred in less than 5 echolocators; (iii) occurred in any of 58 non-nocturnal species or known non-echolocators; (iv) occurred randomly among echolocators, which was estimated through a statistical test by comparing with a null hypothesis derived from neutral evolution (*39*). After filtering, a total of 14 convergent amino acid substitutions in 14 protein-coding genes were retained (**Table S3**), which were used as a training dataset for subsequent machine learning.

#### Machine learning prediction for new mammalian echolocators

Genotypes including adaptive genetic changes have long been used to predict complex traits (*40*). In this study, we developed a machine learning model to predict new mammalian echolocators using the convergent amino acid substitutions closely related to echolocation.

##### Step1: Amino acid representation

Prior to constructing the machine learning framework, we assessed the proper representation of the identified convergent amino acid substitutions (**Table S3**) because they were required to be converted to numerical data for modeling. We tried eight different encoding methods according to amino acid properties, including hydrophobicity, normalized Van der Waals volume, polarity, polarizibility, charge, secondary structures, solvent accessibility and one hot (*41*), and evaluated the performance of these methods based on their corresponding principal component analyses (PCA) (**Fig. S2**). The PCA results showed that the encoding method of the one hot clearly separated the mammals into two groups of echolocators and non-echolocators. Thus, the one hot was selected as an ideal encoding method for further analyses.

##### Step2: Bayesian network to integrate features of convergent amino acid substitutions

To properly model and accurately predict whether a mammal species is an echolocator, we evaluated 5 different machine learning algorithms, including Naïve Bayes, Logistic Regression, Decision Tree, SVM, and Random Forest using k-fold cross validation and being trained from features of 14 convergent amino acid substitutions across 85 representative echolocating and non-echolocating mammalian genera (**Fig. 1B**). We finally selected Naïve Bayes model for prediction cause its superior performance in accuracy, precision, recall, F1 score and Roc_Auc (**Fig. S3**).

In detail, the prediction of echolocation for each mammal species (labelled Y = 1 if it is an echolocator, and 0 otherwise) is based on the use of the Bayes rule to calculate the posterior odds of echolocation given the presence of genomic features *x*_1_, *x*_2_, …, *x*_*n*_ (n=14) as follows:

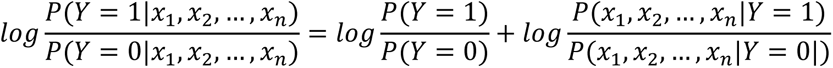

where *P*(*Y* = 1|*x*_1_, *x*_2_, …, *x*_*n*_) is the probability that the mammal is an echolocator given these features. P(Y=1)/P(Y=0) is the prior odds. *P*(*x*_1_, *x*_2_, …, *x*_*n*_|*Y* = 1)/*P*(*x*_1_, *x*_2_, …, *x*_*n*_|*Y* = 0) is the likeli-hood ratio for the combined features. Specifically, we generated a type frequency table for each of the 14 convergent amino acid substitutions that emerged in echolocating and nonecholocating groups, respectively (**Fig. S4**). Then, we calculated the likelihood ratio of the type frequency for each amino acid along the concatenated convergent amino acids. The likelihood ratio of the *i*-th amino acid is the number of *i*-th amino acid xi relative to the total number of amino acids in the echolocating group (Y=1) divided by the number of the *i*-th amino acid *x*_i_ relative to the total number of amino acids in the nonecholocating group (Y=0).

##### Step 3: Echolocation score

When we assume that the genomic features of convergent amino acid substitutions are conditionally independent given echolocation, the likelihood ratio of the combined features (Echolocation Score) is equal to the product of the likelihood ratios for individual features. Briefly, we provide an estimation towards every convergent amino acid substitution based on its amino acid type and sum them up.

#### Resampling simulation for the confidence intervals of echolocation scores

To further improve the prediction accuracy, we randomly picked a site from 14 convergent amino acid substitutions among 28 echolocating mammalian species in the training dataset to generate a simulated dataset and estimated its ES. Such progress was repeated 10000 times to obtain a null distribution of ES values for echolocating mammalian species. Then, the confidence intervals of the echolocation scores can be estimated based on the null distribution (**Fig. S5**).

#### Using the echolocation score to predict new echolocators

The genomic data used for predicting new mammalian echolocators was collected from the Zoonomia Project (*3*) and the OrthoMam (v10c) database (*1*). To make sure the reliability of the results, we only considered the relatively high-quality genomic data with single copy > 60%. After filtering, we obtained the genomic data of 74 species from the Zoonomia Project and of 23 nocturnal species in the OrthoMam database due to lacking sufficient evidence to determine whether they are capable of echolocation (**Tables S5,** and **S6**). This prediction dataset included 12 known echolocating mammals that were used as positive controls to assess the accuracy of the predictions. To calculate the ES values of these 97 mammalian species, we first retrieved the protein-coding sequences of 14 genes containing the convergent amino acids from the genomic data using *genewise* v2.4.1 (*42*) and further obtained the amino acids corresponding to the 14 convergent amino acid substitutions (**Table S13**). Then, the ES value of each species in the prediction dataset was estimated based on the above pipeline. If the sites at the position corresponding to the convergent amino acid substitutions were unavailable in the species with ES > 0, we used PCR to amplify the sequences to obtain these sites and re-estimated the ES of the species. In the list, only *U. gracilis* with ES > 0 had four such sites unavailable in the genomic data and thus we designed the following primers to obtain these sites in the species.

COL11A1-F AGCAATCAAGATCTCTCC

COL11A1-R TATCATTCTGATGATACA

EFL1-F GCTGCCTGGAAGACCTGACG

EFL1-R TCCACCRCGTCRCGCCACTG

OTOF-F GCCTATTTAGCAGAACTCAGTCC

OTOF-R TAGGATCTTCTTGACCAGGTAGC

ZNF316-F GCCCTGGTTCAATGAGCTCCAAATGAG

ZNF316-R GCTGCCCAGCTGGAGAGGGCAGAGG

Based on the completed sequences, we obtained the final ES results for 97 mammals (**Table S7**).

#### Shrew moles for behavioral experiments

Four species of shrew moles (*Uropsilus*) were captured using live-trap, including *U. gracilis* (N = 16) from Mountain Jinfo, Chongqing City; *U. Soricipes* (N = 14) from Mountain Qinling, Shanxi Province; *U. nivatus* (N = 11) from Mountain Yulong, Yunnan Province; and *U. investigator* (N = 7) from Mountain Gaoligong, Yunnan Province. These animals were individually housed in a plastic box (46 × 30 × 16 cm) in a condition-controlled room (18–25 °C, 50–70% humidity) at Kunming Institute of Zoology, Chinese Academy of Sciences. Wood shavings covered the bottom of the plastic box and a small plastic box (12.7×7.6×7.6 cm) was put on one side as a nest. Shrew moles were fed mealworms and fruits. Water was available ad libitum. As shrew moles are nocturnal, behavioral experiments were conducted at night. All related animal experiments were approved by the Ethics Committee of the Kunming Institute of Zoology, Chinese Academy of Sciences.

#### Recording and analyzing vocalizations of shrew moles

We used the Avisoft Bioacoustics CM16/CMPA microphone and the Avisoft UltraSoundGate 416H audio device (Avisoft Bioacoustics, Berlin) to perform audio recording at a sampling rate of 250 kHz (16 bit) and frequency response of 2 ∼ 200 kHz. The recorded acoustic data were analyzed using the Avisoft-SASLab Pro software v5.2 (Avisoft Bioacoustics, Berlin) in a Hamming window with a 512-point fast Fourier transform, a time-window overlap of 75 % and threshold 50. We used the Automatic Parameter Measurements tool (two thresholds, thresholds -10 dB, start/end -10 dB, hold time 2 ms) to obtain the acoustic parameters, including peak frequency, maximum frequency (*f*_max_), minimum frequency (*f*_min_), frequency bandwidth, pulse duration, pulse period, and pulse interval. The numbers of ultrasonic pulses were counted with the Pulse Train Analysis tool (time constant 1 ms, threshold -15 dB). The data of ultrasonic pulses from different shrew mole species were shown in **Table S8**.

#### Behavioral experiments for echolocation in shrew moles

The disc-platform apparatus (*4*) was used to determine whether shrew moles echolocate, which mainly consists of two components: disc-small disc circle and disc- platform (**Fig. 3A**). The small-disc circle is a 40 cm diameter loop with equally arranged eight 3.6 cm diameter discs, which was used as a control. The center disc was equally divided into eight sectors and each sector corresponded to a reward box containing freshly killed mealworms. In principle, if an animal echolocates, when placed on the central disc, it should be able to use sonic signals to find the platform located some distance below the disc and jump down onto the platform without the use of tactile, olfactory, or visual information. To eliminate odors, the center disc and platform were covered with filter paper that was replaced before every trial. The ramp and reward box were rinsed with distilled water, then with 100 % ethyl alcohol, and finally with distilled water before every trial. The microphone was wiped with cotton dipped with 100% ethyl alcohol and distilled water.

We first placed the small-disc ring 7 cm vertically below the center disc to conduct control tests. And then the small-disc ring was replaced with a platform that was also placed 7 cm below one of the eight sectors. The animals on the center disc cannot lean over the edge of the disc to touch the small-disc ring or the platform with the nose. According to the protocol previously described (He et al. 2021), we performed the behavioral experiments under various treatments in the following order: (a) the animals were placed on the disc-small disc ring apparatus without any treatment; (b) the animals were placed on the disc-platform apparatus without any treatment; (c) the animals whose ears were plugged with expandable foam were placed in the disc-platform apparatus; (d) the animals with shams (plastic tubes) in their ears were placed on the disc-platform apparatus; and (e) the animals with the expandable foam or plastic tubes being removed from their ears were placed in the disc-platform apparatus. All of these behavioral experiments were performed in a quiet, enclosed darkroom, thus precluding any impact of vision on animals’ behaviors.

Each of the four shrew mole species *U. gracilis* (N = 16), *U. Soricipes* (N = 14), *U. nivatus* (N = 11), and *U. investigator* (N = 7) was tested for echolocation individually. For each of the behavioral experiments above, we required four trials for each individual to ensure the reliability and repeatability of the results. The emitted ultrasonic vocalizations were recorded using the microphone fixed in one of eight small discs and the platform, respectively. The position of the microphone on the disc-small disc ring apparatus was determined randomly and the same position was set in the disc-platform apparatus for each trial. The infrared camera installed directly above the apparatus was used to perform video recordings during behavioral experiments. The audio and video recordings were analyzed synchronously to examine how the animals emitted ultrasonic calls during their movements in the apparatus. We analyzed video recordings with the VisuTrack software (Shanghai XinRuan Information Technology Co., Ltd.) that automatically documented the time when the animals entered and left the monitored sector, as well as the total exploration time on the center disc. The ultrasonic emissions in the monitored sector and in its adjacent sectors were counted from the audio recordings using the software and methods above. We calculated the percentages of exploration time and pulse rate when the animals were exploring in the monitored sector. The data of the behavioral experiments above were shown in **Tables S9, S10, S11**.

#### Micro-computed tomography (MCT) imaging

The images of the thyroid cartilages were taken with a stereomicroscope (SMZ18, Nikon). MCT images of the spatial relationship between the stylohyal and tympanic bones were obtained for three shrew mole species (*U. gracilis*, *U. investigator*, and *U. Soricipes*). The MCT scanner (Bruker Skyscan 1176) was used to collect images with a resolution of 9 μm at 65 kV and 310 μA. We used three-dimensional analysis software (CT-VOX) to reconstruct and visualize the data.

#### Verifying the tongue as a vocal organ for echolocation in shrew moles

To verify the tongue as a vocal organ for echolocation, we recorded ultrasonic vocalizations of shrew moles while controlling the movements of the tongue. After the animals were completely anesthetized with isoflurane using the anesthesia machine (Reward RWD500), we tied a knot on the middle of the tongue with the surgical suture (0.2 mm). The ligation was removed after behavioral experiments. We used the same methods to treat the rousette bat (*R*. *leschenaultii*) and the soft-furred tree mouse (*T*. *daloushanensis*). Because the rousette bat emitted echolocation calls with the tongue (*11*) and the soft-furred tree mouse with the larynx (*4*), these two kinds of animals were used as the positive and negative controls, respectively. We recorded ultrasonic vocalizations of shrew moles (*U. gracilis*) and soft-furred tree mice in cages (25×25×25 cm), and the rousette bats flying in an empty room (3×3×3 m). Before, during, and after the tongue being ligated, we recorded ultrasonic vocalizations of each animal for 10 minutes, respectively. The experiments were repeated three times for each individual of each species to ensure that the results were reliable and repeatable. Finally, we placed shrew moles with ligated tongue in the disc-platform apparatus to determine whether shrew moles echolocate with the tongue.

**Fig. S1.**
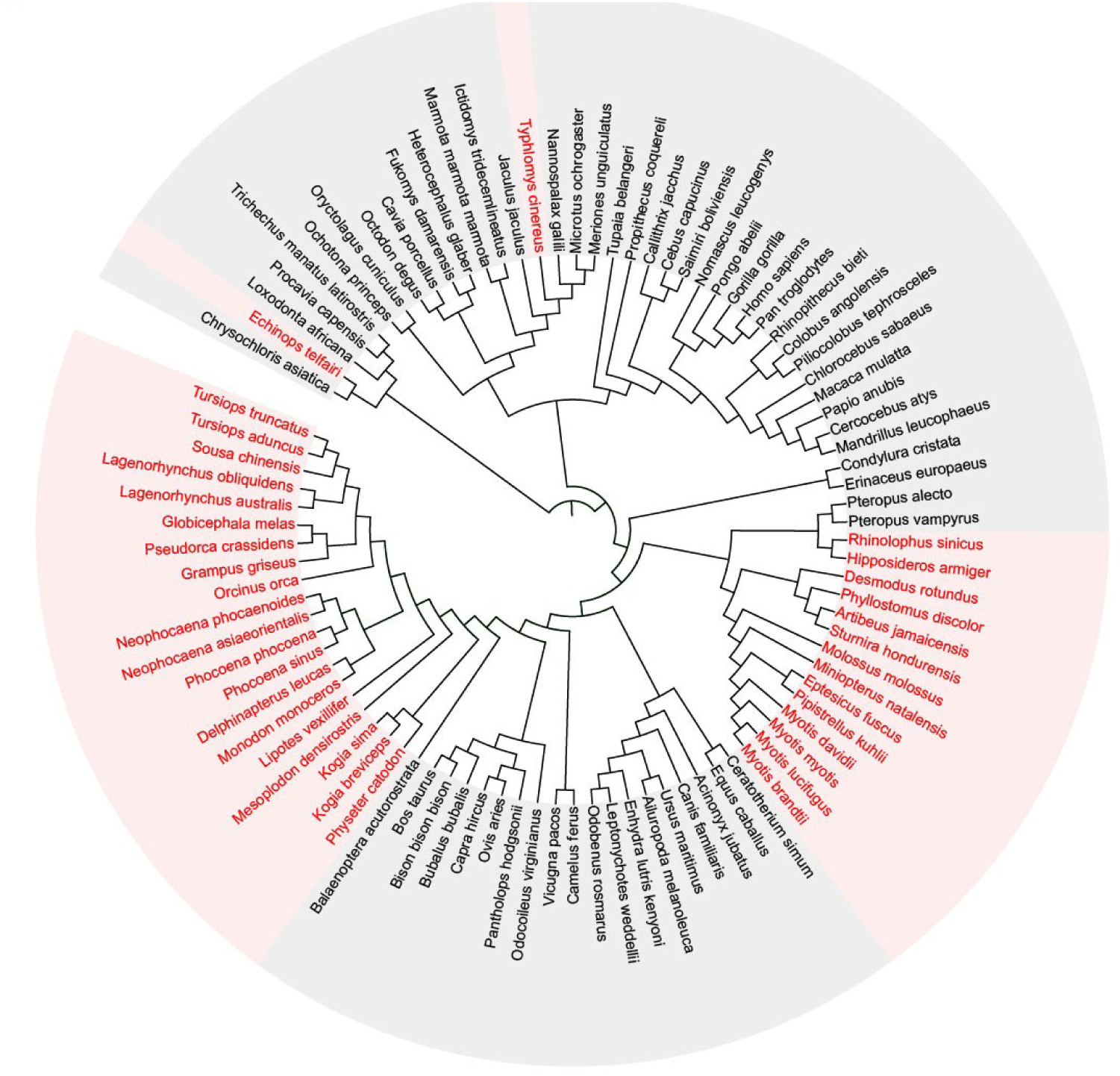
The phylogeny of 93 mammalian species was used to identify molecular convergence.

**Fig. S2.**
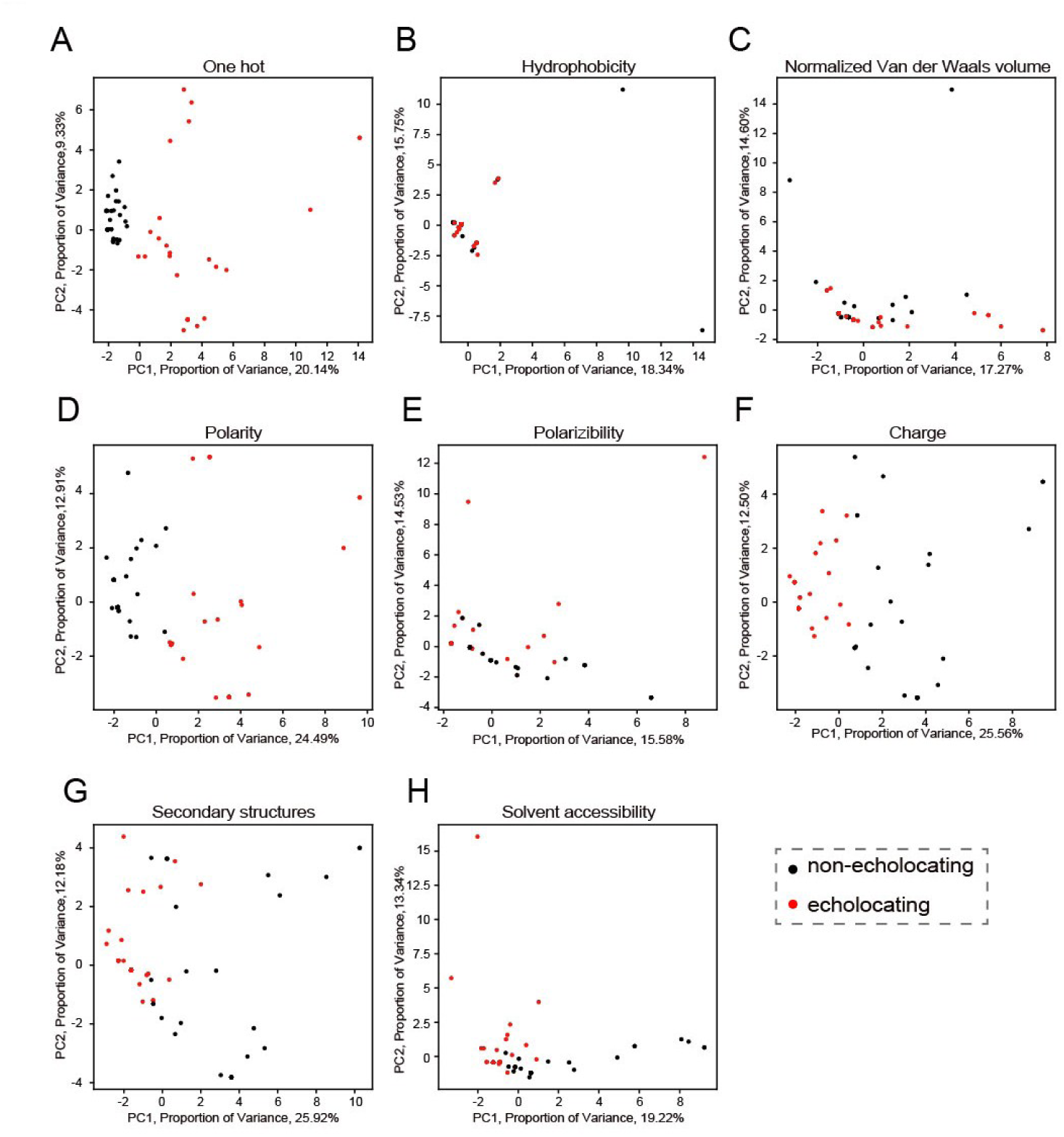
PCA visualization with different encode criteria. (A) PCA visualization with basic one hot encoding. (B) PCA visualization with hydrophobicity encoding. (C) PCA visualization with normalized Van der Waals volume encoding. (D) PCA visualization with polarity encoding. (E) PCA visualization with polarizability encoding. (F) PCA visualization with charge encoding. (G) PCA visualization with secondary structures encoding. (H) PCA visualization with solvent accessibility encoding.

**Fig. S3.**
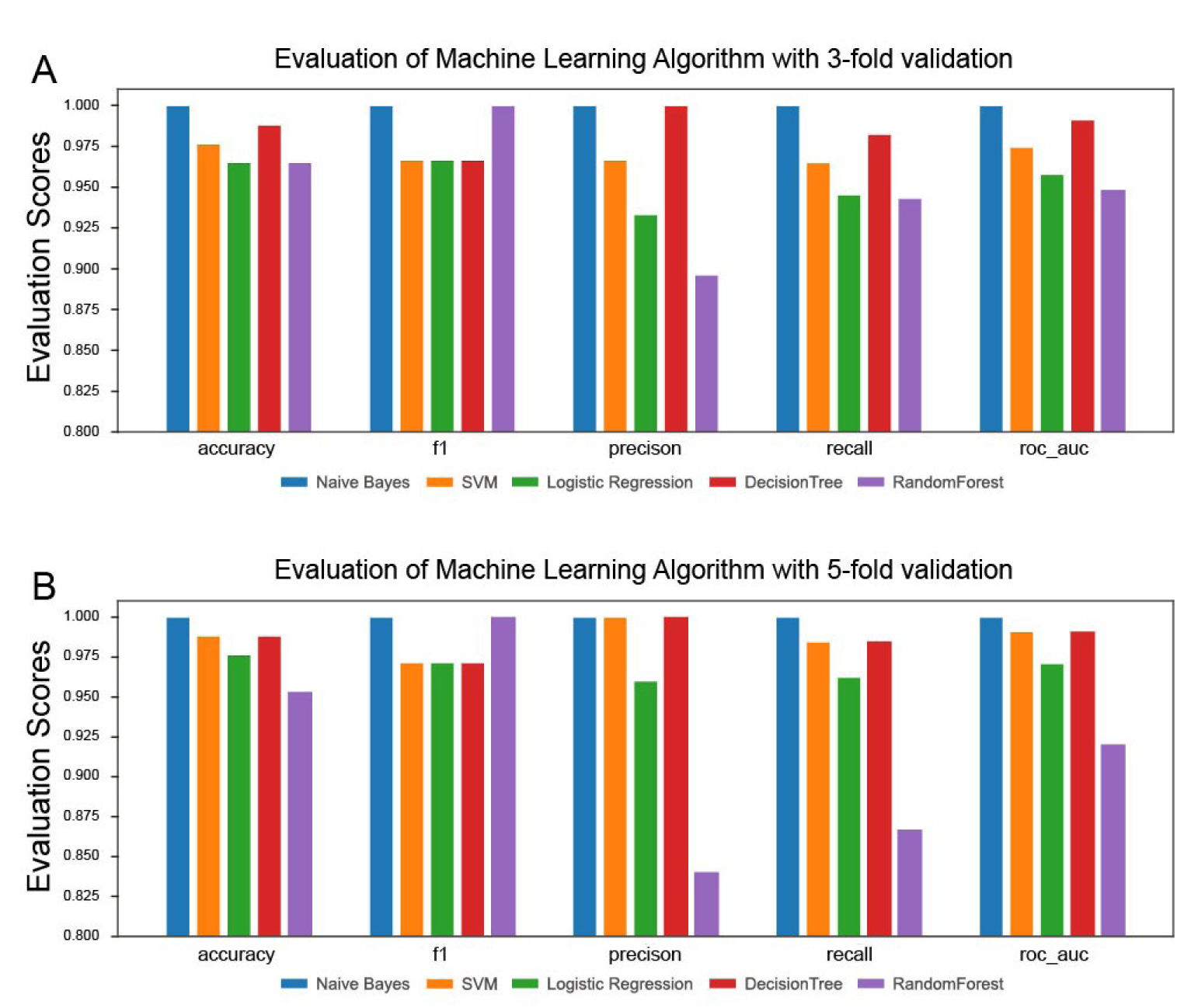
Evaluation of machine learning algorithm. (A) Comparison result among 5 machine learning frameworks with 3-fold cross validation. (B) Comparison result among 5 machine learning frameworks with 5-fold cross validation

**Fig. S4.**
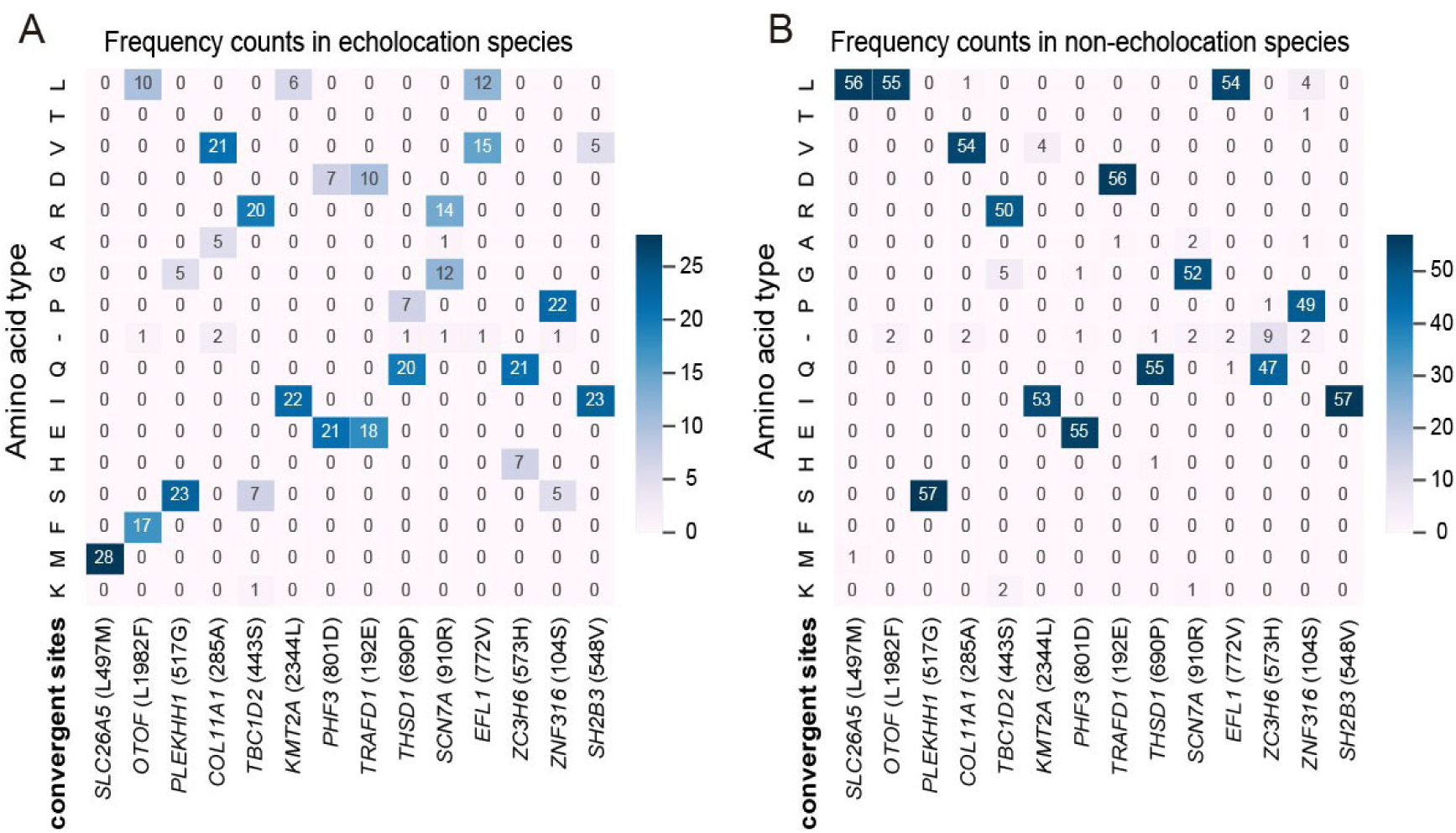
Amino acid frequency counts tables. (A) Amino acid frequency counts in echolocating species at 14 convergent sites. (B) Amino acid frequency counts in non-echolocating species at 14 convergent sites.

**Fig. S5.**
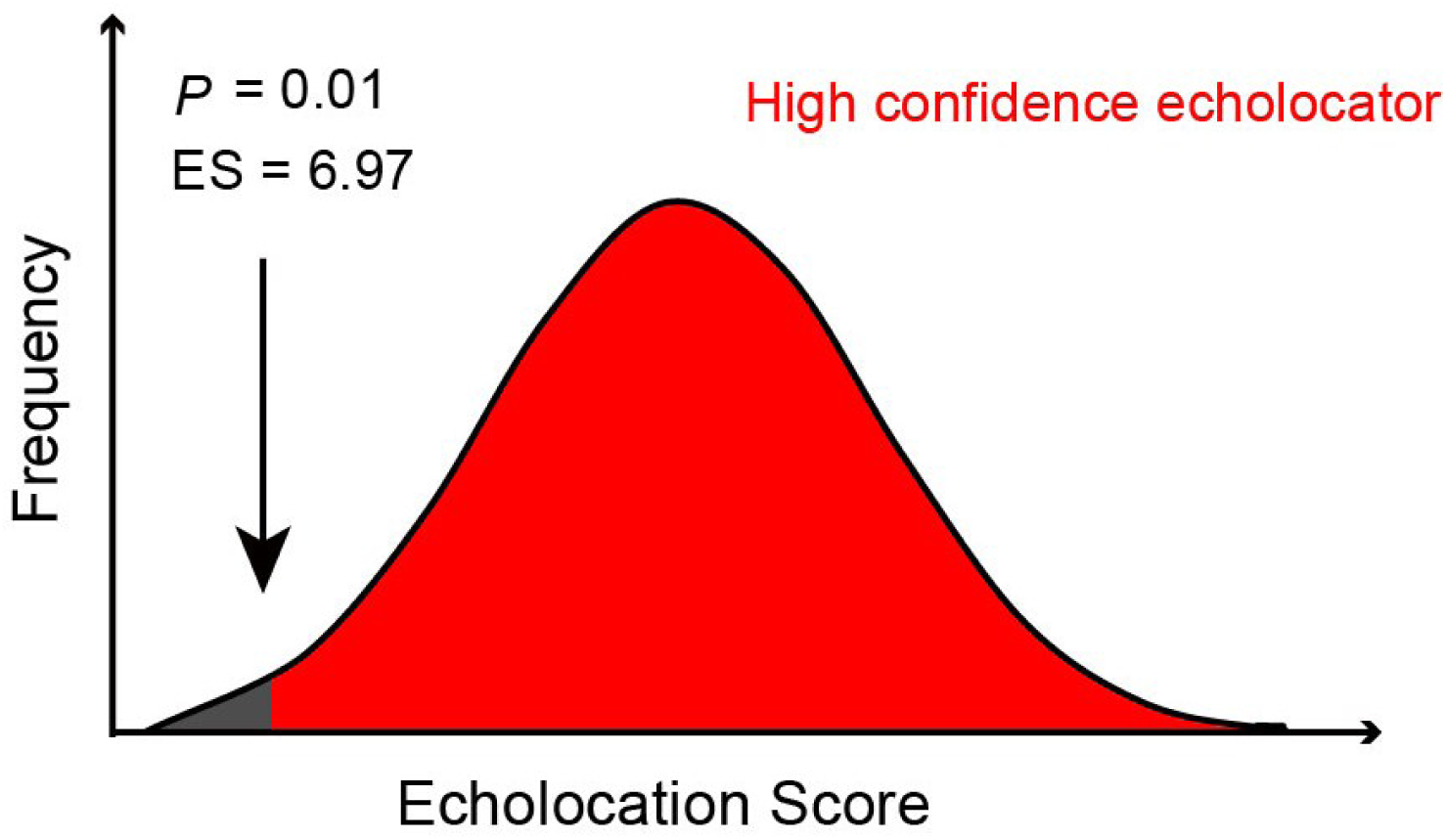
Null distribution of ES values from the simulated datasets.

**Fig. S6.**
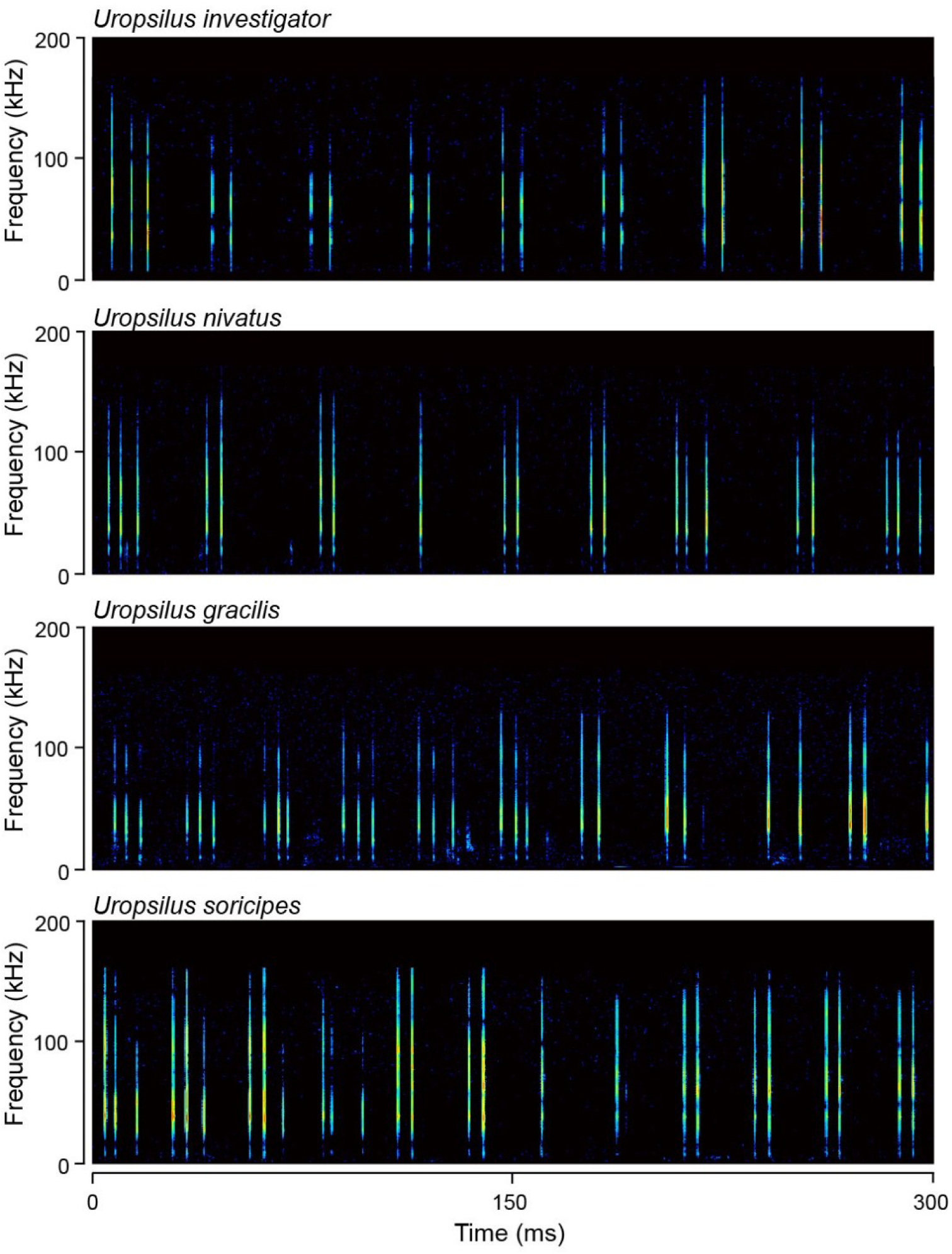
Records of natural sequence of pulses emitted by four shrew mole species.

**Fig. S7.**
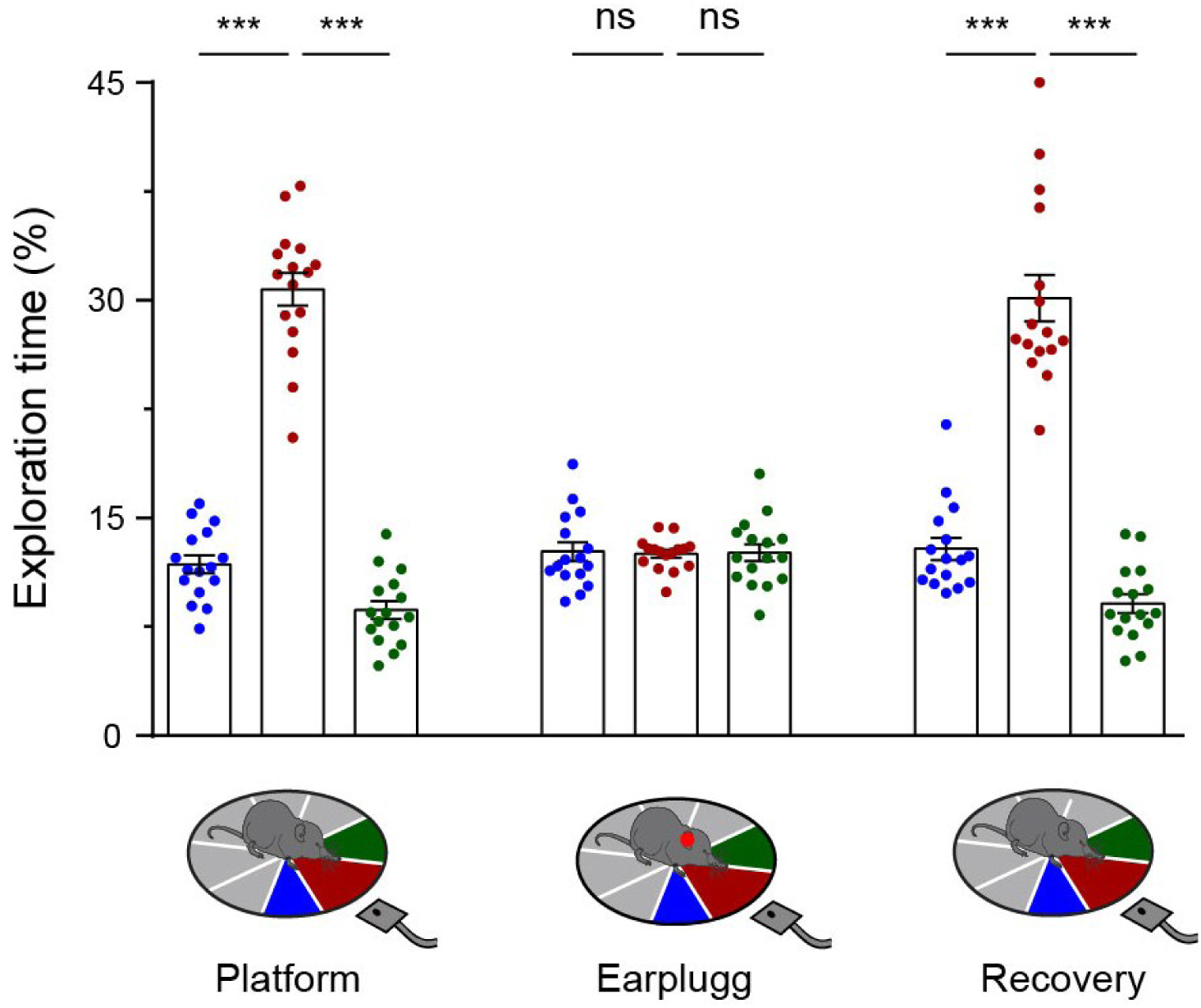
Comparisons of the exploration time and pulse rate within the platform, earplug and recovery groups. The percentage of exploration time was significantly larger when the gracile shrew moles (*U. gracilis*) explored in the monitored sector (red) than in its adjacent sectors (blue and green) in the platform and recovery groups. There was no significant difference in the percentage of exploration time when the gracile shrew moles explored in the monitored sector than in its adjacent sectors with earplugs. Each dot represents the mean of the measurements of each individual (n = 16 biological independent replicates). The bar denotes the mean value across eight individuals, and the error bars represent SEM. values. The *p* values are from two-tailed paired Student’s t tests. *** *p* < 0.001. ns means not significant.

**Fig. S8.**
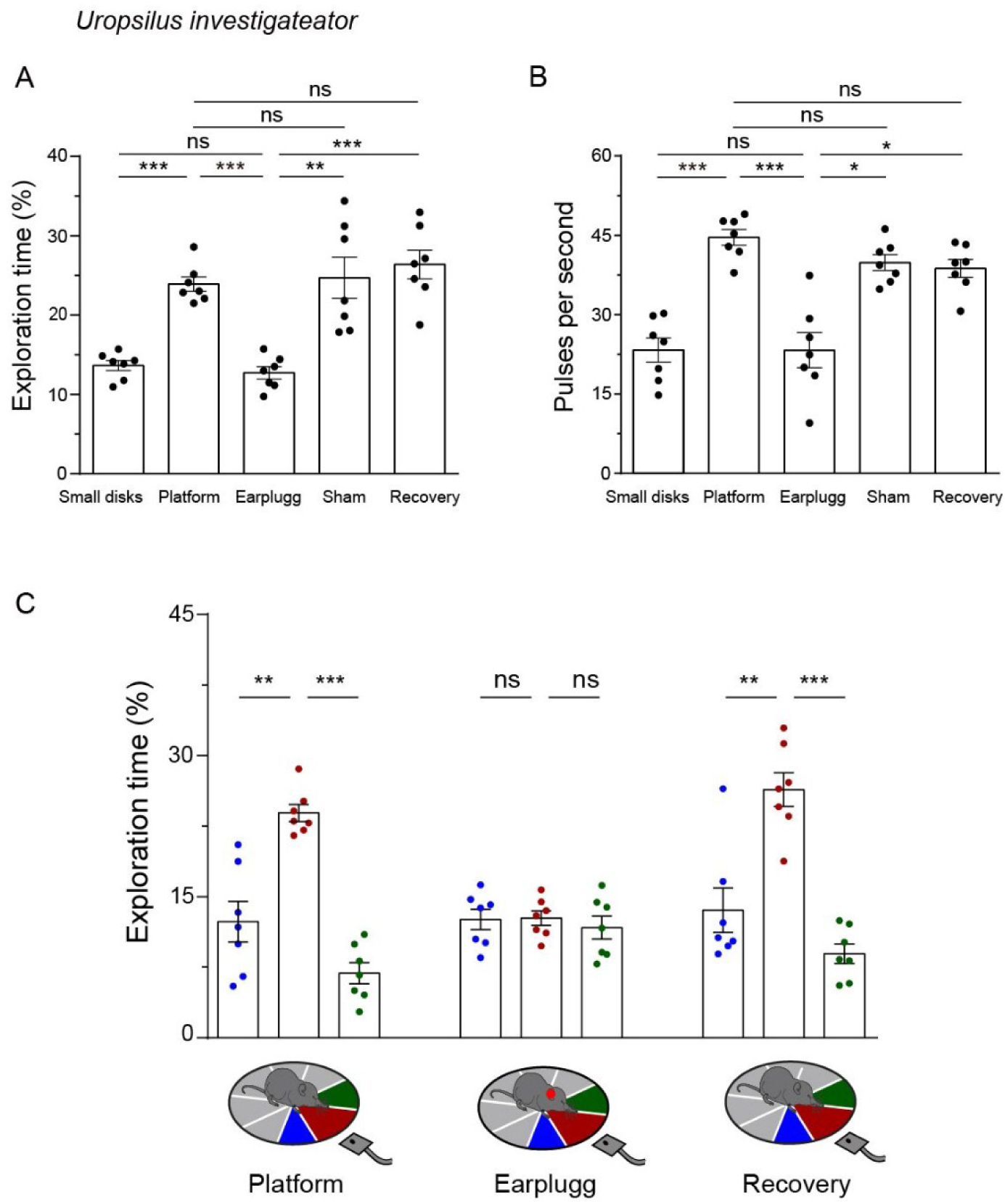
Behavioral experiments for testing echolocation in *U.* investigateator. Comparisons of the percentage of exploration time (A) and pulse rates (B) in the sector corresponding to the ultrasonic probe under various experimental treatments when *U. investigator* explored the central disc. (C) The percentage of exploration time when *U. investigator* exploring in the monitored sector is also compared with its adjacent sectors within platform, earplug and recovery groups. The inset shows the monitored sector (red) and its adjacent (blue and green) sectors. Each dot represents the mean of the measurements of each individual (N = 7). The raw data points are shown in **Table S9**. The bar denotes the mean value across all individuals, and the error bars represent SEM. *P* values are from two-tailed paired Student’s t tests and were corrected with the Bonferroni method. * *p* < 0.05; ** *p* < 0.01; *** *p* < 0.001. ns, not significant.

**Fig. S9.**
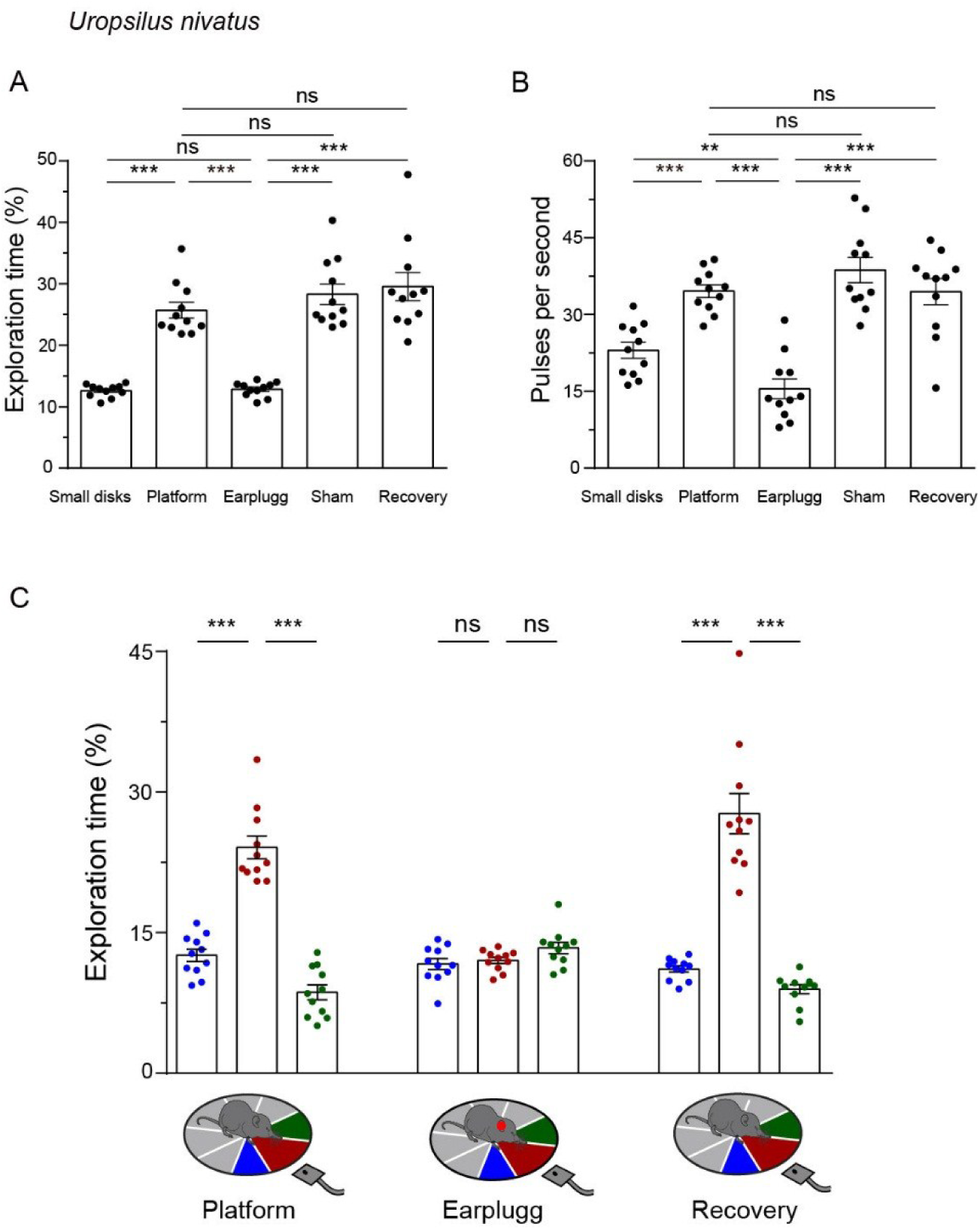
Behavioral experiments for testing echolocation in *U. nivatus*. Comparisons of the percentage of exploration time (A) and pulse rates (B) in the sector corresponding to the ultrasonic probe under various experimental treatments when *U. nivatus* explored the central disc. (C) The percentage of exploration time when *U. nivatus* explored the monitored sector is also compared with its adjacent sectors within the platform, earplug and recovery groups. The inset shows the monitored sector (red) and its adjacent (blue and green) sectors. Each dot represents the mean of the measurements of each individual (N = 11). The raw data points are shown in **Table S10**. The bar denotes the mean value across all individuals, and the error bars represent SEM. *P* values are from two-tailed paired Student’s t tests and were corrected with the Bonferroni method. * *p* < 0.05; ** *p* < 0.01; *** *p* < 0.001. ns, not significant.

**Fig. S10.**
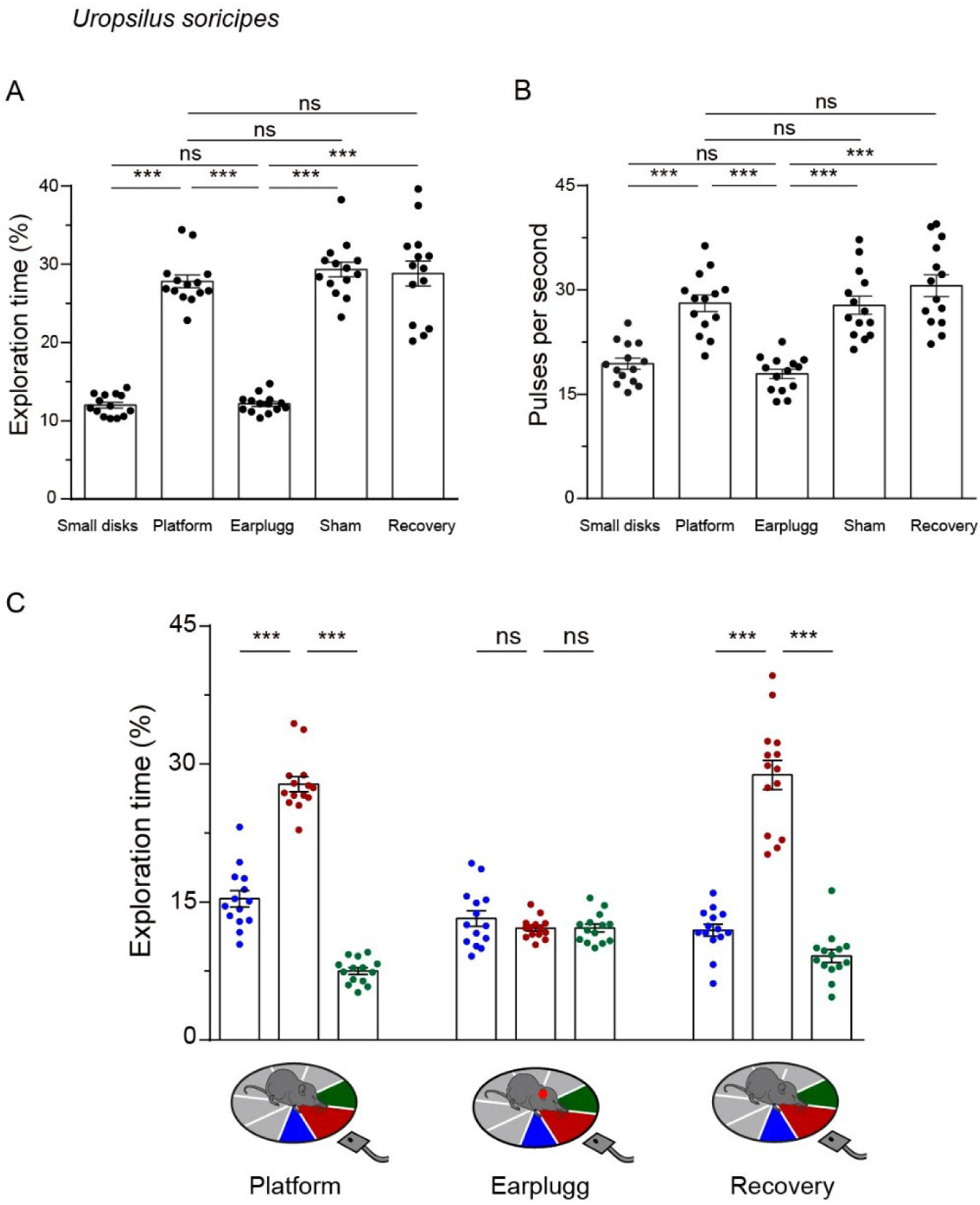
Behavioral experiments for testing echolocation in *U. soricipes*. Comparisons of the percentage of exploration time (A) and pulse rates (B) in the sector corresponding to the ultrasonic probe under various experimental treatments when *U. soricipes* explored the central disc. (C) The percentage of exploration time when *U. Soricipes* explored in the monitored sector is also compared with its adjacent sectors within the platform, earplug and recovery groups. The inset shows the monitored sector (red) and its adjacent (blue and green) sectors. Each dot represents the mean of the measurements of each individual (N = 14). The raw data points are shown in **Table S11**. The bar denotes the mean value across all individuals, and the error bars represent SEM. *P* values are from two-tailed paired Student’s t tests and were corrected with the Bonferroni method. * *p* < 0.05; ** *p* < 0.01; *** *p* < 0.001. ns, not significant.

**Fig. S11.**
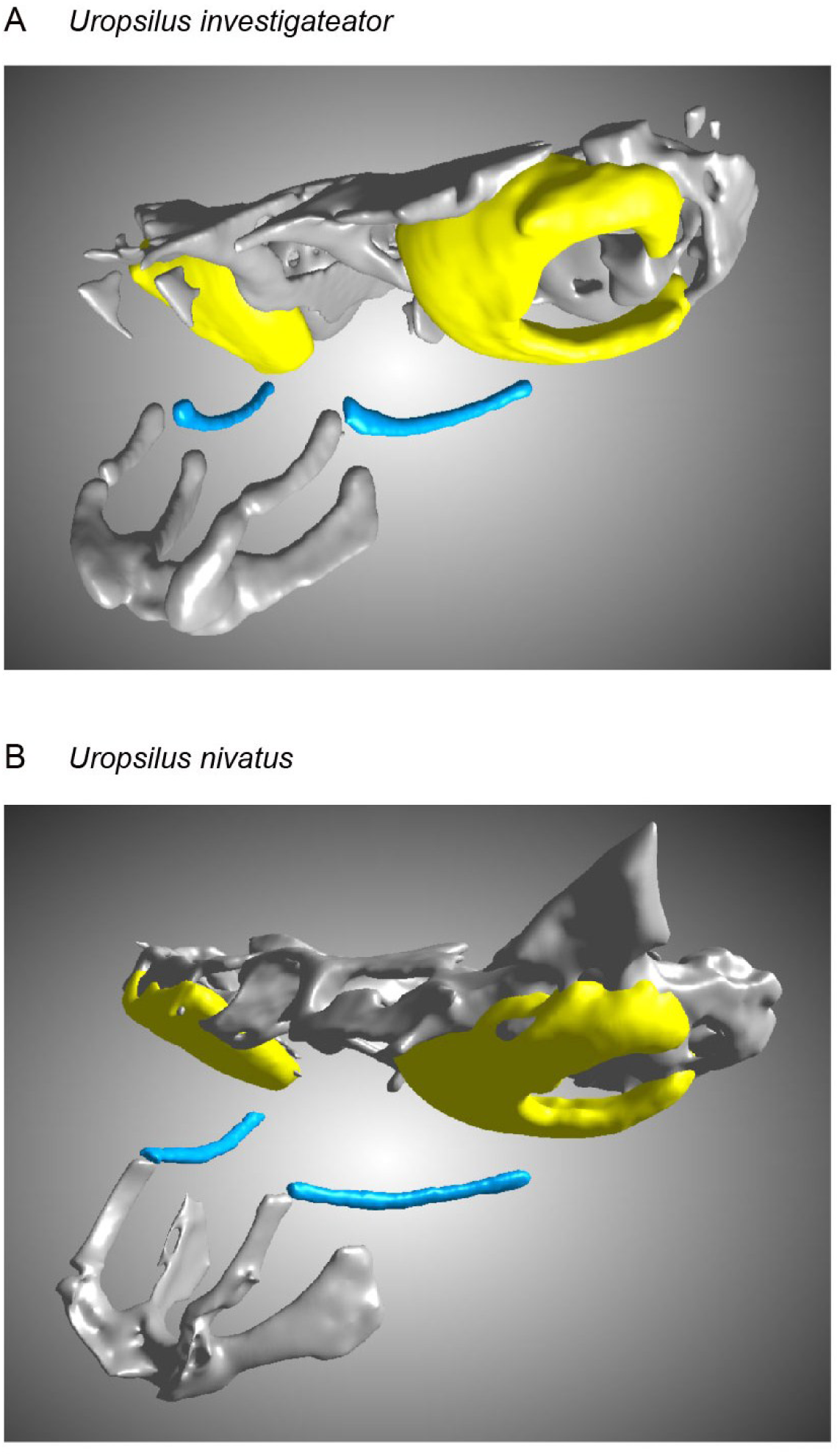
Stylohyal and tympanic bones in *U.* investigateator (A) and *U. nivatus* (B).

**Table S1.** 40 echolocating mammalian species and control species used to identify molecular convergence.

| Species | Common Name | Order | Family | References |
| --- | --- | --- | --- | --- |
| *Chrysochloris asiatica* | Cape golden mole | Afrosoricida | Chrysochloridae | 20 |
| *Echinops telfairi* | Small Madagascar hedgehog | Afrosoricida | Tenrecidae | 20 |
| *Balaenoptera acutorostrata* | Minke whale | Artiodactyla | Balaenopteridae | 20 |
| *Globicephala melas* | Long-finned pilot whale | Artiodactyla | Delphinidae | 35 |
| *Grampus griseus* | Risso's dolphin | Artiodactyla | Delphinidae | 35 |
| *Lagenorhynchus australis* | Peale's dolphin | Artiodactyla | Delphinidae | 35 |
| *Lagenorhynchus obliquidens* | Pacific white-sided dolphin | Artiodactyla | Delphinidae | 35 |
| *Orcinus orca* | Killer whale | Artiodactyla | Delphinidae | 20 |
| *Pseudorca crassidens* | False killer whale | Artiodactyla | Delphinidae | 35 |
| *Sousa chinensis* | Indo-pacific humpbacked dolphin | Artiodactyla | Delphinidae | 35 |
| *Tursiops aduncus* | Common bottlenose dolphin | Artiodactyla | Delphinidae | 20 |
| *Lipotes vexillifer* | Yangtze River dolphin | Artiodactyla | Lipotidae | 20 |
| *Delphinapterus leucas* | Beluga whale | Artiodactyla | Monodontidae | 20 |
| *Monodon monoceros* | Narwhal | Artiodactyla | Monodontidae | 35 |
| *Neophocaena asiaeorientalis* | Narrow-ridged finless porpoise | Artiodactyla | Phocoenidae | 35 |
| *Neophocaena phocaenoides* | Indo-Pacific finless porpoise | Artiodactyla | Phocoenidae | 35 |
| *Phocoena phocoena* | Harbor porpoise | Artiodactyla | Phocoenidae | 35 |
| *Phocoena sinus* | Vaquita | Artiodactyla | Phocoenidae | 35 |
| *Kogia breviceps* | Pygmy sperm whale | Artiodactyla | Physeteridae | 35 |
| *Kogia sima* | Dwarf sperm whale | Artiodactyla | Physeteridae | 35 |
| *Physeter catodon* | Sperm whale | Artiodactyla | Physeteridae | 20 |
| *Mesoplodon densirostris* | Blainville's beaked whale | Artiodactyla | Ziphiidae | 35 |
| *Hipposideros armiger* | Great roundleaf bat | Chiroptera | Hipposideridae | 20 |
| *Molossus molossus* | Pallas's mastiff Bat | Chiroptera | Molossidae | 43 |
| *Artibeus jamaicensis* | Jamaican fruit-eating bat | Chiroptera | Phyllostomidae | 43 |
| *Desmodus rotundus* | Common vampire bat | Chiroptera | Phyllostomidae | 43 |
| *Phyllostomus discolor* | Pale spear-nosed bat | Chiroptera | Phyllostomidae | 43 |
| *Sturnira hondurensis* | Honduran yellow-shouldered bat | Chiroptera | Phyllostomidae | 43 |
| *Pteropus alecto* | Black flying fox | Chiroptera | Pteropodidae | 20 |
| *Pteropus vampyrus* | Large flying fox | Chiroptera | Pteropodidae | 20 |
| *Rhinolophus sinicus* | Chinese rufous horseshoe bat | Chiroptera | Rhinolophidae | 20 |
| *Eptesicus fuscus* | Big brown bat | Chiroptera | Vespertilionidae | 20 |
| *Miniopterus natalensis* | Long-fingered bats | Chiroptera | Vespertilionidae | 20 |
| *Myotis brandtii* | Brandt's myotis | Chiroptera | Vespertilionidae | 20 |
| *Myotis davidii* | David's myotis | Chiroptera | Vespertilionidae | 20 |
| *Myotis lucifugus* | Little brown bat | Chiroptera | Vespertilionidae | 20 |
| *Myotis myotis* | Mouse-eared bat | Chiroptera | Vespertilionidae | 43 |
| *Pipistrellus kuhlii* | Kuhl's pipistrelle | Chiroptera | Vespertilionidae | 43 |
| *Jaculus jaculus* | Lesser Egyptian jerboa | Rodentia | Dipodidae | 20 |
| *Typhlomys cinereus* | Soft-furred tree mouse | Rodentia | Platacanthomyidae | 6 |

**Table S2.** 53 non-echolocating mammalian species from OrthoMaM used as outgroup to identify molecular convergence.

| Species | Common Name | Order | Family | References |
| --- | --- | --- | --- | --- |
| *Bison bison* | American bison | Artiodactyla | Bovidae | 45, 46, 47 |
| *Bos taurus* | Cattle | Artiodactyla | Bovidae | 45, 48 |
| *Bubalus bubalis* | Water buffalo | Artiodactyla | Bovidae | 45, 47 |
| *Capra hircus* | Goat | Artiodactyla | Bovidae | 45, 49 |
| *Ovis aries* | Sheep | Artiodactyla | Bovidae | 45, 50 |
| *Pantholops hodgsonii* | Chiru | Artiodactyla | Bovidae | 47, 51 |
| *Camelus ferus* | Wild Bactrian camel | Artiodactyla | Camelidae | 45, 47 |
| *Vicugna pacos* | Alpaca | Artiodactyla | Camelidae | 45, 47 |
| *Odocoileus virginianus* | White-tailed deer | Artiodactyla | Cervidae | 45, 46, 47 |
| *Canis familiaris* | Dog | Carnivora | Canidae | 45, 52 |
| *Acinonyx jubatus* | Cheetah | Carnivora | Felidae | 45, 46, 47 |
| *Enhydra lutris* | Sea otter | Carnivora | Mustelidae | 45, 46, 47 |
| *Odobenus rosmarus* | Walrus | Carnivora | Odobenidae | 46, 48 |
| *Leptonychotes weddellii* | Weddell seal | Carnivora | Phocidae | 46, 53 |
| *Ailuropoda melanoleuca* | Giant panda | Carnivora | Ursidae | 45, 46, 47, 48 |
| *Ursus maritimus* | Polar bear | Carnivora | Ursidae | 45, 46, 47 |
| *Erinaceus europaeus* | Western European hedgehog | Eulipotyphla | Erinaceidae | 45, 46, 47, 48 |
| *Condylura cristata* | Star-nosed mole | Eulipotyphla | Talpidae | 45, 46, 47 |
| *Procavia capensis* | Cape rock hyrax | Hyracoidea | Procaviidae | 45, 46, 47, 48 |
| *Oryctolagus cuniculus* | Rabbit | Lagomorpha | Leporidae | 45, 46, 47 |
| *Ochotona princeps* | American pika | Lagomorpha | Ochotonidae | 45, 46, 47, 48 |
| *Equus caballus* | Horse | Perissodactyla | Equidae | 45, 47, 48 |
| *Ceratotherium simum* | White rhinoceros | Perissodactyla | Rhinocerotidae | 45, 46, 47, 48 |
| *Callithrix jacchus* | White-tufted-ear | Primates | Callitrichinae | 45, 46, 47, 48 |
| *Cebus capucinus* | White-faced sapajou | Primates | Cebidae | 45, 46, 47 |
| *Saimiri boliviensis* | Bolivian squirrel monkey | Primates | Cebidae | 45, 46, 47 |
| *Cercocebus atys* | Sooty mangabey | Primates | Cercopithecidae | 45, 46, 47 |
| *Chlorocebus sabaeus* | Green monkey | Primates | Cercopithecidae | 45, 47 |
| *Colobus angolensis* | Angolan colobus | Primates | Cercopithecidae | 45, 46, 47 |
| *Macaca mulatta* | Rhesus monkey | Primates | Cercopithecidae | 45, 46, 47, 48 |
| *Mandrillus leucophaeus* | Drill | Primates | Cercopithecidae | 45, 46, 47 |
| *Papio anubis* | Olive baboon | Primates | Cercopithecidae | 45, 46, 47 |
| *Ptilocolobus tephrosceles* | Uganda colobus monkey | Primates | Cercopithecidae | 45, 54 |
| *Rhinopithecus bieti* | Black snub-nosed monkey | Primates | Cercopithecidae | 45, 46, 47 |
| *Gorilla gorilla* | Western gorilla | Primates | Hominidae | 45, 46, 47 |
| *Homo sapiens* | Human | Primates | Hominidae | 45, 46, 48 |
| *Pan troglodytes* | Chimpanzee | Primates | Hominidae | 45, 46, 47 |
| *Pongo abelii* | Sumatran orangutan | Primates | Hominidae | 45, 47 |
| *Nomascus leucogenys* | Northern white-cheeked gibbon | Primates | Hylobatidae | 45, 46, 47, 48 |
| *Propithecus coquereli* | Coquerel's sifaka | Primates | Indriidae | 45, 46, 47 |
| *Loxodonta africana* | African savanna elephant | Proboscidea | Elephantidae | 45, 46, 47, 48 |
| *Fukomys damarensis* | Damara mole rat | Rodentia | Bathyergidae | 46, 55 |
| *Heterocephalus glaber* | Naked mole-rat female | Rodentia | Bathyergidae | 46, 48, 56 |
| *Cavia porcellus* | Domestic guinea pig | Rodentia | Caviidae | 45, 48 |
| *Microtus ochrogaster* | prairie vole | Rodentia | Cricetidae | 45, 46, 47 |
| *Meriones unguiculatus* | Mongolian gerbil | Rodentia | Muridae | 46, 47 |
| *Mus musculus* | House mouse | Rodentia | Muridae | 6 |
| *Octodon degus* | Degu | Rodentia | Octodontidae | 45, 46, 47 |
| *Ictidomys tridecemlineatus* | Thirteen-lined ground squirrel | Rodentia | Sciuridae | 45, 46, 48 |
| *Marmota marmota* | European marmot | Rodentia | Sciuridae | 47, 57 |
| *Nannospalax galili* | Upper Galilee mountains blind mole rat | Rodentia | Spalacidae | 58 |
| *Tupaia belangeri* | Northern tree shrew | Scandentia | Tupaiaidae | 47, 59 |
| *Trichechus manatus* | West Indian manatee | Sirenia | Trichechidae | 46, 48 |

**Table S3.**
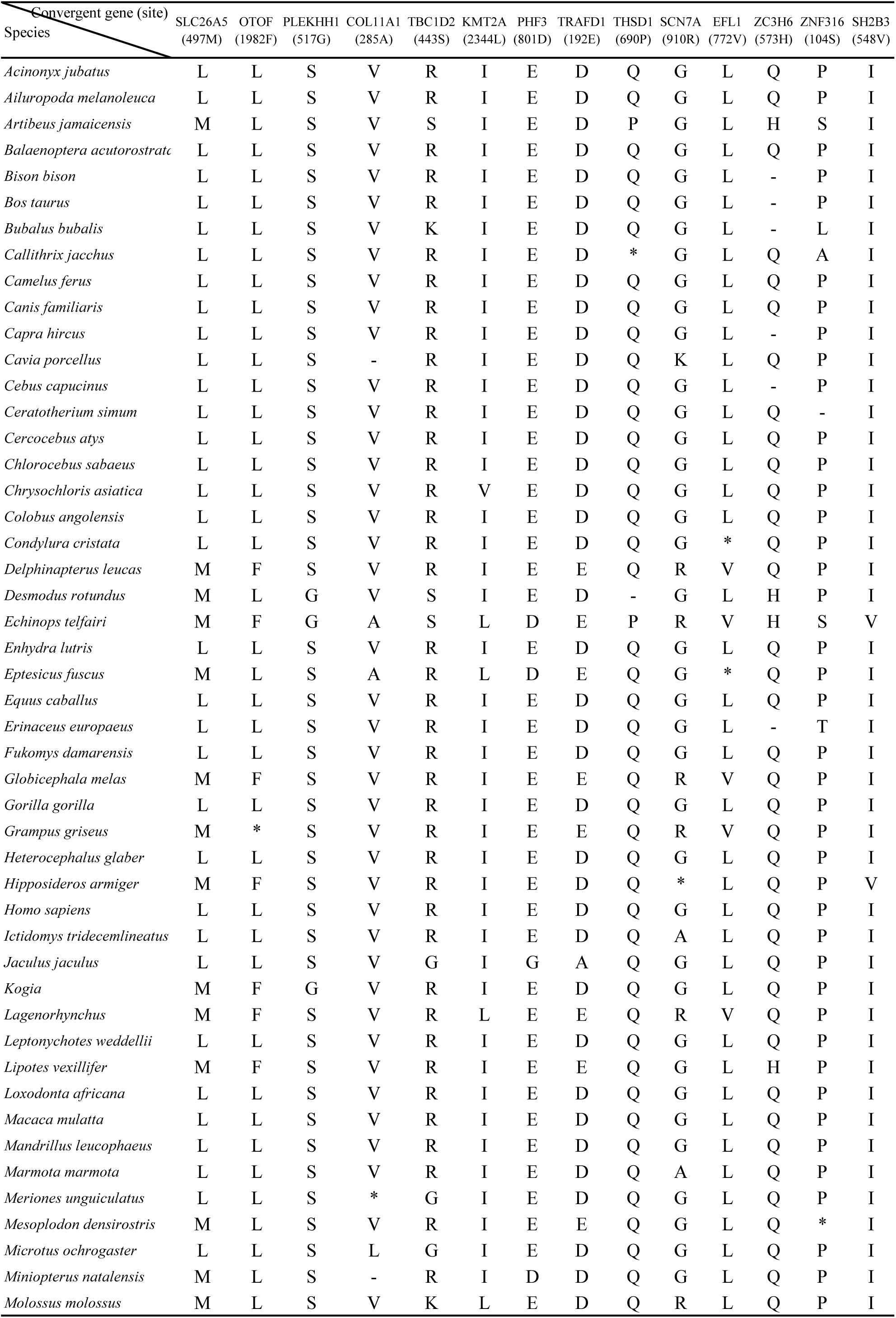
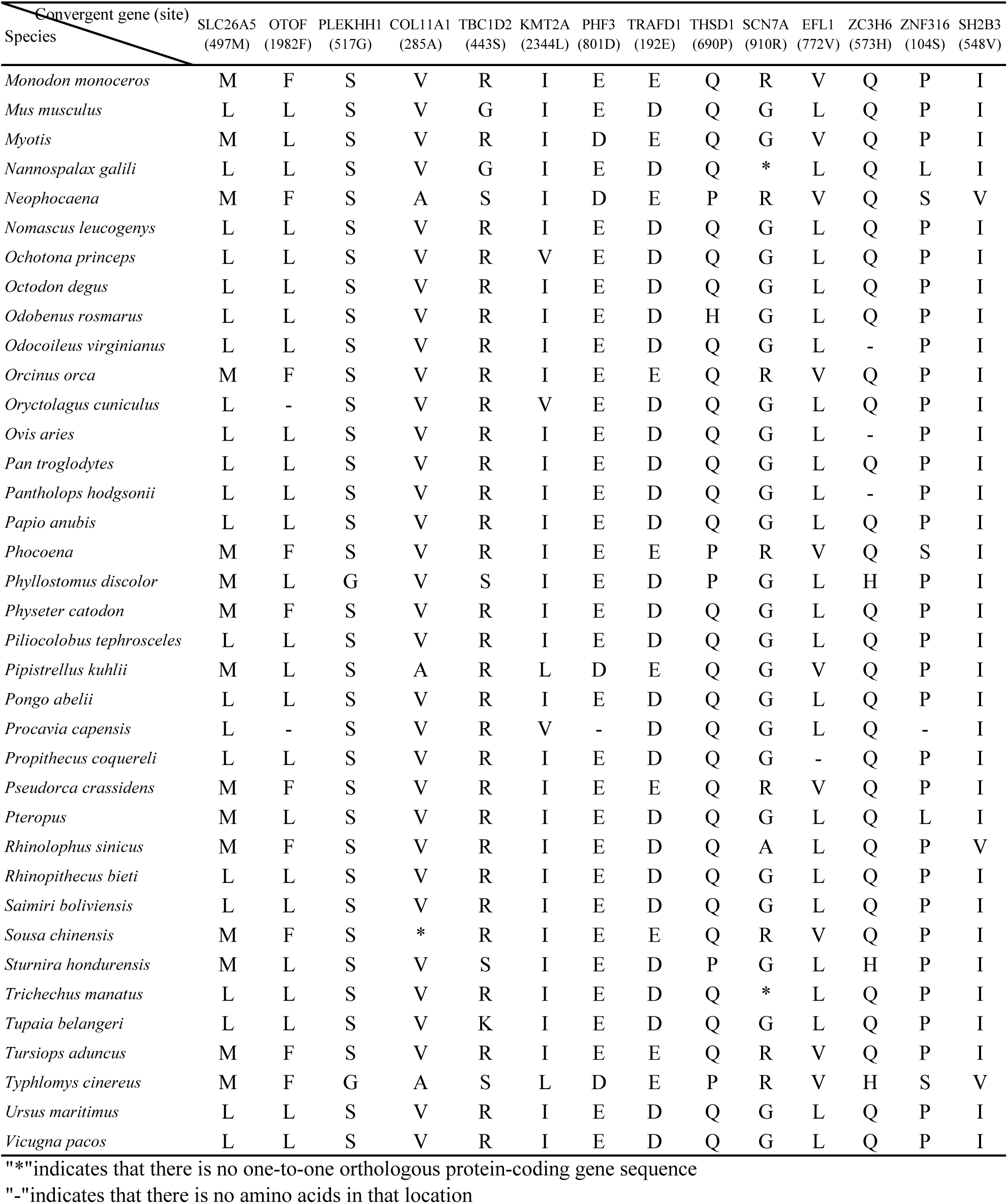
14 convergent amino acid substitutions across 85 representative echolocating and non-echolocating mammalian genera.

| Convergent gene (site)<br>Species | SLC26A5<br>(497M) | OTOF<br>(1982F) | PLEKHH1<br>(517G) | COL11A1<br>(285A) | TBC1D2<br>(443S) | KMT2A<br>(2344L) | PHF3<br>(801D) | TRAFD1<br>(192E) | THSD1<br>(690P) | SCN7A<br>(910R) | EFL1<br>(772V) | ZC3H6<br>(573H) | ZNF316<br>(104S) | SH2B3<br>(548V) |
| --- | --- | --- | --- | --- | --- | --- | --- | --- | --- | --- | --- | --- | --- | --- |
| *Acinonyx jubatus* | L | L | S | V | R | I | E | D | Q | G | L | Q | P | I |
| *Ailuropoda melanoleuca* | L | L | S | V | R | I | E | D | Q | G | L | Q | P | I |
| *Artibeus jamaicensis* | M | L | S | V | S | I | E | D | P | G | L | H | S | I |
| *Balaenoptera acutorostrata* | L | L | S | V | R | I | E | D | Q | G | L | Q | P | I |
| *Bison bison* | L | L | S | V | R | I | E | D | Q | G | L | - | P | I |
| *Bos taurus* | L | L | S | V | R | I | E | D | Q | G | L | - | P | I |
| *Bubalus bubalis* | L | L | S | V | K | I | E | D | Q | G | L | - | L | I |
| *Callithrix jacchus* | L | L | S | V | R | I | E | D | * | G | L | Q | A | I |
| *Camelus ferus* | L | L | S | V | R | I | E | D | Q | G | L | Q | P | I |
| *Canis familiaris* | L | L | S | V | R | I | E | D | Q | G | L | Q | P | I |
| *Capra hircus* | L | L | S | V | R | I | E | D | Q | G | L | - | P | I |
| *Cavia porcellus* | L | L | S | - | R | I | E | D | Q | K | L | Q | P | I |
| *Cebus capucinus* | L | L | S | V | R | I | E | D | Q | G | L | - | P | I |
| *Ceratotherium simum* | L | L | S | V | R | I | E | D | Q | G | L | Q | - | I |
| *Cercocebus atys* | L | L | S | V | R | I | E | D | Q | G | L | Q | P | I |
| *Chlorocebus sabaeus* | L | L | S | V | R | I | E | D | Q | G | L | Q | P | I |
| *Chrysochloris asiatica* | L | L | S | V | R | V | E | D | Q | G | L | Q | P | I |
| *Colobus angolensis* | L | L | S | V | R | I | E | D | Q | G | L | Q | P | I |
| *Condylura cristata* | L | L | S | V | R | I | E | D | Q | G | * | Q | P | I |
| *Delphinapterus leucas* | M | F | S | V | R | I | E | E | Q | R | V | Q | P | I |
| *Desmodus rotundus* | M | L | G | V | S | I | E | D | - | G | L | H | P | I |
| *Echinops telfairi* | M | F | G | A | S | L | D | E | P | R | V | H | S | V |
| *Enhydra lutris* | L | L | S | V | R | I | E | D | Q | G | L | Q | P | I |
| *Eptesicus fuscus* | M | L | S | A | R | L | D | E | Q | G | * | Q | P | I |
| *Equus caballus* | L | L | S | V | R | I | E | D | Q | G | L | Q | P | I |
| *Erinaceus europaeus* | L | L | S | V | R | I | E | D | Q | G | L | - | T | I |
| *Fukomys damarensis* | L | L | S | V | R | I | E | D | Q | G | L | Q | P | I |
| *Globicephala melas* | M | F | S | V | R | I | E | E | Q | R | V | Q | P | I |
| *Gorilla gorilla* | L | L | S | V | R | I | E | D | Q | G | L | Q | P | I |
| *Grampus griseus* | M | * | S | V | R | I | E | E | Q | R | V | Q | P | I |
| *Heterocephalus glaber* | L | L | S | V | R | I | E | D | Q | G | L | Q | P | I |
| *Hipposideros armiger* | M | F | S | V | R | I | E | D | Q | * | L | Q | P | V |
| *Homo sapiens* | L | L | S | V | R | I | E | D | Q | G | L | Q | P | I |
| *Ictidomys tridecemlineatus* | L | L | S | V | R | I | E | D | Q | A | L | Q | P | I |
| *Jaculus jaculus* | L | L | S | V | G | I | G | A | Q | G | L | Q | P | I |
| *Kogia* | M | F | G | V | R | I | E | D | Q | G | L | Q | P | I |
| *Lagenorhynchus* | M | F | S | V | R | L | E | E | Q | R | V | Q | P | I |
| *Leptonychotes weddellii* | L | L | S | V | R | I | E | D | Q | G | L | Q | P | I |
| *Lipotes vexillifer* | M | F | S | V | R | I | E | E | Q | G | L | H | P | I |
| *Loxodonta africana* | L | L | S | V | R | I | E | D | Q | G | L | Q | P | I |
| *Macaca mulatta* | L | L | S | V | R | I | E | D | Q | G | L | Q | P | I |
| *Mandrillus leucophaeus* | L | L | S | V | R | I | E | D | Q | G | L | Q | P | I |
| *Marmota marmota* | L | L | S | V | R | I | E | D | Q | A | L | Q | P | I |
| *Meriones unguiculatus* | L | L | S | * | G | I | E | D | Q | G | L | Q | P | I |
| *Mesoplodon densirostris* | M | L | S | V | R | I | E | E | Q | G | L | Q | * | I |
| *Microtus ochrogaster* | L | L | S | L | G | I | E | D | Q | G | L | Q | P | I |
| *Miniopterus natalensis* | M | L | S | - | R | I | D | D | Q | G | L | Q | P | I |
| *Molossus molossus* | M | L | S | V | K | L | E | D | Q | R | L | Q | P | I |
| *Monodon monoceros* | M | F | S | V | R | I | E | E | Q | R | V | Q | P | I |
| *Mus musculus* | L | L | S | V | G | I | E | D | Q | G | L | Q | P | I |
| *Myotis* | M | L | S | V | R | I | D | E | Q | G | V | Q | P | I |
| *Nannospalax galili* | L | L | S | V | G | I | E | D | Q | * | L | Q | L | I |
| *Neophocaena* | M | F | S | A | S | I | D | E | P | R | V | Q | S | V |
| *Nomascus leucogenys* | L | L | S | V | R | I | E | D | Q | G | L | Q | P | I |
| *Ochotona princeps* | L | L | S | V | R | V | E | D | Q | G | L | Q | P | I |
| *Octodon degus* | L | L | S | V | R | I | E | D | Q | G | L | Q | P | I |
| *Odobenus rosmarus* | L | L | S | V | R | I | E | D | H | G | L | Q | P | I |
| *Odocoileus virginianus* | L | L | S | V | R | I | E | D | Q | G | L | - | P | I |
| *Orcinus orca* | M | F | S | V | R | I | E | E | Q | R | V | Q | P | I |
| *Oryctolagus cuniculus* | L | - | S | V | R | V | E | D | Q | G | L | Q | P | I |
| *Ovis aries* | L | L | S | V | R | I | E | D | Q | G | L | - | P | I |
| *Pan troglodytes* | L | L | S | V | R | I | E | D | Q | G | L | Q | P | I |
| *Pantholops hodgsonii* | L | L | S | V | R | I | E | D | Q | G | L | - | P | I |
| *Papio anubis* | L | L | S | V | R | I | E | D | Q | G | L | Q | P | I |
| *Phocoena* | M | F | S | V | R | I | E | E | P | R | V | Q | S | I |
| *Phyllostomus discolor* | M | L | G | V | S | I | E | D | P | G | L | H | P | I |
| *Physeter catodon* | M | F | S | V | R | I | E | D | Q | G | L | Q | P | I |
| *Piliocolobus tephrosceles* | L | L | S | V | R | I | E | D | Q | G | L | Q | P | I |
| *Pipistrellus kuhlii* | M | L | S | A | R | L | D | E | Q | G | V | Q | P | I |
| *Pongo abelii* | L | L | S | V | R | I | E | D | Q | G | L | Q | P | I |
| *Procapra capensis* | L | - | S | V | R | V | - | D | Q | G | L | Q | - | I |
| *Propithecus coquereli* | L | L | S | V | R | I | E | D | Q | G | - | Q | P | I |
| *Pseudorca crassidens* | M | F | S | V | R | I | E | E | Q | R | V | Q | P | I |
| *Pteropus* | M | L | S | V | R | I | E | D | Q | G | L | Q | L | I |
| *Rhinolophus sinicus* | M | F | S | V | R | I | E | D | Q | A | L | Q | P | V |
| *Rhinopithecus bieti* | L | L | S | V | R | I | E | D | Q | G | L | Q | P | I |
| *Saimiri boliviensis* | L | L | S | V | R | I | E | D | Q | G | L | Q | P | I |
| *Sousa chinensis* | M | F | S | * | R | I | E | E | Q | R | V | Q | P | I |
| *Sturnira hondurensis* | M | L | S | V | S | I | E | D | P | G | L | H | P | I |
| *Trichechus manatus* | L | L | S | V | R | I | E | D | Q | * | L | Q | P | I |
| *Tupaia belangeri* | L | L | S | V | K | I | E | D | Q | G | L | Q | P | I |
| *Tursiops aduncus* | M | F | S | V | R | I | E | E | Q | R | V | Q | P | I |
| *Typhlomys cinereus* | M | F | G | A | S | L | D | E | P | R | V | H | S | V |
| *Ursus maritimus* | L | L | S | V | R | I | E | D | Q | G | L | Q | P | I |
| *Vicugna pacos* | L | L | S | V | R | I | E | D | Q | G | L | Q | P | I |

"\*" indicates that there is no one-to-one orthologous protein-coding gene sequence
"-" indicates that there is no amino acids in that location

**Table S4.**
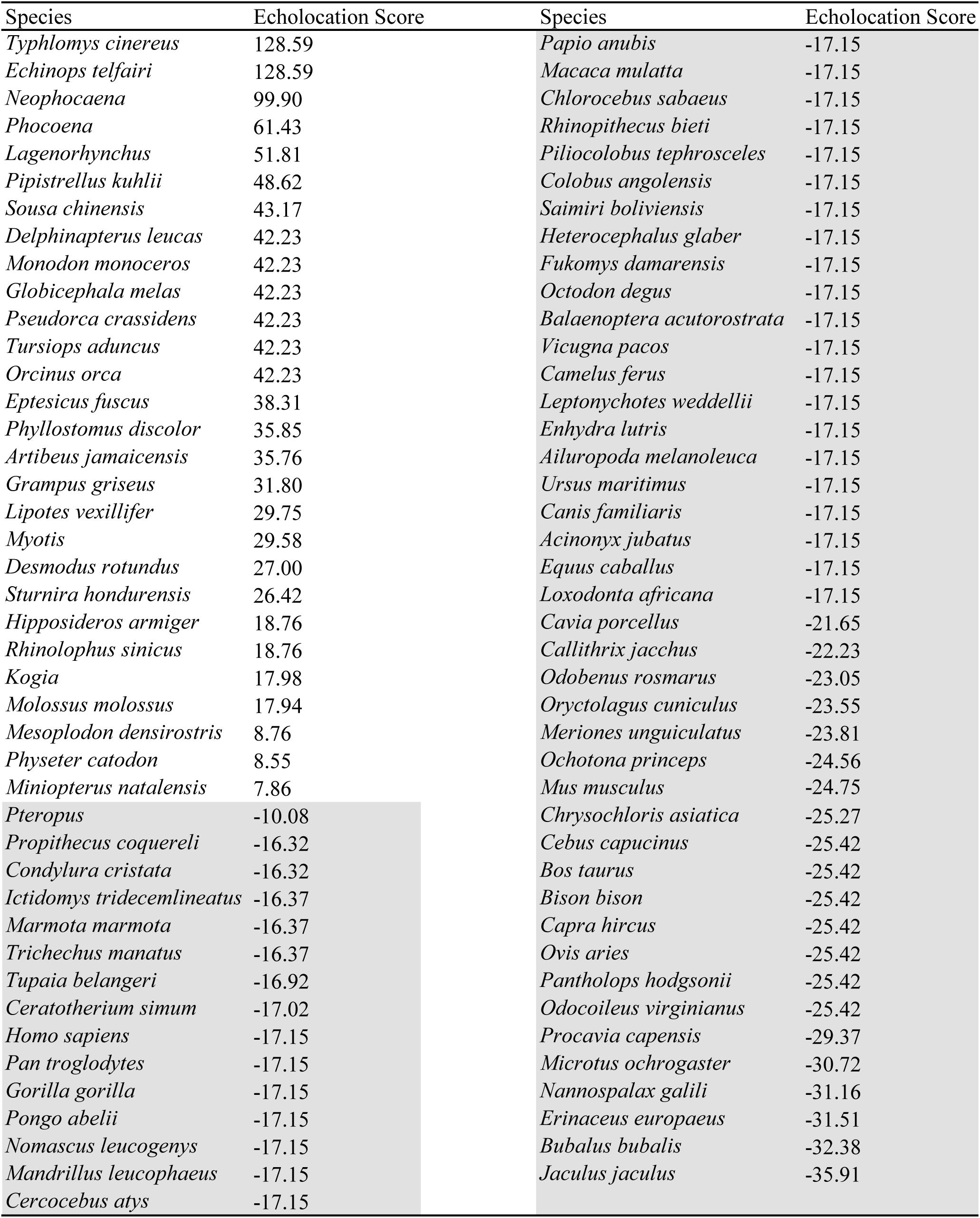
The Echolocation Score of the training dataset.

**Table S5.** 74 mammalian species used to predict new putative echolocators from Zoonomia Project.

| Species | Common name | Order | Family | Single Copy (%) |
| --- | --- | --- | --- | --- |
| *Vulpes lagopus* | Arctic fox | Carnivora | Canidae | 80.3 |
| *Cryptoprocta ferox* | Fossa | Carnivora | Eupleridae | 84.9 |
| *Panthera onca* | Jaguar | Carnivora | Felidae | 78.0 |
| *Helogale parvula* | Dwarf mongoose | Carnivora | Herpestidae | 83.2 |
| *Mungos mungo* | South African banded mongoose | Carnivora | Herpestidae | 86.6 |
| *Suricata suricatta* | Meerkat | Carnivora | Herpestidae | 85.0 |
| *Hyaena hyaena* | Striped hyena | Carnivora | Hyaenidae | 70.1 |
| *Spilogale gracilis* | Western spotted skunk | Carnivora | Mephitidae | 76.1 |
| *Mellivora capensis* | Honey badger | Carnivora | Mustelidae | 72.1 |
| *Pteronura brasiliensis* | Giant otter | Carnivora | Mustelidae | 81.4 |
| *Zalophus californianus* | California sea lion | Carnivora | Otariidae | 83.3 |
| *Mirounga angustirostris* | Northern elephant seal | Carnivora | Phocidae | 71.6 |
| *Paradoxurus hermaphroditus* | Asian palm civet | Carnivora | Viverridae | 74.2 |
| *Antilocapra americana* | Pronghorn | Cetartiodactyla | Antilocapridae | 78.3 |
| *Eubalaena japonica* | North Pacific right whale | Cetartiodactyla | Balaenidae | 67.7 |
| *Beatragus hunteri* | Hirola | Cetartiodactyla | Bovidae | 75.9 |
| *Hemitragus hylocrius* | Nilgiri tahr | Cetartiodactyla | Bovidae | 78.4 |
| *Ovis canadensis cremnobates* | Peninsular bighorn sheep | Cetartiodactyla | Bovidae | 75.2 |
| *Rangifer tarandus* | Siberian reindeer | Cetartiodactyla | Cervidae | 77.8 |
| *Hippopotamus amphibius* | Hippopotamus | Cetartiodactyla | Hippopotamidae | 79.0 |
| *Moschus moschiferus* | Siberian musk deer | Cetartiodactyla | Moschidae | 71.6 |
| *Catagonus wagneri* | Chacoan peccary | Cetartiodactyla | Tayassuidae | 85.0 |
| *Tragulus javanicus* | Java lesser chevrotain | Cetartiodactyla | Tragulidae | 75.2 |
| *Mesoplodon bidens* | Sowerby's beaked whale | Cetartiodactyla | Ziphiidae | 61.6 |
| *Hipposideros galeritus* | Cantor's leafnosed bat | Chiroptera | Hipposideridae | 62.9 |
| *Megaderma lyra* | Greater false vampire bat | Chiroptera | Megadermatidae | 87.0 |
| *Mormoops blainvillei* | Ghost-faced bat | Chiroptera | Mormoopidae | 84.6 |
| *Noctilio leporinus* | Greater bulldog bat | Chiroptera | Noctilionidae | 83.7 |
| *Anoura caudifer* | Tailed tailless bat | Chiroptera | Phyllostomidae | 85.6 |
| *Micronycteris hirsuta* | Hairy big-eared bat | Chiroptera | Phyllostomidae | 75.1 |
| *Tonatia saurophila* | Stripe-headed round-eared bat | Chiroptera | Phyllostomidae | 83.2 |
| *Macroglossus sobrinus* | Long-tongued fruit bat | Chiroptera | Pteropodidae | 90.3 |
| *Rhinolophus ferrumequinum* | Greater horseshoe bat | Chiroptera | Rhinolophidae | 86.8 |
| *Antrozous pallidus* | Pallid bat | Chiroptera | Vespertilionidae | 76.0 |
| *Miniopterus schreibersii* | Common bentwing bat | Chiroptera | Vespertilionidae | 84.9 |
| *Solenodon paradoxus* | Hispaniolan solenodon | Eulipotyphla | Solenodontidae | 90.1 |
| *Scalopus aquaticus* | Eastern mole | Eulipotyphla | Talpidae | 81.9 |
| *Uropsilus gracilis* | Gracile shrewlike mole | Eulipotyphla | Talpidae | 64.8 |
| *Heterohyrax brucei* | African yellowspotted rock hyrax | Hyracoidea | Procaviidae | 75.6 |
| *Diceros bicornis* | Black rhinoceros | Perissodactyla | Rhinocerotidae | 82.9 |
| *Tapirus indicus* | Malayan tapir | Perissodactyla | Tapiridae | 83.7 |
| *Tapirus terrestris* | South American tapir | Perissodactyla | Tapiridae | 81.6 |
| *Manis tricuspis* | Tree pangolin | Pholidota | Manidae | 90.4 |
| *Alouatta palliata mexicana* | Mexican howler monkey | Primates | Atelidae | 68.0 |
| *Ateles geoffroyi* | Geoffroy's spider monkey | Primates | Atelidae | 77.7 |
| *Saguinus imperator* | Emperor tamarin | Primates | Cebidae | 73.7 |
| *Erythrocebus patas* | Patas monkey | Primates | Cercopithecidae | 61.5 |
| *Pygathrix nemaeus* | Red-shanked douc | Primates | Cercopithecidae | 70.8 |
| *Cheirogaleus medius* | Fat-tailed dwarf lemur | Primates | Cheirogaleidae | 79.0 |
| *Mirza coquereli* | Coquerel's giant mouse lemur | Primates | Cheirogaleidae | 65.9 |
| *Daubentonia madagascariensis* | Aye-aye | Primates | Daubentoniidae | 90.4 |
| *Indri indri* | Indri | Primates | Indridae | 60.0 |
| *Lemur catta* | Ring tailed lemur | Primates | Lemuridae | 75.0 |
| *Callicebus donacophilus* | White-eared titi | Primates | Pitheciidae | 60.2 |
| *Pithecia pithecia* | White-faced saki | Primates | Pitheciidae | 74.8 |
| *Aplodontia rufa* | Mountain beaver | Rodentia | Aplodontiidae | 66.8 |
| *Cavia tschudii* | Montane guinea pig | Rodentia | Caviidae | 82.4 |
| *Dolichotis patagonum* | Patagonian mara | Rodentia | Caviidae | 70.2 |
| *Hydrochoerus hydrochaeris* | Capybara | Rodentia | Caviidae | 85.5 |
| *Ondatra zibethicus* | Muskrat | Rodentia | Cricetidae | 79.7 |
| *Sigmodon hispidus* | Hispid cotton rat | Rodentia | Cricetidae | 80.5 |
| *Ctenodactylus gundi* | Common gundi | Rodentia | Ctenodactylidae | 89.7 |
| *Ctenomys sociabilis* | Social tuco-tuco | Rodentia | Ctenomyidae | 63.3 |
| *Dasyprocta punctata* | Central American agouti | Rodentia | Dasyproctidae | 66.4 |
| *Dinomys branickii* | Pacarana | Rodentia | Dinomyidae | 76.5 |
| *Muscardinus avellanarius* | Hazel dormouse | Rodentia | Gliridae | 67.1 |
| *Dipodomys stephensi* | Stephen's kangaroo rat | Rodentia | Heteromyidae | 63.8 |
| *Perognathus longimembris pacificus* | Pacific pocket mouse | Rodentia | Heteromyidae | 60.5 |
| *Hystrix cristata* | Northern crested porcupine | Rodentia | Hystriidae | 76.0 |
| *Acomys cahirinus* | Cairo spiny mouse | Rodentia | Muridae | 70.2 |
| *Cricetomys gambianus* | Gambian pouched rat | Rodentia | Nesomyidae | 83.7 |
| *Pedetes capensis* | South African springhare | Rodentia | Pedetidae | 76.0 |
| *Petromus typicus* | Dassie rat | Rodentia | Petromuridae | 60.9 |
| *Xerus inauris* | Cape ground squirrel | Rodentia | Sciuridae | 74.2 |

**Table S6.** 23 mammalian species used to predict new putative echolocators from OrthoMaM.

| Species | Common name | Order | Family | References |
| --- | --- | --- | --- | --- |
| *Sus scrofa* | Pig | Artiodactyla | Suidae | 45, 46, 47, 48 |
| *Felis catus* | Domestic cat | Carnivora | Felidae | 45, 48 |
| *Panthera tigris* | Tiger | Carnivora | Felidae | 45, 46, 47 |
| *Mustela putorius* | European polecat | Carnivora | Mustelidae | 45, 46, 47 |
| *Monachus schauinslandi* | Hawaiian monk seal | Carnivora | Phocidae | 45, 46 |
| *Dasypus novemcinctus* | Nine-banded armadillo | Cingulata | Dasypodidae | 45, 46, 47, 48 |
| *Galeopterus variegatus* | Sunda flying lemur | Dermoptera | Cynocephalidae | 45, 46, 47, 48 |
| *Sorex araneus* | European shrew | Eulipotyphla | Soricidae | 45, 46, 47, 48 |
| *Elephantulus edwardii* | Cape elephant shrew | Macroscelidea | Macroscelididae | 46, 47, 60 |
| *Manis javanica* | Malayan pangolin | Pholidota | Manidae | 45, 47 |
| *Choloepus hoffmanni* | Hoffmann's two-fingered sloth | Pilosa | Megalonychidae | 45, 46, 47, 48 |
| *Aotus nancymaae* | Ma's night monkey | Primates | Aotidae | 45, 46, 47 |
| *Microcebus murinus* | Gray mouse lemur | Primates | Cheirogaleidae | 45, 46, 47, 48 |
| *Otolemur garnettii* | Small-eared galago | Primates | Galagidae | 45, 47 |
| *Carlito syrichta* | Philippine tarsier | Primates | Tarsiidae | 45, 46, 47 |
| *Castor canadensis* | American beaver | Rodentia | Castoridae | 45, 46, 47, 48 |
| *Chinchilla lanigera* | Long-tailed chinchilla | Rodentia | Chinchillidae | 45, 46, 47, 48 |
| *Cricetulus griseus* | Chinese hamster | Rodentia | Cricetidae | 61 |
| *Mesocricetus auratus* | Golden hamster | Rodentia | Cricetidae | 45, 46, 47, 48 |
| *Peromyscus maniculatus* | North American deer mouse | Rodentia | Cricetidae | 45, 46, 47 |
| *Dipodomys ordii* | Ord's kangaroo rat | Rodentia | Heteromyidae | 45, 46, 47 |
| *Rattus norvegicus* | Norway rat | Rodentia | Muridae | 45, 47 |
| *Orycteropus afer* | Aardvark | Tubulidentata | Orycteropodidae | 46, 47, 48 |

**Table S7.** The Echolocation Score of the predicting dataset.

| Species | Echolocation Score | Species | Echolocation |
| --- | --- | --- | --- |
| *Antrozous pallidus* | 26.99 | *Tapirus terrestris* | -14.60 |
| *Anoura caudifer* | 26.55 | *Eubalaena japonica* | -15.07 |
| *Micronycteris hirsuta* | 26.55 | *Spilogale gracilis* | -15.20 |
| *Tonatia saurophila* | 22.62 | *Pithecia pithecia* | -15.30 |
| *Rhinolophus ferrumequinum* | 17.98 | *Mellivora capensis* | -15.38 |
| *Mormoops blainvillei* | 16.65 | *Pedetes capensis* | -15.38 |
| *Uropsilus gracilis* | 15.88 | *Saguinus imperator* | -15.38 |
| *Sorex araneus* | 12.77 | *Xerus inauris* | -15.38 |
| *Dasypus novemcinctus* | 9.88 | *Vulpes lagopus* | -16.01 |
| *Miniopterus schreibersii* | 8.69 | *Galeopterus variegatus* | -16.01 |
| *Hipposideros galeritus* | 8.44 | *Catagonus wagneri* | -16.14 |
| *Noctilio leporinus* | 6.97 | *Aplodontia rufa* | -16.19 |
| *Solenodon paradoxus* | 6.88 | *Tragulus javanicus* | -16.19 |
| *Megaderma lyra* | 6.69 | *Cheirogaleus medius* | -16.21 |
| *Mesoplodon bidens* | 6.46 | *Sus scrofa* | -16.27 |
| *Hydrochoerus hydrochaeris* | 6.38 | *Orycteropus afer* | -16.27 |
| *Manis javanica* | 2.66 | *Dolichotis patagonum* | -17.02 |
| *Mirza coquereli* | 0.10 | *Pteronura brasiliensis* | -17.02 |
| *Muscardinus avellanarius* | -0.12 | *Tapirus indicus* | -17.02 |
| *Ctenodactylus gundi* | -1.66 | *Panthera tigris* | -17.02 |
| *Dinomys branickii* | -1.84 | *Ctenomys sociabilis* | -17.15 |
| *Carlito syrichta* | -2.76 | *Daubentonia madagascariensis* | -17.15 |
| *Microcebus murinus* | -2.89 | *Diceros bicornis* | -17.15 |
| *Cricetomys gambianus* | -3.83 | *Lemur catta* | -17.15 |
| *Castor canadensis* | -4.26 | *Chinchilla lanigera* | -17.15 |
| *Indri indri* | -4.30 | *Mustela putorius* | -17.15 |
| *Scalopus aquaticus* | -4.94 | *Felis catus* | -17.15 |
| *Dipodomys stephensi* | -6.05 | *Elephantulus edwardii* | -17.15 |
| *Monachus schauinslandi* | -6.13 | *Hystrix cristata* | -19.64 |
| *Dasyprocta punctata* | -6.22 | *Cavia tschudii* | -20.75 |
| *Acomys cahirinus* | -6.96 | *Callicebus donacophilus* | -21.20 |
| *Aotus nancymaae* | -7.34 | *Panthera onca* | -21.36 |
| *Hippopotamus amphibius* | -7.44 | *Cryptoprocta ferox* | -21.83 |
| *Macroglossus sobrinus* | -10.08 | *Zalophus californianus* | -22.09 |
| *Ondatra zibethicus* | -12.05 | *Choloepus hoffmanni* | -22.63 |
| *Beatragus hunteri* | -12.59 | *Sigmodon hispidus* | -22.98 |
| *Perognathus longimembris* | -13.09 | *Otolemur garnettii* | -23.05 |
| *Suricata suricatta* | -13.29 | *Helogale parvula* | -23.37 |
| *Manis tricuspis* | -13.32 | *Hemitragus hylocrius* | -23.44 |
| *Paradoxurus hermaphroditus* | -13.32 | *Mesocricetus auratus* | -23.81 |
| *Heterohyrax brucei* | -13.36 | *Cricetulus griseus* | -23.81 |
| *Erythrocebus patas* | -13.37 | *Pygathrix nemaeus* | -24.37 |
| *Dipodomys ordii* | -13.81 | *Mungos mungo* | -24.43 |
| *Ateles geoffroyi* | -14.16 | *Rangifer tarandus* | -24.59 |
| *Hyaena hyaena* | -14.26 | *Rattus norvegicus* | -24.75 |
| *Antilocapra americana* | -14.27 | *Moschus moschiferus* | -25.29 |
| *Peromyscus maniculatus* | -14.38 | *Ovis canadensis cremnobates* | -25.29 |
| *Alouatta palliata mexicana* | -14.47 | *Petromus typicus* | -30.30 |
| *Mirounga angustirostris* | -14.47 |  |  |

**Table S8.** Acoustic variables of ultrasonic pulses from *Uropsilus* species.

| Species | *U. gracilis* | *U. investigator* | *U. nivatus* | *U. Soricipes* |
| --- | --- | --- | --- | --- |
| Acoustic parameters |  |  |  |  |
| N | 16 | 7 | 11 | 14 |
| n | 480 | 210 | 330 | 420 |
| Pulse duration (ms) | 0.80 ± 0.04<br>(0.4-1.23) | 1.05 ± 0.06<br>(0.97-1.41) | 0.77 ± 0.05<br>(0.46-1.53) | 0.63 ± 0.02<br>(0.55-1.75) |
| Pulse interval between two single pulse groups (ms) | 31.74 ± 0.64<br>(28.14-31.74) | 31.74 ± 2.15<br>(22.07-38.72) | 30.54 ± 0.33<br>(23.41-37.09) | 36.66 ± 2.16<br>(29.47-48.94) |
| Pulse interval within dyad pulse group (ms) | 3.14 ± 0.03<br>(1.78-7.2) | 3.18 ± 0.22<br>(2.08-6.68) | 3.18 ± 0.22<br>(2.08-6.68) | 3.68 ± 0.35<br>(2.21-6.20) |
| Pulse interval within triad pulse group (ms) | 2.85 ± 0.21<br>(2.32-4.44) | 2.67 ± 0.31<br>(1.52-3.57) | 2.67 ± 0.31<br>(1.52-3.57) | 3.17 ± 0.19<br>(2.43-4.87) |
| f<sub>max</sub> (kHz) | 107.68 ± 1.83<br>(79-147) | 92.86 ± 0.82<br>(90.49-95.29) | 100.47 ± 2.85<br>(71.40-125.90) | 102.43 ± 2.27<br>(91.43-113.95) |
| f<sub>min</sub> (kHz) | 42.2 ± 1.38<br>(22-68) | 30.96 ± 2.41<br>(22.26-40.93) | 26.19 ± 0.74<br>(20.50-33.30) | 28.24 ± 1.63<br>(22.42-42.71) |
| Bandwidth (kHz) | 65.48 ± 0.56<br>(56-79) | 61.86 ± 2.58<br>(51.65-70.15) | 74.23 ± 2.72<br>(43.50-90.20) | 74.15 ± 2.33<br>(67.84-86.72) |
| Peak frequency (kHz) | 63.03 ± 1.37<br>(42-89) | 57.66 ± 1.49<br>(51.94-64.56) | 47.63 ± 1.05<br>(35.71-56.20) | 57.85 ± 1.31<br>(51.27-65.72) |

N: Number of individuals
n: Number of pulses
Results are presented as Mean ± SEM (range).

**Table S9.** Performance of *Uropsilus investigator* under different experimental treatments and conditions.

| Test groups | Treatment | Experiment apparatus | Sample # | Percentage of exploration time in the left of monitored sector (%) | Percentage of exploration time in monitored sector (%) | Percentage of exploration time in the opposite sector (%) | # of ultrasonic emissions per sec in monitored sector |
| --- | --- | --- | --- | --- | --- | --- | --- |
| Small disks | None | Disc-small discs | UIN01 | 14.46 ± 1.99 | 13.85 ± 2.53 | 15.33 ± 4.76 | 14.8 ± 7.74 |
|  |  |  | UIN02 | 13.75 ± 2.33 | 14.12 ± 3.34 | 5.86 ± 2.06 | 30.23 ± 12.79 |
|  |  |  | UIN03 | 11.67 ± 0.7 | 11.77 ± 2.05 | 9.71 ± 1.77 | 24.9 ± 7.97 |
|  |  |  | UIN04 | 12.69 ± 0.98 | 10.95 ± 0.65 | 14.05 ± 1.02 | 19.81 ± 4.82 |
|  |  |  | UIN05 | 13.7 ± 4.2 | 14.85 ± 5.15 | 11.23 ± 1.65 | 17.56 ± 11.16 |
|  |  |  | UIN06 | 12.13 ± 4.18 | 14.26 ± 3.87 | 10.15 ± 3.13 | 26.07 ± 11.58 |
|  |  |  | UIN07 | 9.38 ± 1.09 | 15.7 ± 2.12 | 13.67 ± 2.29 | 29.79 ± 3.82 |
| Platform | None | Disc-platform | UIN01 | 9.97 ± 2.01 | 28.58 ± 4.73 | 6.66 ± 2.09 | 42.87 ± 6.6 |
|  |  |  | UIN02 | 6.52 ± 5.29 | 23.02 ± 4.79 | 4.55 ± 2.63 | 37.92 ± 10.61 |
|  |  |  | UIN03 | 11.77 ± 0.86 | 21.49 ± 4.37 | 10.97 ± 4.49 | 47.67 ± 13.71 |
|  |  |  | UIN04 | 20.52 ± 3.14 | 22.83 ± 3.18 | 9.94 ± 3 | 49 ± 5.38 |
|  |  |  | UIN05 | 18.74 ± 4.5 | 25.15 ± 7.85 | 8.15 ± 3.03 | 41.93 ± 8.93 |
|  |  |  | UIN06 | 13.32 ± 9.73 | 24.1 ± 5.49 | 2.76 ± 1.83 | 45.3 ± 5.81 |
|  |  |  | UIN07 | 5.49 ± 2.36 | 22.08 ± 2.09 | 5.02 ± 2.27 | 47.58 ± 7.71 |
| Earplug | Ears plugged | Disc-platform | UIN01 | 10.11 ± 3.64 | 9.75 ± 3.2 | 16.19 ± 3.88 | 7.15 ± 3.14 |
|  |  |  | UIN02 | 14.14 ± 2.26 | 12.99 ± 3.4 | 8.87 ± 1.66 | 25.71 ± 6.01 |
|  |  |  | UIN03 | 16.25 ± 0.78 | 13.51 ± 1.41 | 9.08 ± 1.22 | 37.41 ± 5.72 |
|  |  |  | UIN04 | 10.48 ± 3.63 | 11.14 ± 0.61 | 14.4 ± 3.24 | 29.26 ± 2.94 |
|  |  |  | UIN05 | 14.7 ± 1.59 | 15.73 ± 2.27 | 11.65 ± 1.6 | 18.5 ± 5.63 |
|  |  |  | UIN06 | 13.81 ± 2.41 | 14.44 ± 2.73 | 7.85 ± 1.01 | 22.44 ± 5.65 |
|  |  |  | UIN07 | 8.52 ± 1.46 | 11.49 ± 4.63 | 13.87 ± 2.08 | 20.04 ± 4.38 |
| Sham | Hollow tubes | Disc-platform | UIN01 | 12.5 ± 4.38 | 17.85 ± 3.79 | 14.71 ± 5.95 | 46.18 ± 8.67 |
|  |  |  | UIN02 | 17.15 ± 7.91 | 18.04 ± 3.17 | 6.65 ± 2.32 | 41.85 ± 12.41 |
|  |  |  | UIN03 | 11.82 ± 2.3 | 19.86 ± 3.16 | 10.14 ± 1.75 | 34.83 ± 3.86 |
|  |  |  | UIN04 | 14.66 ± 2.66 | 21.81 ± 3.6 | 6.17 ± 2 | 26.24 ± 8.29 |
|  |  |  | UIN05 | 15.11 ± 2.13 | 31.21 ± 4.5 | 8.83 ± 4.5 | 42.62 ± 11.09 |
|  |  |  | UIN06 | 12.04 ± 2.06 | 29.59 ± 7.27 | 7.78 ± 2.25 | 39.39 ± 8.75 |
|  |  |  | UIN07 | 10.01 ± 3.03 | 34.37 ± 9.78 | 4.23 ± 1.34 | 37.81 ± 6.55 |
| Recovery | None | Disc-platform | UIN01 | 8.91 ± 1.37 | 23.55 ± 3.53 | 10.14 ± 3.35 | 39.98 ± 5.68 |
|  |  |  | UIN02 | 10.28 ± 2.34 | 24.55 ± 4.57 | 8.33 ± 3.09 | 43.24 ± 13.01 |
|  |  |  | UIN03 | 16.61 ± 4.51 | 32.93 ± 5.25 | 5.56 ± 1.84 | 30.67 ± 7.72 |
|  |  |  | UIN04 | 10.64 ± 5.88 | 27.14 ± 8.89 | 8.19 ± 2.93 | 43.65 ± 4.3 |
|  |  |  | UIN05 | 12.23 ± 3.09 | 18.77 ± 3.46 | 12.45 ± 2.68 | 36.17 ± 5.61 |
|  |  |  | UIN06 | 9.76 ± 3.98 | 31.27 ± 10.61 | 5.77 ± 2.75 | 37.63 ± 0.95 |
|  |  |  | UIN07 | 26.48 ± 8.85 | 26.46 ± 6.17 | 12.1 ± 3.54 | 39.79 ± 1.92 |

**Table S10.** Performance of *Uropsilus nivatus* under different experimental treatments and conditions.

| Test groups | Treatment | Experiment apparatus | Sample # | Percentage of exploration time in the left of monitored sector (%) | Percentage of exploration time in monitored sector (%) | Percentage of exploration time in the opposite sector (%) | # of ultrasonic emissions per sec in monitored sector |
| --- | --- | --- | --- | --- | --- | --- | --- |
| Small disks | None | Disc-small discs | UNI01 | 11.06 ± 0.33 | 13.68 ± 0.48 | 13.05 ± 0.92 | 16.2 ± 3.65 |
|  |  |  | UNI02 | 11.46 ± 1.31 | 10.59 ± 2.01 | 14.05 ± 2.28 | 27.61 ± 5.29 |
|  |  |  | UNI03 | 10.13 ± 0.89 | 11.25 ± 0.48 | 16.03 ± 2.54 | 26.99 ± 2.85 |
|  |  |  | UNI04 | 11.59 ± 0.92 | 13.2 ± 0.88 | 12.3 ± 1.02 | 23.13 ± 5.92 |
|  |  |  | UNI05 | 12.31 ± 1.47 | 13.25 ± 1.11 | 13.1 ± 0.61 | 20.4 ± 7.92 |
|  |  |  | UNI06 | 13.27 ± 1.89 | 13.92 ± 1.67 | 10.89 ± 2 | 28.19 ± 4.63 |
|  |  |  | UNI07 | 12.77 ± 1.17 | 12.32 ± 0.27 | 12.46 ± 0.86 | 16.96 ± 7.01 |
|  |  |  | UNI08 | 14.64 ± 1.73 | 12.94 ± 1.02 | 11.55 ± 1.12 | 41.64 ± 7.04 |
|  |  |  | UNI09 | 11.36 ± 1.14 | 11.86 ± 1.99 | 13.83 ± 1.67 | 24.75 ± 9.94 |
|  |  |  | UNI10 | 12.02 ± 1.07 | 12.52 ± 0.8 | 12.28 ± 1.18 | 18.37 ± 1.87 |
|  |  |  | UNI11 | 11.51 ± 1.74 | 13.09 ± 0.9 | 13.83 ± 1.26 | 18.74 ± 5.79 |
| Platform | None | Disc-platform | UNI01 | 17.09 ± 4.69 | 26.07 ± 2.01 | 7.05 ± 1.82 | 31.48 ± 9.52 |
|  |  |  | UNI02 | 15.32 ± 2.24 | 23.29 ± 2.37 | 6.28 ± 2.78 | 36.52 ± 11.13 |
|  |  |  | UNI03 | 14.93 ± 1.65 | 24.81 ± 1.5 | 6.36 ± 1.57 | 39.94 ± 4.83 |
|  |  |  | UNI04 | 14.08 ± 2.01 | 23.16 ± 1.75 | 12.2 ± 2.57 | 29.58 ± 1.23 |
|  |  |  | UNI05 | 11.95 ± 0.57 | 21.85 ± 1.53 | 12.35 ± 1.48 | 37.8 ± 5.88 |
|  |  |  | UNI06 | 9.99 ± 2.24 | 22.89 ± 1.31 | 13.73 ± 3.58 | 33.48 ± 5.52 |
|  |  |  | UNI07 | 12.55 ± 1.73 | 21.86 ± 0.86 | 9.53 ± 1.54 | 27.7 ± 5.9 |
|  |  |  | UNI08 | 11.73 ± 0.55 | 23.92 ± 1.56 | 8.13 ± 0.44 | 40.74 ± 6.65 |
|  |  |  | UNI09 | 10.36 ± 1.79 | 35.69 ± 4.96 | 5.4 ± 2.05 | 32.43 ± 10.73 |
|  |  |  | UNI10 | 13.6 ± 2.02 | 30.2 ± 6.77 | 11.19 ± 5.93 | 35.05 ± 2.22 |
|  |  |  | UNI11 | 15.93 ± 0.92 | 28.81 ± 4.5 | 9.07 ± 1.58 | 35.43 ± 5.94 |
| Earplug | Ears plugged | Disc-platform | UNI01 | 12.29 ± 1.2 | 14.01 ± 1.86 | 11.73 ± 0.43 | 8.79 ± 1.49 |
|  |  |  | UNI02 | 11.09 ± 1.97 | 14.41 ± 2.41 | 13.23 ± 1.67 | 28.91 ± 12.82 |
|  |  |  | UNI03 | 13.91 ± 2.08 | 13.51 ± 1.79 | 12.91 ± 1.49 | 12.59 ± 3.39 |
|  |  |  | UNI04 | 10.91 ± 0.83 | 12.69 ± 1.26 | 14.47 ± 1.99 | 23.28 ± 1.27 |
|  |  |  | UNI05 | 12.26 ± 2.97 | 10.63 ± 1.88 | 14.93 ± 1.86 | 13.61 ± 3.29 |
|  |  |  | UNI06 | 15.23 ± 0.96 | 13.4 ± 1.26 | 14.63 ± 1.62 | 13.3 ± 2.72 |
|  |  |  | UNI07 | 12.8 ± 1 | 11.98 ± 0.75 | 15.45 ± 1.2 | 18.68 ± 6.03 |
|  |  |  | UNI08 | 14.72 ± 1.84 | 11.17 ± 0.65 | 14.08 ± 0.76 | 13.99 ± 2.54 |
|  |  |  | UNI09 | 7.91 ± 2.77 | 13.66 ± 0.31 | 19.21 ± 4.35 | 7.95 ± 1.83 |
|  |  |  | UNI10 | 11.52 ± 0.63 | 12.62 ± 2.3 | 14.93 ± 1.16 | 10.47 ± 1.67 |
|  |  |  | UNI11 | 14.12 ± 2.48 | 13.12 ± 1.97 | 11.22 ± 1.2 | 18.71 ± 4.76 |
| Sham | Hollow tubes | Disc-platform | UNI01 | 9.58 ± 0.91 | 23.48 ± 3.56 | 10.15 ± 0.95 | 41.69 ± 6.71 |
|  |  |  | UNI02 | 10.86 ± 3.24 | 27.03 ± 4.78 | 7.93 ± 2.2 | 50.65 ± 6.83 |
|  |  |  | UNI03 | 17.9 ± 3.6 | 25.69 ± 3.92 | 8.92 ± 3.05 | 34.35 ± 3.72 |
|  |  |  | UNI04 | 12.66 ± 0.86 | 24.2 ± 4.05 | 10.23 ± 0.85 | 52.74 ± 7.86 |
|  |  |  | UNI05 | 12.69 ± 2.89 | 24.72 ± 3.85 | 11.5 ± 1.51 | 41.93 ± 3.98 |
|  |  |  | UNI06 | 14.94 ± 3.81 | 22.94 ± 1.67 | 8.83 ± 0.95 | 33.35 ± 4.63 |
|  |  |  | UNI07 | 11.88 ± 0.99 | 24.97 ± 1.82 | 8.49 ± 2.22 | 35.09 ± 5.23 |
|  |  |  | UNI08 | 14.13 ± 1.69 | 33.42 ± 3.52 | 7.24 ± 2.32 | 33.02 ± 9.2 |
|  |  |  | UNI09 | 9.78 ± 2.81 | 30.44 ± 5.99 | 11.61 ± 6.4 | 43.9 ± 9.41 |
|  |  |  | UNI10 | 17.1 ± 1.93 | 40.33 ± 5.7 | 3.72 ± 1.35 | 31.04 ± 7.82 |
|  |  |  | UNI11 | 14.83 ± 2.06 | 34.1 ± 3.38 | 5.35 ± 2.43 | 27.8 ± 8.56 |
| Recovery | None | Disc-platform | UNI01 | 12.21 ± 2.21 | 24.22 ± 2.65 | 10.41 ± 2.77 | 37 ± 3.68 |
|  |  |  | UNI02 | 12.72 ± 3.19 | 28.83 ± 3.47 | 7.18 ± 3.23 | 44.53 ± 12.12 |
|  |  |  | UNI03 | 13.53 ± 2.83 | 25.14 ± 0.89 | 9 ± 1.78 | 15.67 ± 0.81 |
|  |  |  | UNI04 | 11.72 ± 1.17 | 20.55 ± 0.44 | 12.09 ± 0.77 | 38.82 ± 6.26 |
|  |  |  | UNI05 | 11.95 ± 2.26 | 23.85 ± 1.17 | 9.95 ± 0.77 | 33.55 ± 2.2 |
|  |  |  | UNI06 | 12.46 ± 2.92 | 28.29 ± 2.82 | 10.2 ± 0.36 | 42.57 ± 15.64 |
|  |  |  | UNI07 | 13.04 ± 2.65 | 27.59 ± 0.21 | 10.51 ± 4.41 | 37.51 ± 5.89 |
|  |  |  | UNI08 | 12.31 ± 2.06 | 37.43 ± 3.76 | 9.87 ± 3 | 27.68 ± 6.92 |
|  |  |  | UNI09 | 9.6 ± 0.84 | 32.7 ± 6.36 | 9.8 ± 2.8 | 25.55 ± 3.54 |
|  |  |  | UNI10 | 10.5 ± 2.56 | 47.78 ± 3.39 | 5.86 ± 2.07 | 37.22 ± 6.93 |
|  |  |  | UNI11 | 10.35 ± 1.97 | 28.68 ± 4.61 | 10.28 ± 0.28 | 39.06 ± 12.39 |

**Table S11.** Performance of *Uropsilus soricipes* under different experimental treatments and conditions.

| Test groups | Treatment | Experiment apparatus | Sample # | Percentage of exploration time in the left of monitored sector (%) | Percentage of exploration time in monitored sector (%) | Percentage of exploration time in the opposite sector (%) | # of ultrasonic emissions per sec in monitored sector |
| --- | --- | --- | --- | --- | --- | --- | --- |
| Small disks | None | Disc-small discs | USO01 | 13.87 ± 3.91 | 10.3 ± 0.64 | 13.96 ± 2.91 | 18.55 ± 2.75 |
|  |  |  | USO02 | 9.53 ± 2.96 | 13.5 ± 1.66 | 12.21 ± 2.04 | 17.68 ± 1.17 |
|  |  |  | USO03 | 9.53 ± 1.14 | 12.55 ± 0.96 | 11.15 ± 2.19 | 22.22 ± 7.31 |
|  |  |  | USO04 | 12.76 ± 1.16 | 10.29 ± 0.68 | 11.87 ± 1.25 | 16.42 ± 1.61 |
|  |  |  | USO05 | 10.24 ± 1.55 | 11.28 ± 0.69 | 11.84 ± 1.09 | 20.2 ± 3.03 |
|  |  |  | USO06 | 10.84 ± 1.69 | 11.24 ± 0.98 | 10.81 ± 2.36 | 22.91 ± 1.88 |
|  |  |  | USO07 | 11.06 ± 1.48 | 11.62 ± 0.61 | 11.34 ± 1.67 | 16.85 ± 3.21 |
|  |  |  | USO08 | 14.27 ± 0.94 | 13.46 ± 1.02 | 11.34 ± 1.19 | 15.24 ± 2.75 |
|  |  |  | USO09 | 13.65 ± 2.46 | 13.3 ± 0.95 | 12.36 ± 1.76 | 19.28 ± 3.64 |
|  |  |  | USO10 | 13.08 ± 3.19 | 11.9 ± 1.2 | 15.42 ± 3.57 | 22.34 ± 4.32 |
|  |  |  | USO11 | 12.52 ± 4.8 | 10.51 ± 0.36 | 13.24 ± 2.34 | 16.14 ± 2.11 |
|  |  |  | USO12 | 17.34 ± 3.98 | 13.21 ± 1.63 | 11.52 ± 2.59 | 18.45 ± 3.27 |
|  |  |  | USO13 | 14.35 ± 1.71 | 14.24 ± 0.76 | 9.99 ± 0.9 | 19.71 ± 4.15 |
|  |  |  | USO14 | 10.61 ± 2.8 | 10.57 ± 1.3 | 9.97 ± 0.91 | 25.22 ± 7.1 |
| Platform | None | Disc-platform | USO01 | 13 ± 1.12 | 28.8 ± 2.74 | 5.98 ± 1.87 | 28.7 ± 2.7 |
|  |  |  | USO02 | 17.72 ± 2.75 | 27.65 ± 2.39 | 6.42 ± 1.5 | 26.54 ± 3.05 |
|  |  |  | USO03 | 10.39 ± 1.89 | 33.74 ± 4.55 | 7.81 ± 1.47 | 29.89 ± 2.78 |
|  |  |  | USO04 | 14.22 ± 2.65 | 25.5 ± 3.82 | 5.79 ± 1.73 | 22.57 ± 4.5 |
|  |  |  | USO05 | 15.34 ± 0.82 | 26.86 ± 2.16 | 8.27 ± 1.65 | 23.29 ± 1.96 |
|  |  |  | USO06 | 19.33 ± 1.88 | 26.61 ± 2.04 | 7.36 ± 2.37 | 29.42 ± 2.65 |
|  |  |  | USO07 | 23.15 ± 8.36 | 28.73 ± 2 | 5.16 ± 1.48 | 36.33 ± 3.23 |
|  |  |  | USO08 | 17.56 ± 3.17 | 34.4 ± 7.11 | 7.4 ± 2.63 | 26.05 ± 5.91 |
|  |  |  | USO09 | 14.55 ± 1.54 | 26.36 ± 2.24 | 9.3 ± 1.49 | 30.01 ± 6.21 |
|  |  |  | USO10 | 15.08 ± 3.43 | 26.6 ± 2.35 | 9.11 ± 1.03 | 20.49 ± 3.81 |
|  |  |  | USO11 | 16.35 ± 3.36 | 22.84 ± 1.33 | 7.86 ± 1.86 | 32.29 ± 4.25 |
|  |  |  | USO12 | 13.5 ± 0.71 | 27.41 ± 4.2 | 8.16 ± 0.54 | 25.05 ± 3.31 |
|  |  |  | USO13 | 11.71 ± 3.49 | 25.82 ± 3.01 | 6.58 ± 1.79 | 33.57 ± 4.39 |
|  |  |  | USO14 | 12.88 ± 1.69 | 27.92 ± 2.22 | 9.54 ± 3.28 | 28.75 ± 6.78 |
| Earplug | Ears plugged | Disc-platform | USO01 | 19.21 ± 3.71 | 10.35 ± 0.65 | 10.56 ± 2.91 | 19.37 ± 4.93 |
|  |  |  | USO02 | 15.23 ± 3.26 | 12.71 ± 1.1 | 10.79 ± 5.21 | 20.31 ± 5.36 |
|  |  |  | USO03 | 9.95 ± 1.78 | 11.48 ± 0.75 | 15.44 ± 2.97 | 14.02 ± 2.34 |
|  |  |  | USO04 | 10.7 ± 1.47 | 13.82 ± 0.36 | 12.59 ± 2.59 | 22.53 ± 1.55 |
|  |  |  | USO05 | 15.63 ± 3.81 | 11.48 ± 1.2 | 12.48 ± 1.42 | 13.92 ± 3.55 |
|  |  |  | USO06 | 11.05 ± 4.15 | 11.68 ± 1.54 | 10.02 ± 1.98 | 18.78 ± 5.55 |
|  |  |  | USO07 | 11.6 ± 3.19 | 10.89 ± 0.91 | 11.92 ± 2.81 | 19.78 ± 3.48 |
|  |  |  | USO08 | 12.41 ± 1.6 | 12.68 ± 1.21 | 11.63 ± 1.04 | 15.65 ± 3.97 |
|  |  |  | USO09 | 14.9 ± 3.64 | 14.74 ± 0.85 | 10.5 ± 1.11 | 15.52 ± 1.96 |
|  |  |  | USO10 | 13.74 ± 1.42 | 11.16 ± 0.88 | 13.64 ± 1.94 | 19.94 ± 5.89 |
|  |  |  | USO11 | 18.58 ± 4.2 | 12.19 ± 0.53 | 10.92 ± 2.35 | 18.36 ± 4.67 |
|  |  |  | USO12 | 10.21 ± 0.82 | 12.55 ± 1.31 | 12.78 ± 2.8 | 17.52 ± 3.96 |
|  |  |  | USO13 | 12.52 ± 2.24 | 12.26 ± 1.19 | 12.57 ± 1.45 | 16.17 ± 2.01 |
|  |  |  | USO14 | 9.11 ± 1.67 | 12.18 ± 0.5 | 14.62 ± 2.76 | 18.91 ± 2.96 |
| Sham | Hollow tubes | Disc-platform | USO01 | 11.89 ± 1.53 | 26.3 ± 2.61 | 10.09 ± 1.8 | 26.91 ± 7.08 |
|  |  |  | USO02 | 14.48 ± 4.66 | 25.63 ± 1.82 | 6.8 ± 1.8 | 28.25 ± 1.47 |
|  |  |  | USO03 | 10.82 ± 0.85 | 27.99 ± 4.02 | 8.69 ± 0.5 | 25.29 ± 3.85 |
| Sham | Hollow tubes | Disc-platform | USO04 | 12.84 ± 1.15 | 29.5 ± 4.71 | 6.12 ± 0.54 | 22.87 ± 3.74 |
|  |  |  | USO05 | 8.82 ± 2.71 | 31.45 ± 6.38 | 10.1 ± 0.93 | 21.41 ± 1.01 |
|  |  |  | USO06 | 9.11 ± 2.37 | 38.24 ± 5.81 | 7.41 ± 0.93 | 37.21 ± 2.48 |
|  |  |  | USO07 | 20 ± 3.33 | 32.42 ± 3.28 | 4.03 ± 1.87 | 35.47 ± 3.27 |
|  |  |  | USO08 | 12.78 ± 4.59 | 30.47 ± 3.13 | 7 ± 2.33 | 31.01 ± 3.34 |
|  |  |  | USO09 | 14.33 ± 3.12 | 27.55 ± 2.86 | 5.48 ± 1.32 | 23.54 ± 5.34 |
|  |  |  | USO10 | 13.83 ± 2.39 | 30.46 ± 4.2 | 6.63 ± 1.11 | 25.26 ± 4.78 |
|  |  |  | USO11 | 14.99 ± 1.92 | 28.41 ± 2.09 | 9.93 ± 1.8 | 32.68 ± 4.74 |
|  |  |  | USO12 | 13.13 ± 2.71 | 23.24 ± 0.86 | 9.03 ± 4.55 | 23.46 ± 1.82 |
|  |  |  | USO13 | 12.54 ± 1.02 | 28.7 ± 2.44 | 10.39 ± 2.15 | 29.56 ± 0.74 |
|  |  |  | USO14 | 11.65 ± 1.3 | 30.09 ± 4.39 | 9.53 ± 0.44 | 25.98 ± 2.83 |
| Recovery | None | Disc-platform | USO01 | 12.38 ± 5.1 | 29.83 ± 7.39 | 4.67 ± 1.23 | 37.67 ± 4.15 |
|  |  |  | USO02 | 13.81 ± 6.11 | 37.5 ± 5.79 | 6.06 ± 4.29 | 32.2 ± 11.51 |
|  |  |  | USO03 | 12.07 ± 4.14 | 27.42 ± 5.2 | 9.24 ± 2.6 | 23.39 ± 2.36 |
|  |  |  | USO04 | 13.29 ± 4.83 | 31.05 ± 3.75 | 9.85 ± 3.47 | 36.04 ± 2.33 |
|  |  |  | USO05 | 11.63 ± 6.43 | 29.47 ± 4.21 | 10.17 ± 6.6 | 39.07 ± 5.04 |
|  |  |  | USO06 | 8.2 ± 2.14 | 32.48 ± 4.87 | 10.01 ± 3.75 | 33.26 ± 6.45 |
|  |  |  | USO07 | 10.88 ± 2.18 | 39.62 ± 8.99 | 7.67 ± 1.38 | 25.42 ± 1.99 |
|  |  |  | USO08 | 6.14 ± 2.17 | 32.3 ± 5.13 | 16.24 ± 3.64 | 27.33 ± 1.02 |
|  |  |  | USO09 | 11.65 ± 2.85 | 21.75 ± 3.55 | 9.69 ± 1.85 | 28.64 ± 6.58 |
|  |  |  | USO10 | 14.42 ± 5.49 | 22.18 ± 0.82 | 8.68 ± 1.52 | 25.28 ± 1.9 |
|  |  |  | USO11 | 11.68 ± 3.14 | 20.18 ± 0.58 | 8.11 ± 1.5 | 22.19 ± 1.01 |
|  |  |  | USO12 | 15.97 ± 3.72 | 20.88 ± 0.32 | 10.99 ± 1.52 | 39.46 ± 4.64 |
|  |  |  | USO13 | 11.21 ± 1.85 | 30.96 ± 10.4 | 8.41 ± 3.28 | 31.26 ± 3.9 |
|  |  |  | USO14 | 13.66 ± 3.1 | 27.89 ± 6.61 | 7.97 ± 2.44 | 26.96 ± 5.87 |

**Table S12.** Descriptive statistics for acoustic variables of ultrasonic pulses from echolocating species.

| Scientific name | Duration<br>(ms) | $f_{\min}$<br>(kHz) | $f_{\max}$<br>(kHz) | Bandwidth<br>(kHz) | Peak<br>frequency<br>(kHz) | Echolocation<br>calls type | Reference |
| --- | --- | --- | --- | --- | --- | --- | --- |
| *Anoura geoffroyi* | 2.08 | 66.3 | 105.87 | 39.57 | 83.08 | FM | 62 |
| *Antrozous pallidus* | 5.63 | 28.21 | 62.23 | 34.02 | 36.46 | FM | 62 |
| *Arielulus circumdatus* | 3.7 | 37.7 | 100.6 | 62.9 | 62.3 | FM | our data |
| *Artibeus jamaicensis* | 2.45 | 44.01 | 69.25 | 25.24 | 57.04 | FM | 62 |
| *Artibeus lituratus* | 2.04 | 49.92 | 76.68 | 26.76 | 61.44 | FM | 62 |
| *Artibeus lituratus* | 2.04 | 49.92 | 76.68 | 26.76 | 61.44 | FM | 63 |
| *Balantiopteryx infusca* | 6.7 | 51.1 | 57 | 5.9 | 55.7 | FM | 64 |
| *Balantiopteryx io* | 7.8 | 45.9 | 50.2 | 4.3 | 49.1 | FM | 64 |
| *Balantiopteryx plicata* | 12.1 | 39.4 | 42.2 | 2.8 | 41.2 | FM | 64 |
| *Balantiopteryx plicata* | 7.94 | 39.1 | 43.72 | 4.62 | 43.26 | FM | 63 |
| *Barbastella barbastellus* | 3.4 | 28 | 39.4 | 11.4 | 33.2 | FM | 63 |
| *Brachyphylla cavernarum* | 2.6 | 38 | 66.8 | 28.8 | 51.4 | FM | 63 |
| *Carollia perspicillata* | 2.42 | 45.67 | 75.37 | 29.69 | 62.66 | FM | 62 |
| *Centronycteris centralis* | 5.9 | NA | NA | NA | 41.3 | FM | 63 |
| *Chaerephon johorensis* | 17.5 | 14.1 | 21.8 | 7.7 | 16.7 | FM | 63 |
| *Chaerephon nigeriae* | 10 | NA | NA | NA | 17 | FM | 63 |
| *Chaerephon pumilus* | 13.6 | NA | NA | NA | 23.9 | FM | 63 |
| *Coelops frithii* | 11.8 | 112.5 | 173.7 | 61.2 | 132.9 | FM | 65 |
| *Coleura afra* | 7.7 | NA | NA | NA | 31.2 | FM | 63 |
| *Cormura brevirostris* | 8.2 | NA | NA | NA | 25.2 | FM | 63 |
| *Corynorhinus mexicanus* | 3.66 | 23.77 | 45.07 | 21.31 | 34.92 | FM | 62 |
| *Corynorhinus townsendii* | 2.27 | 24.47 | 41.53 | 17.06 | 33.75 | FM | 62 |
| *Craseonycteris thonglongyai* | 3.5 | NA | NA | NA | 73.2 | FM | 63 |
| *Cyttarops alecto* | 9.8 | NA | NA | NA | 35.9 | FM | 63 |
| *Desmodus rotundus* | 2.54 | 43.98 | 72.35 | 28.37 | 56.87 | FM | 62 |
| *Dididurus albus* | 9.4 | NA | NA | NA | 23.5 | FM | 63 |
| *Emballonura monticola* | 5.42 | 38.91 | 53.55 | 14.64 | 51.24 | FM | 63 |
| *Eptesicus bottae* | 6.19 | 30.51 | 42.63 | 12.12 | 32.08 | FM | 63 |
| *Eptesicus brasiliensis* | 7.8 | 32.93 | 55.83 | 22.89 | 37.12 | FM | 62 |
| *Eptesicus capensis* | 3.3 | 36.7 | 67.5 | 30.8 | 39.8 | FM | 66 |
| *Eptesicus furinalis* | 6.91 | 37.05 | 56.34 | 19.29 | 39.77 | FM | 62 |
| *Eptesicus fuscus* | 2.5 | 24.8 | 45.9 | 21.1 | 34.6 | FM | 67 |
| *Eptesicus melckorum* | 2.8 | 38.3 | 56.2 | 18 | 41 | FM | 66 |
| *Eptesicus nilssonii* | 6.3 | 26.1 | 57.9 | 31.8 | 30.5 | FM | 63 |
| *Eptesicus serotinus* | 7.3 | 27.1 | 50.4 | 23.3 | 29.9 | FM | 63 |
| *Erophylla bombifrons* | 4.7 | 26.8 | 54.2 | 27.4 | 37.9 | FM | 63 |
| *Erophylla sezekorni* | 2.3 | NA | NA | NA | 65.3 | FM | 63 |
| *Euderma maculatum* | 5 | NA | NA | NA | 10.9 | FM | 63 |
| *Eumops glaucinus* | 14.2 | NA | NA | NA | 17.9 | FM | 63 |
| *Eumops underwoodi* | 14.22 | 15.4 | 18.09 | 2.69 | 16.57 | FM | 68 |
| *Furipterus horrens* | 2.3 | NA | NA | NA | 157.2 | FM | 63 |
| *Glauconycteris variegata* | 5.3 | NA | NA | NA | 37.5 | FM | 63 |
| *Glossophaga soricina* | 2 | NA | NA | NA | 94.5 | FM | 63 |
| *Ia io* | 3.8 | 18 | 38.6 | 20.6 | 24.8 | FM | 63 |
| *Idionycteris phyllotis* | 4.28 | 13.83 | 30 | 16.17 | 21.04 | FM | 62 |
| *Kerivoula argentata* | 2 | 85 | 120 | 35 | 90 | FM | 66 |
| *Kerivoula hardwickii* | 3.15 | 90.73 | 169.57 | 78.84 | 118.25 | FM | 63 |
| *Kerivoula kachinensis* | 3.3 | 84 | 132 | 48 | 109.2 | FM | our data |
| *Lasionycteris noctivagans* | 3.29 | 39.49 | 90.03 | 50.54 | 53.75 | FM | 62 |
| *Lasiurus blossevillii* | 8.83 | 37.84 | 55.45 | 17.62 | 40.36 | FM | 62 |
| *Lasiurus borealis* | 4.25 | 26.79 | 57.24 | 30.45 | 35.47 | FM | 62 |
| *Lasiurus cinereus* | 2.93 | 37.45 | 61.13 | 23.68 | 55.2 | FM | 62 |
| *Lasiurus ega* | 9.53 | 26.14 | 49.39 | 23.25 | 30.04 | FM | 62 |
| *Lasiurus egregius* | 4.8 | 27.3 | 45.4 | 18.1 | 30.2 | FM | 63 |
| *Lasiurus intermedius* | 9.8 | NA | NA | NA | 28.3 | FM | 63 |
| *Lasiurus varius* | 7.64 | 33.18 | 52.25 | 19.07 | 36.44 | FM | 63 |
| *Lasiurus xanthinus* | 5.52 | 33.13 | 62.78 | 29.65 | 38.47 | FM | 62 |
| *Lavia frons* | 3.25 | NA | NA | NA | 42.21 | FM | 63 |
| *Leptonycteris yerbabuenae* | 6.22 | 34.12 | 79.28 | 45.16 | 49.72 | FM | 62 |
| *Macrophyllum macrophyllum* | 2.3 | NA | NA | NA | 55 | FM | 63 |
| *Macrotus californicus* | 3.31 | 47.45 | 80.44 | 32.99 | 60.39 | FM | 62 |
| *Megaderma lyra* | 2.6 | 32.9 | 56.4 | 23.5 | 42.5 | FM | 63 |
| *Megaderma spasma* | 2.06 | 38.87 | 99.79 | 60.92 | 55.9 | FM | 63 |
| *Miniopterus australis* | 4.42 | 47.2 | 95.4 | 48.2 | 61.46 | FM | 63 |
| *Miniopterus fraterculus* | 3.7 | NA | NA | NA | 62.1 | FM | 63 |
| *Miniopterus magnater* | 4.9 | 39 | 117.6 | 78.6 | 46.5 | FM | our data |
| *Miniopterus natalensis* | 3.4 | NA | NA | NA | 51.4 | FM | 63 |
| *Miniopterus pusillus* | 4.65 | 52.28 | 106.17 | 53.89 | 62.85 | FM | 63 |
| *Miniopterus schreibersii* | 5.8 | 52.1 | 85.2 | 33.1 | 54.2 | FM | 63 |
| *Molossus molossus* | 8.72 | 34.71 | 38.93 | 4.22 | 38.38 | FM | 62 |
| *Molossus rufus* | 9.73 | 28.79 | 32.28 | 3.49 | 31.89 | FM | 62 |
| *Molossus sinaloae* | 9.54 | 33.48 | 36.29 | 2.8 | 35.99 | FM | 62 |
| *Mops condylurus* | 10 | NA | NA | NA | 26.7 | FM | 63 |
| *Mops mops* | 16.7 | 15.5 | 25.9 | 10.4 | 18.5 | FM | 63 |
| *Mops niveiventer* | 8.1 | NA | NA | NA | 20.3 | FM | 63 |
| *Mormoops blainvillei* | 2.42 | NA | NA | NA | 61.03 | FM | 63 |
| *Mormoops megalophylla* | 4.78 | 42.1 | 55.66 | 13.56 | 52 | FM | 62 |
| *Murina leucogaster* | 3.5 | NA | NA | NA | 45.7 | FM | 63 |
| *Murina suilla* | 2.91 | 82.42 | 142.62 | 60.2 | 101.93 | FM | 63 |
| *Myotis adversus* | 4.68 | NA | NA | NA | 46.2 | FM | 63 |
| *Myotis albescens* | 2.5 | NA | NA | NA | 43 | FM | 63 |
| *Myotis auriculus* | 4.62 | 31.5 | 76.33 | 44.83 | 43.74 | FM | 62 |
| *Myotis bechsteinii* | 4.6 | 35.6 | 92.8 | 57.2 | 44.3 | FM | 63 |
| *Myotis blythii* | 4.3 | 30.4 | 74.4 | 44 | 41.4 | FM | 63 |
| *Myotis brandtii* | 3.06 | 33.7 | 85.5 | 51.8 | 47.9 | FM | 63 |
| *Myotis californicus* | 3.74 | 41.87 | 88.85 | 46.98 | 55.18 | FM | 62 |
| *Myotis capaccinii* | 4.6 | 35.1 | 89.7 | 54.6 | 48.9 | FM | 63 |
| *Myotis chinensis* | 3.6 | 25.2 | 83.6 | 58.4 | 49 | FM | 63 |
| *Myotis dasycneme* | 8.98 | NA | NA | NA | 33.79 | FM | 63 |
| *Myotis daubentonii* | 4.2 | 30.5 | 88 | 57.5 | 49.3 | FM | 63 |
| *Myotis emarginatus* | 3.6 | 41.2 | 109 | 67.8 | 58 | FM | 63 |
| *Myotis horsfieldii* | 3.17 | 39.63 | 87.25 | 47.62 | 56.93 | FM | 63 |
| *Myotis ikonnikovi* | 4 | NA | NA | NA | 51.2 | FM | 63 |
| *Myotis keaysi* | 3.3 | 58.71 | 88.47 | 29.76 | 60.87 | FM | 62 |
| *Myotis leibii* | 5 | NA | NA | NA | 44 | FM | 63 |
| *Myotis longipes* | 3.5 | 62.9 | 90.7 | 27.8 | 68.2 | FM | our data |
| *Myotis lucifugus* | 5 | NA | NA | NA | 45 | FM | 63 |
| *Myotis macrodactylus* | 6.27 | 42.31 | 73.25 | 30.94 | 49.57 | FM | 63 |
| *Myotis melanorhinus* | 2.77 | 39.83 | 99.06 | 59.23 | 56.55 | FM | 62 |
| *Myotis myotis* | 4.6 | 27.9 | 79.6 | 51.7 | 39.1 | FM | 63 |
| *Myotis mystacinus* | 4.2 | 32.4 | 96.4 | 64 | 47.5 | FM | 63 |
| *Myotis nattereri* | 4.7 | 24.4 | 111.8 | 87.4 | 46.9 | FM | 63 |
| *Myotis nigricans* | 5.75 | 51.25 | 78.45 | 27.2 | 54.6 | FM | 63 |
| *Myotis pequinius* | 5.72 | NA | NA | NA | 32.8 | FM | 63 |
| *Myotis pilosus* | 2.75 | 33.05 | 67.39 | 34.34 | 45.9 | FM | our data |
| *Myotis thysanodes* | 3.71 | 17.29 | 68.92 | 51.63 | 29.02 | FM | 62 |
| *Myotis tricolor* | 3.3 | NA | NA | NA | 47.8 | FM | 63 |
| *Myotis velifer* | 4.67 | 38.12 | 80.7 | 42.58 | 46.23 | FM | 62 |
| *Myotis volans* | 5.12 | 37.11 | 84.39 | 47.29 | 47.64 | FM | 62 |
| *Myotis yumanensis* | 3.69 | 42.76 | 87.56 | 44.8 | 52.64 | FM | 62 |
| *Mystacina tuberculata* | 3.5 | NA | NA | NA | 45.5 | FM | 63 |
| *Natalus stramineus* | 2.11 | 94.99 | 156.14 | 61.15 | 119.4 | FM | 62 |
| *Neoromicia capensis* | 5.1 | NA | NA | NA | 39.4 | FM | 63 |
| *Neoromicia nana* | 4 | NA | NA | NA | 70 | FM | 63 |
| *Nyctalus leisleri* | 5.3 | 27.7 | 55 | 27.3 | 30.7 | FM | 63 |
| *Nycteris grans* | 1.7 | 17 | 110 | 93 | 20 | FM | 66 |
| *Nycteris thebaica* | 2 | 61 | 97 | 36 | 94 | FM | 66 |
| *Nycteris tragata* | 2.87 | 71.13 | 111.88 | 40.75 | 97.64 | FM | 63 |
| *Nycteris woodi* | 2 | 35 | 55 | 20 | 43 | FM | 66 |
| *Nycticeinops schlieffeni* | 3.5 | NA | NA | NA | 42.5 | FM | 63 |
| *Nycticeius humeralis* | 5.55 | 17.08 | 36.55 | 19.46 | 24.38 | FM | 62 |
| *Nyctiellus lepidus* | 2.7 | NA | NA | NA | 83.4 | FM | 63 |
| *Nyctinomops femorosaccus* | 6.16 | 38.5 | 67.4 | 28.9 | 42.44 | FM | 62 |
| *Nyctinomops laticaudatus* | 4.85 | 17.96 | 41.03 | 23.07 | 25.68 | FM | 62 |
| *Nyctinomops laticaudatus* | 4.85 | 17.97 | 41.03 | 23.06 | 25.68 | FM | 63 |
| *Nyctinomops macrotis* | 7.92 | 13.79 | 28.66 | 14.86 | 22.34 | FM | 62 |
| *Otomops martiensseni* | 27 | NA | NA | NA | 11.8 | FM | 63 |
| *Otonycteris hemprichii* | 5.19 | 20.94 | 36.86 | 15.92 | 24.45 | FM | 63 |
| *Peropteryx kappleri* | 9.6 | NA | NA | NA | 31.6 | FM | 63 |
| *Peropteryx macrotis* | 7.3 | 37.54 | 41.87 | 4.33 | 41.61 | FM | 62 |
| *Phyllonycteris poeyi* | 4.69 | NA | NA | NA | 38.74 | FM | 63 |
| *Phyllops falcatus* | 4.2 | 23.9 | 73.9 | 50 | 56.2 | FM | 63 |
| *Phyllostomus hastatus* | 2.7 | NA | NA | NA | 47.1 | FM | 63 |
| *Pipistrellus abramus* | 7.94 | 43.11 | 55.4 | 12.29 | 44.81 | FM | 63 |
| *Pipistrellus affinis* | 2.3 | 55.1 | 128 | 72.9 | 77.9 | FM | 69 |
| *Pipistrellus alaschanicus* | 7.8 | NA | NA | NA | 34 | FM | 63 |
| *Pipistrellus hesperidus* | 2.5 | NA | NA | NA | 59.1 | FM | 63 |
| *Pipistrellus hesperus* | 4.78 | 43.25 | 66.51 | 23.26 | 46.49 | FM | 62 |
| *Pipistrellus kuhlii* | 6.4 | 37.5 | 59.8 | 22.3 | 39.7 | FM | 63 |
| *Pipistrellus maderensis* | 6.7 | 43.7 | 50.9 | 7.2 | 44.6 | FM | 63 |
| *Pipistrellus nathusii* | 6.2 | 38.7 | 75.7 | 37 | 40.8 | FM | 63 |
| *Pipistrellus pipistrellus* | 5.9 | 46.6 | 68.8 | 22.2 | 46.9 | FM | 63 |
| *Pipistrellus pygmaeus* | 5.5 | 56.8 | 79.6 | 22.8 | 57.7 | FM | 63 |
| *Pipistrellus rueppellii* | 4.22 | 51.94 | 62.24 | 10.3 | 52.19 | FM | 63 |
| *Pipistrellus rusticus* | 4.5 | NA | NA | NA | 55.7 | FM | 63 |
| *Pipistrellus savii* | 8.1 | 32.8 | 47.3 | 14.5 | 34.6 | FM | 63 |
| *Pipistrellus stenopterus* | 13.8 | 28 | 42.8 | 14.8 | 31 | FM | 63 |
| *Pipistrellus subflavus* | 7.58 | 41.69 | 63.88 | 22.19 | 44.02 | FM | 62 |
| *Plecotus auritus* | 2.3 | 26 | 44.7 | 18.7 | 33.1 | FM | 63 |
| *Plecotus austriacus* | 3.8 | 23.6 | 41.4 | 17.8 | 32.6 | FM | 63 |
| *Plecotus macrobullaris* | 3.98 | 43.95 | 19.35 | 24.6 | 28.53 | FM | 63 |
| *Promops centralis* | 3.73 | 38.93 | 40.72 | NA | 40.41 | FM | 68 |
| *Pteronotus davyi* | 6.7 | 69.2 | 79.2 | 10 | 71.5 | FM | 70 |
| *Pteronotus gymnonotus* | 5.33 | 45.81 | 55.51 | 9.7 | 51.34 | FM | 62 |
| *Pteronotus personatus* | 5.71 | 64.12 | 82.88 | 18.76 | 70.53 | FM | 62 |
| *Rhinopoma hardwickii* | 8.57 | 32.72 | 35.85 | 3.13 | 33.99 | FM | 63 |
| *Rhinopoma microphyllum* | 8.37 | 28.99 | 30.21 |  | 29.42 | FM | 63 |
| *Rhogeessa aeneus* | 3.79 | 45.17 | 76.66 | 31.49 | 49.73 | FM | 62 |
| *Rhogeessa io* | 2.8 | NA | NA | NA | 52.4 | FM | 63 |
| *Rhogeessa parvula* | 3.27 | 42.8 | 83.35 | 40.55 | 51.06 | FM | 62 |
| *Rhynchonycteris naso* | 4.38 | 81.84 | 98.71 | 16.86 | 95.79 | FM | 62 |
| *Saccolaimus saccolaimus* | 12.2 | NA | NA | NA | 22.6 | FM | 63 |
| *Saccopteryx bilineata* | 9.7 | 44 | 46.4 | 2.4 | 45.7 | FM | 70 |
| *Saccopteryx leptura* | 7.5 | 51.4 | 54.5 | 3.1 | 53.6 | FM | 70 |
| *Scotoecus albobfuscus* | 3.3 | NA | NA | NA | 39.3 | FM | 63 |
| *Scotophilus dinganii* | 4.9 | NA | NA | NA | 33.6 | FM | 63 |
| *Scotophilus heathii* | 2.4 | 60.12 | 37.65 | 22.47 | 41.2 | FM | 63 |
| *Scotophilus kuhlii* | 3.2 | 41 | 117.4 | 76.4 | 52.8 | FM | 69 |
| *Scotophilus viridis* | 10 | NA | NA | NA | 40 | FM | 63 |
| *Stenoderma rufum* | 3.1 | 26 | 95 | 69 | 67.6 | FM | 63 |
| *Sturnira lilium* | 3.13 | 52.62 | 90.86 | 38.24 | 72.44 | FM | 62 |
| *Sturnira ludovici* | 3.99 | 49.06 | 82.43 | 33.37 | 68.65 | FM | 62 |
| *Tadarida aegyptiaca* | 15.5 | 16.67 | 24.44 | 7.77 | 19.44 | FM | 63 |
| *Tadarida brasiliensis* | 7.2 | 26.09 | 46.83 | 20.74 | 32.61 | FM | 62 |
| *Taphozous mauritanus* | 15 | 12 | 55 | 43 | 25 | FM | 66 |
| *Taphozous melanopogon* | 6.02 | 22.58 | 36.6 | 14.02 | 29.71 | FM | 63 |
| *Taphozous perforatus* | 4.8 | 29.9 | 35.9 | 6 | 32.5 | FM | 63 |
| *Thyroptera tricolor* | 2.76 | 43.5 | 66.38 | 22.88 | 53.09 | FM | 62 |
| *Tylonycteris robustula* | 3.3 | 41 | 93.75 | 52.75 | 51.03 | FM | 63 |
| *Vampyrus spectrum* | 2.8 | NA | NA | NA | 79.4 | FM | 63 |
| *Vespertilio sinensis* | 6.2 | 21.8 | 48.1 | 26.3 | 24.2 | FM | 63 |
| *Mystacina tuberculata* | 2.2 | 20.9 | 35 | 14.1 | 27.1 | FM | 71 |
| *Typhlomys chapensis* | 2.42 | 39.5 | 113.4 | 74 | 97 | FM | our data |
| *Typhlomys cinereus* | 1.77 | 34.8 | 118.6 | 83.7 | 98.1 | FM | our data |
| *Typhlomys daloushanensis* | 1.72 | 37.4 | 119.4 | 82.1 | 96.1 | FM | our data |
| *Typhlomys nanus* | 1.82 | 27.8 | 115.3 | 87.5 | 101.7 | FM | our data |
| *Cloeotis percivali* | 2.67 | 188.91 | 209.38 | 20.74 | 209.31 | CF | 72 |
| *Aselliscus tricuspidatus* | 3.31 | 100.1 | 112.99 | 12.81 | 112.82 | CF | 72 |
| *Pteronotus macleayii* | 4.03 | NA | NA | NA | 69.02 | CF | 63 |
| *Pteronotus davyi* | 4.6 | 51 | 70.3 | 19.3 | 67 | CF | 63 |
| *Aselliscus stoliczkanus* | 4.7 | 110.8 | 127.9 | 17.1 | 127.5 | CF | 63 |
| *Molossus rufus* | 5.59 | 26.69 | 41.91 | 12.94 | 38.41 | CF | 72 |
| *Pteronotus personatus* | 5.71 | 64.12 | 82.83 | 18.71 | 70.53 | CF | 63 |
| *Paracoelops megalotis* | 6.46 | 97.61 | 117.07 | 14.84 | 113.18 | CF | 72 |
| *Asellia patrizii* | 6.48 | 94.92 | 114.06 | 18.81 | 109.84 | CF | 72 |
| *Molossus aztecus* | 6.66 | 29.11 | 41.28 | 9.23 | 34.33 | CF | 72 |
| *Molossus currentium* | 6.83 | 28.08 | 39.59 | 8.89 | 33.18 | CF | 72 |
| *Molossus sinaloae* | 6.92 | 27.41 | 38.6 | 8.77 | 32.46 | CF | 72 |
| *Molossus barnesi* | 7.06 | 26.74 | 37.3 | 8.43 | 31.66 | CF | 72 |
| *Molossus coibensis* | 7.06 | 26.74 | 37.3 | 8.43 | 31.66 | CF | 72 |
| *Asellia tridens* | 7.97 | 106.96 | 117.74 | 10.78 | 116.6 | CF | 63 |
| *Hipposideros pratti* | 8.01 | 53.45 | 60.83 | 7.35 | 60.77 | CF | 72 |
| *Hipposideros coronatus* | 8.03 | 82.11 | 91.36 | 10.46 | 90.11 | CF | 72 |
| *Noctilio leporinus* | 8.41 | 23.52 | 50.96 | 27.43 | 31.03 | CF | 62 |
| *Trienops rufus* | 8.43 | 35.82 | 43.27 | 7.48 | 39.82 | CF | 72 |
| *Trienops persicus* | 8.5 | 73.33 | 86.23 | 10.83 | 83 | CF | 72 |
| *Anthops ornatus* | 8.51 | 58.56 | 74.06 | 10.21 | 66.75 | CF | 72 |
| *Hipposideros demissus* | 8.65 | 73.7 | 83.83 | 9.39 | 81.29 | CF | 72 |
| *Hipposideros durgadasi* | 8.65 | 73.7 | 83.83 | 9.39 | 81.29 | CF | 72 |
| *Hipposideros edwardshilli* | 8.65 | 73.7 | 83.83 | 9.39 | 81.29 | CF | 72 |
| *Hipposideros grandis* | 8.65 | 73.7 | 83.83 | 9.39 | 81.29 | CF | 72 |
| *Hipposideros hypophyllus* | 8.65 | 73.7 | 83.83 | 9.39 | 81.29 | CF | 72 |
| *Hipposideros madurae* | 8.65 | 73.7 | 83.83 | 9.39 | 81.29 | CF | 72 |
| *Hipposideros orbiculus* | 8.65 | 73.7 | 83.83 | 9.39 | 81.29 | CF | 72 |
| *Hipposideros pelingensis* | 8.65 | 73.7 | 83.83 | 9.39 | 81.29 | CF | 72 |
| *Hipposideros scutinares* | 8.65 | 73.7 | 83.83 | 9.39 | 81.29 | CF | 72 |
| *Hipposideros sorenseni* | 8.65 | 73.7 | 83.83 | 9.39 | 81.29 | CF | 72 |
| *Hipposideros sumbae* | 8.65 | 73.7 | 83.83 | 9.39 | 81.29 | CF | 72 |
| *Hipposideros thomensis* | 8.65 | 73.7 | 83.83 | 9.39 | 81.29 | CF | 72 |
| *Hipposideros coxi* | 8.68 | 73.41 | 83.56 | 9.35 | 80.96 | CF | 72 |
| *Hipposideros crumeniferus* | 8.68 | 73.41 | 83.56 | 9.35 | 80.96 | CF | 72 |
| *Hipposideros lamottei* | 8.92 | 70.6 | 80.89 | 8.97 | 78.02 | CF | 72 |
| *Hipposideros lankadiva* | 9.17 | 60.16 | 74.32 | 10.6 | 69.34 | CF | 72 |
| *Hipposideros abae* | 9.26 | 70.74 | 95.32 | 13.98 | 84.86 | CF | 72 |
| *Hipposideros muscinus* | 9.28 | 87.53 | 106.92 | 13.72 | 100.89 | CF | 72 |
| *Hipposideros lekaguli* | 9.6 | 45.84 | 50.78 | 4.91 | 50.71 | CF | 72 |
| *Hipposideros lylei* | 9.65 | 58.52 | 67.34 | 8.85 | 67.27 | CF | 72 |
| *Trienops menamena* | 10.1 | 82 | 94.2 | 15.6 | 93.2 | CF | 63 |
| *Noctilio albiventris* | 10.5 | 52.35 | 85.08 | 27.4 | 69.5 | CF | 72 |
| *Hipposideros diadema* | 11.12 | 51.27 | 60.19 | 8.89 | 59.98 | CF | 72 |
| *Hipposideros dinops* | 11.55 | 41.68 | 51.45 | 6.46 | 48.28 | CF | 72 |
| *Hipposideros inexpectatus* | 11.75 | 40.81 | 50.24 | 6.32 | 47.32 | CF | 72 |
| *Hipposideros inornatus* | 12 | 69 | 64 | 5 | 69.2 | CF | 72 |
| *Hipposideros vittatus* | 12 | 55.09 | 64.53 | 6.14 | 61 | CF | 72 |
| *Rhinonictis aurantia* | 12 | 100 | 115 | 15 | 114 | CF | 72 |
| *Trienops auritus* | 12.19 | 82.9 | 104.2 | 21.3 | 100.3 | CF | 63 |
| *Trienops furculus* | 12.2 | 81.5 | 104.7 | 23.2 | 103.4 | CF | 63 |
| *Hipposideros commersoni* | 12.3 | 53.5 | 67.7 | 14.2 | 66.6 | CF | 63 |
| *Macronycteris commersoni* | 14.2 | 58.3 | 73.8 | 15.5 | 64 | CF | 62 |
| *Hipposideros camerunensis* | 14.75 | 50.91 | 60.11 | 5.3 | 56.77 | CF | 72 |
| *Hipposideros cyclops* | 15.86 | 70.18 | 75.53 | 5.43 | 75.19 | CF | 72 |
| *Hipposideros wollastoni* | 18.39 | 88.12 | 92.08 | 3.88 | 92.03 | CF | 72 |
| *Rhinolophus trifolius* | 19.51 | 44.24 | 52.56 | 8.21 | 52.41 | CF | 72 |
| *Hipposideros stenotis* | 20 | 99 | 104 | 5 | 103 | CF | 72 |
| *Hipposideros gigas* | 20.9 | 52.22 | 59.45 | 7.24 | 59.28 | CF | 72 |
| *Pteronotus parnellii* | 21.21 | 52.86 | 64.97 | 12.1 | 63.61 | CF | 62 |
| *Rhinolophus euryale* | 21.82 | 91.24 | 106.91 | 15.58 | 105.67 | CF | 72 |
| *Rhinolophus denti* | 22.28 | 90.72 | 110.71 | 19.98 | 110.6 | CF | 72 |
| *Rhinolophus swinnyi* | 22.3 | 88.57 | 107.32 | 18.78 | 104.36 | CF | 72 |
| *Rhinolophus keyensis* | 24.22 | 78.96 | 92.91 | 13.95 | 90.47 | CF | 72 |
| *Hipposideros armiger* | 24.8 | 61.3 | 68.7 | 7.4 | 68.6 | CF | 65 |
| *Rhinolophus robinsoni* | 26.15 | 70.67 | 85.08 | 12.69 | 81.37 | CF | 72 |
| *Rhinolophus stheno* | 26.53 | 79.48 | 93.99 | 14.52 | 93.89 | CF | 72 |
| *Rhinolophus cognatus* | 26.79 | 68.24 | 82.74 | 12.31 | 78.73 | CF | 72 |
| *Rhinolophus celebensis* | 27.74 | 64.91 | 79.35 | 11.76 | 75.04 | CF | 72 |
| *Rhinolophus imaizumii* | 27.74 | 64.91 | 79.35 | 11.76 | 75.04 | CF | 72 |
| *Rhinolophus deckenii* | 28.28 | 61.25 | 76.42 | 11.87 | 72 | CF | 72 |
| *Rhinolophus macrotis* | 28.47 | 40.58 | 47 | 6.44 | 46.95 | CF | 72 |
| *Rhinolophus osgoodi* | 28.67 | 61.87 | 76.16 | 11.24 | 71.66 | CF | 72 |
| *Rhinolophus formosae* | 29.2 | 60.28 | 74.53 | 10.98 | 69.97 | CF | 72 |
| *Rhinolophus hillorum* | 29.2 | 60.28 | 74.53 | 10.98 | 69.97 | CF | 72 |
| *Rhinolophus madurensis* | 29.2 | 60.28 | 74.53 | 10.98 | 69.97 | CF | 72 |
| *Rhinolophus maendeleo* | 29.2 | 60.28 | 74.53 | 10.98 | 69.97 | CF | 72 |
| *Rhinolophus monoceros* | 29.2 | 93.6 | 108.7 | 15.1 | 108.2 | CF | 65 |
| *Rhinolophus montanus* | 29.2 | 60.28 | 74.53 | 10.98 | 69.97 | CF | 72 |
| *Rhinolophus rufus* | 29.2 | 60.28 | 74.53 | 10.98 | 69.97 | CF | 72 |
| *Rhinolophus sakejiensis* | 29.2 | 60.28 | 74.53 | 10.98 | 69.97 | CF | 72 |
| *Rhinolophus shortridgei* | 29.2 | 60.28 | 74.53 | 10.98 | 69.97 | CF | 72 |
| *Rhinolophus ziama* | 29.2 | 60.28 | 74.53 | 10.98 | 69.97 | CF | 72 |
| *Rhinolophus hilli* | 29.34 | 59.8 | 73.99 | 10.9 | 69.41 | CF | 72 |
| *Rhinolophus inops* | 29.37 | 59.74 | 73.93 | 10.88 | 69.34 | CF | 72 |
| *Rhinolophus nereis* | 29.37 | 59.74 | 73.93 | 10.88 | 69.34 | CF | 72 |
| *Rhinolophus simulator* | 29.37 | 66.2 | 80.76 | 14.59 | 80.71 | CF | 72 |
| *Rhinolophus bocharicus* | 30.08 | 57.69 | 71.69 | 10.52 | 67.09 | CF | 72 |
| *Rhinolophus guineensis* | 30.08 | 57.74 | 71.77 | 10.53 | 67.15 | CF | 72 |
| *Rhinolophus sedulus* | 30.5 | 59.91 | 67.92 | 8.07 | 67.87 | CF | 72 |
| *Rhinolophus siamensis* | 30.85 | 55.92 | 69.69 | 10.09 | 65 | CF | 72 |
| *Rhinolophus adami* | 31.22 | 54.76 | 68.43 | 9.99 | 63.82 | CF | 72 |
| *Rhinolophus canuti* | 31.44 | 54.16 | 67.77 | 9.88 | 63.18 | CF | 72 |
| *Rhinolophus borneensis* | 31.94 | 69.97 | 83.8 | 12.81 | 81.8 | CF | 72 |
| *Rhinolophus malayanus* | 32.04 | 71.68 | 85.62 | 13.95 | 85.55 | CF | 72 |
| *Rhinolophus lepidus* | 32.2 | 74.9 | 94.9 | 20 | 95.2 | CF | 69 |
| *Rhinolophus sinicus* | 32.33 | 71.74 | 86.84 | 13.46 | 84 | CF | 72 |
| *Rhinolophus silvestris* | 32.79 | 51.01 | 64.11 | 9.29 | 59.68 | CF | 72 |
| *Rhinolophus thomasi* | 33.89 | 74.82 | 89.61 | 14.8 | 89.53 | CF | 72 |
| *Rhinolophus creaghi* | 34.64 | 57.97 | 70.95 | 10.53 | 68 | CF | 72 |
| *Rhinolophus shameli* | 34.87 | 61.98 | 69.65 | 7.66 | 69.59 | CF | 72 |
| *Rhinolophus coelophyllus* | 35.01 | 66.29 | 78.77 | 12.5 | 78.46 | CF | 72 |
| *Rhinolophus arcuatus* | 35.34 | 57.28 | 69.76 | 10 | 66.5 | CF | 72 |
| *Rhinolophus alcyone* | 35.41 | 72.53 | 91.45 | 15.39 | 87 | CF | 72 |
| *Rhinolophus subbadius* | 35.45 | 58.67 | 71.64 | 9.98 | 67.29 | CF | 72 |
| *Rhinolophus mehelyi* | 35.9 | 97.42 | 89.93 | 5 | 110 | CF | 72 |
| *Rhinolophus darlingi* | 36.05 | 73.24 | 86.29 | 13.04 | 86.01 | CF | 72 |
| *Rhinolophus pusillus* | 36.08 | 97.72 | 109.93 | 12.18 | 109.85 | CF | 72 |
| *Rhinolophus mitratus* | 37.49 | 48.62 | 61.64 | 8.98 | 57.45 | CF | 72 |
| *Rhinolophus eloquens* | 38.28 | 36.6 | 45.79 | 6.71 | 43.29 | CF | 72 |
| *Rhinolophus virgo* | 38.44 | 59.86 | 70.49 | 9.34 | 67.49 | CF | 72 |
| *Rhinolophus convexus* | 38.63 | 62.68 | 75.74 | 10.47 | 72.02 | CF | 72 |
| *Rhinolophus ruwenzorii* | 39.73 | 39.57 | 48.92 | 6.76 | 46.57 | CF | 72 |
| *Rhinolophus hipposideros* | 40.83 | 94.59 | 109.47 | 14.83 | 109.32 | CF | 72 |
| *Rhinolophus fumigatus* | 41.27 | 46.57 | 53.28 | 6.64 | 53.24 | CF | 72 |
| *Rhinolophus rouxii* | 41.4 | 61.5 | 79.8 | 18.3 | 80 | CF | 69 |
| *Rhinolophus landeri* | 41.85 | 86.25 | 105.46 | 19.2 | 105.41 | CF | 72 |
| *Rhinolophus pearsonii* | 42.3 | 51.42 | 56.72 | 5.33 | 56.68 | CF | 72 |
| *Rhinolophus macclaudi* | 42.52 | 37.98 | 46.85 | 6.34 | 44.93 | CF | 72 |
| *Rhinolophus capensis* | 42.67 | 71.91 | 84.42 | 12.5 | 84.36 | CF | 72 |
| *Rhinolophus subrufus* | 42.73 | 44.08 | 54.08 | 7.27 | 51 | CF | 72 |
| *Rhinolophus affinis* | 43.2 | 56.6 | 56.9 | 15.78 | 71.1 | CF | 63 |
| *Rhinolophus rex* | 43.52 | 22.06 | 26.04 | 3.99 | 26.01 | CF | 72 |
| *Rhinolophus cornutus* | 43.58 | 92.67 | 106.05 | 13.35 | 105.97 | CF | 72 |
| *Rhinolophus blasii* | 44.1 | 54.33 | 58.31 | 4 | 60.5 | CF | 72 |
| *Rhinolophus hildebrandtii* | 46.62 | 28.5 | 34.31 | 5.79 | 34.27 | CF | 72 |
| *Rhinolophus ferrumequinum* | 48.09 | 69.81 | 82.34 | 12.55 | 82.18 | CF | 72 |
| *Rhinolophus paradoxolophus* | 48.76 | 38.71 | 46.23 | 5.07 | 44 | CF | 72 |
| *Rhinolophus acuminatus* | 48.86 | 80.97 | 89.61 | 8.63 | 89.56 | CF | 72 |
| *Rhinolophus yunnanensis* | 49.34 | 45.03 | 49.77 | 4.74 | 49.7 | CF | 72 |
| *Rhinolophus marshalli* | 49.45 | 38.36 | 45.55 | 4.82 | 43 | CF | 72 |
| *Rhinolophus euryotis* | 50.82 | 44.92 | 52.03 | 7.1 | 51.99 | CF | 72 |
| *Hipposideros semoni* | 51.43 | 60.39 | 67.5 | 7.13 | 67.46 | CF | 72 |
| *Rhinolophus beddomei* | 53.1 | 31.2 | 38.3 | 7.1 | 38.6 | CF | 69 |
| *Rhinolophus clivosus* | 54.48 | 78.81 | 78.92 | 12.5 | 85.18 | CF | 63 |
| *Rhinolophus megaphyllus* | 56.06 | 61.29 | 68.85 | 7.56 | 68.8 | CF | 72 |
| *Rhinolophus luctus* | 64.97 | 29.65 | 31.11 | NA | 31.07 | CF | 72 |
| *Rhinolophus philippinensis* | 81.84 | 35.1 | 39.85 | 4.71 | 39.78 | CF | 72 |
| *Rousettus leschenaulti* | 1.1 | 13.84 | 74.15 | 60.31 | 25.03 | click | our data |
| *Rousettus aegyptiacus* | 0.41 | 12 | 70 | 58 | 30 | click | 73, 74 |
| *Rousettus semirudus* | 0.8 | \ | \ | \ | \ | click | 75 |
| *Rousettus amplexicaudatus* | 0.9 | 13.5 | 65 | 51.5 | 32 | click | 75, 76 |
| *Echinops telfairi* | 0.9 | \ | \ | \ | 14 | click | 5 |
| *Hemicentetes semispinosus* | 0.4 | \ | \ | \ | 13 | click | 5 |
| *Microgale dobsoni* | 0.5 | \ | \ | \ | 11 | click | 5 |
| *Cephalorhynchus commersonii* | 0.078 | \ | \ | 21 | 133 | click | 77 |
| *Cephalorhynchus hectori* | 0.057 | \ | \ | 30 | 128 | click | 77 |
| *Globicephala melas* | 0.23 | \ | \ | 80 | 50 | click | 78 |
| *Grampus griseus* | 0.04 | \ | \ | 66 | 49 | click | 79 |
| *Hyperoodon ampullatus* | 0.276 | \ | \ | \ | 43 | click | 90 |
| *Inia geoffrensis* | 0.014 | \ | \ | 117.5 | 95.7 | click | 80 |
| *Kogia breviceps* | 0.6 | \ | \ | 140 | 125 | click | 81 |
| *Kogia sima* | 0.142 | \ | \ | 16 | 129 | click | 82 |
| *Lagenorhynchus albirostris* | 0.015 | \ | \ | 75 | 115 | click | 83 |
| *Lagenorhynchus obscurus* | 0.07 | \ | \ | 67.4 | 73.8 | click | 84 |
| *Lagenorhynchus cruciger* | 0.115 | \ | \ | 13 | 128 | click | 77 |
| *Lagenorhynchus australis* | 0.092 | \ | \ | 15 | 129 | click | 77 |
| *Mesoplodon densirostris* | 0.271 | \ | \ | 54.9 | 51.3 | click | 85 |
| *Orcaella brevirostris* | 0.013 | \ | \ | 117.9 | 100.7 | click | 86 |
| *Orcinus orca* | 0.41 | \ | \ | 44 | 29 | click | 90 |
| *Neophocaena phocaenoides asia* | 0.039 | \ | \ | 47 | 130 | click | 87 |
| *Peponocephala electra* | 0.46 | \ | \ | 40.9 | 25.5 | click | 88 |
| *Phocoena phocoena* | 0.175 | \ | \ | 46 | 142 | click | 90 |
| *Physeter macrocephalus* | 0.114 | \ | \ | 10 | 15 | click | 90 |
| *Platanista gangetica gangetica* | 0.022 | \ | \ | 73.2 | 58.8 | click | 86 |
| *Pseudorca crassidens* | 0.05 | \ | \ | 100 | 48 | click | 79 |
| *Pontoporia blainvillei* | 0.212 | \ | \ | 19 | 139 | click | 90 |
| *Sousa chinensis* | 0.019 | \ | \ | 102 | 109 | click | 89 |
| *Sousa plumbea* | 0.016 | \ | \ | 125 | 91 | click | 90 |
| *Sousa sahalensis* | 0.015 | \ | \ | 116 | 114 | click | 89 |
| *Stenella attenuata* | 0.043 | \ | \ | NA | 83.4 | click | 89 |
| *Stenella longirostris* | 0.21 | \ | \ | 68.9 | 33.8 | click | 88 |
| *Tursiops aduncus* | 0.014 | \ | \ | 140 | 124 | click | 89 |
| *Tursiops truncatus* | 0.072 | \ | \ | 100 | 102 | click | 89 |
| *Ziphius cavirostris* | 0.2 | \ | \ | 23 | 42 | click | 85 |

**Table S13.** The amino acid of 14 convergent amino acid substitutions across 97 mammals from predicting dataset.

| Convergent gene (site)<br>Species | SLC26A5<br>(497M) | OTOF<br>(1982F) | PLEKHH1<br>(517G) | COL11A1<br>(285A) | TBC1D2<br>(443S) | KMT2A<br>(2344L) | PHF3<br>(801D) | TRAFD1<br>(192E) | THSD1<br>(690P) | SCN7A<br>(910R) | EFL1<br>(772V) | ZC3H6<br>(573H) | ZNF316<br>(104S) | SH2B3<br>(548V) |
| --- | --- | --- | --- | --- | --- | --- | --- | --- | --- | --- | --- | --- | --- | --- |
| *Acomys cahirinus* | M | L | - | V | - | I | E | D | H | G | L | Q | P | I |
| *Alouatta palliata mexicana* | L | L | S | - | R | I | E | D | Q | G | * | Q | P | - |
| *Anoura caudifer* | M | L | S | V | S | I | E | D | P | G | L | H | - | I |
| *Antilocapra americana* | L | L | S | V | R | I | E | D | Q | R | L | - | - | I |
| *Antrozous pallidus* | M | F | S | A | K | L | - | E | Q | G | - | - | - | - |
| *Aotus nancymae* | L | L | S | V | R | I | D | D | Q | G | L | Q | P | I |
| *Aplodontia rufa* | L | L | S | V | R | I | E | D | Q | G | - | Q | - | I |
| *Ateles geoffroyi* | L | - | S | - | R | I | E | D | Q | G | L | Q | - | - |
| *Beatragus hunteri* | * | L | S | - | R | I | E | D | Q | G | - | - | - | I |
| *Callicebus donacophilus* | L | L | S | - | R | I | - | D | Q | G | - | Q | - | I |
| *Carlito syrichta* | M | L | S | V | R | I | E | D | Q | G | L | Q | * | I |
| *Castor canadensis* | * | L | S | V | M | I | E | D | Q | G | L | Q | - | - |
| *Catagonus wagneri* | L | L | S | V | R | A | E | D | Q | G | L | Q | - | I |
| *Cavia tschudii* | L | - | S | V | R | I | E | D | Q | K | - | Q | P | I |
| *Cheirogaleus medius* | L | L | S | - | R | I | E | D | Q | G | L | Q | P | I |
| *Chinchilla lanigera* | L | L | S | V | R | I | E | D | Q | G | L | Q | P | I |
| *Choloepus hoffmanni* | L | L | S | V | R | M | E | D | Q | G | L | P | L | I |
| *Cricetomys gambianus* | * | L | S | V | G | I | E | D | P | G | L | Q | * | I |
| *Cricetulus griseus* | L | L | S | L | G | I | E | D | Q | G | L | Q | P | I |
| *Cryptoprocta ferox* | L | L | S | * | R | V | E | D | Q | G | - | P | - | I |
| *Ctenodactylus gundi* | L | L | N | A | R | I | E | D | H | R | L | Q | P | I |
| *Ctenomys sociabilis* | L | L | S | V | R | I | E | D | Q | G | L | Q | P | I |
| *Dasyprocta punctata* | * | L | S | V | R | I | E | D | Q | G | L | Q | P | I |
| *Dasypus novemcinctus* | M | L | S | V | - | I | E | D | Q | R | L | R | P | I |
| *Daubentonia madagascariensis* | L | L | S | V | R | I | E | D | Q | G | L | Q | P | I |
| *Diceros bicornis* | L | L | S | V | R | I | E | D | Q | G | L | Q | P | I |
| *Dinomys branickii* | L | - | S | - | R | * | E | E | Q | G | - | Q | - | I |
| *Dipodomys ordii* | L | L | S | V | R | I | E | D | Q | G | L | Q | T | V |
| *Dipodomys stephensi* | * | - | S | * | R | I | - | D | Q | G | - | Q | T | V |
| *Dolichotis patagonum* | L | L | S | V | R | I | E | D | Q | G | L | Q | - | I |
| *Elephantulus edwardii* | L | L | S | V | R | I | E | D | Q | G | L | Q | P | I |
| *Erythrocebus patas* | L | - | S | - | - | I | E | D | Q | G | L | Q | P | - |
| *Eubalaena japonica* | L | L | S | - | R | I | E | D | - | G | L | Q | - | I |
| *Felis catus* | L | L | S | V | R | I | E | D | Q | G | L | Q | P | I |
| *Galeopterus variegatus* | L | - | S | V | R | I | E | D | Q | G | L | Q | * | I |
| *Helogale parvula* | L | L | S | V | K | V | E | D | Q | G | - | Q | - | I |
| *Hemitragus hylocrius* | L | L | N | - | R | I | E | D | Q | G | L | - | - | I |
| *Heterohyrax brucei* | L | - | S | - | R | I | E | D | - | G | - | Q | P | I |
| *Hippopotamus amphibius* | L | L | S | V | R | L | E | D | Q | G | L | Q | - | I |
| *Hipposideros galeritus* | M | L | S | * | R | I | E | D | Q | G | - | Q | - | V |
| *Hyaena hyaena* | L | - | S | V | - | I | E | D | Q | G | - | Q | - | I |
| *Hydrochoerus hydrochaeris* | L | P | S | V | R | I | E | E | Q | R | L | Q | P | I |
| *Hystrix cristata* | L | L | - | V | R | I | * | - | Q | G | - | Q | P | I |

| Convergent gene (site) | SLC26A5<br>(497M) | OTOF<br>(1982F) | PLEKHH1<br>(517G) | COL11A1<br>(285A) | TBC1D2<br>(443S) | KMT2A<br>(2344L) | PHF3<br>(801D) | TRAFD1<br>(192E) | THSD1<br>(690P) | SCN7A<br>(910R) | EFL1<br>(772V) | ZC3H6<br>(573H) | ZNF316<br>(104S) | SH2B3<br>(548V) |
| --- | --- | --- | --- | --- | --- | --- | --- | --- | --- | --- | --- | --- | --- | --- |
| Species |  |  |  |  |  |  |  |  |  |  |  |  |  |  |
| *Indri indri* | * | - | - | V | R | I | E | D | Q | G | L | Q | P | I |
| *Lemur catta* | L | L | S | V | R | I | E | D | Q | G | L | Q | P | I |
| *Macroglossus*<br>*sobrinus* | M | L | S | V | R | I | E | D | Q | G | L | Q | L | I |
| *Manis javanica* | L | L | S | A | R | I | E | D | A | G | L | Q | S | I |
| *Manis tricuspis* | L | L | S | - | M | I | E | D | - | G | - | Q | - | I |
| *Megaderma lyra* | M | L | S | V | R | L | E | D | Q | G | L | Q | P | I |
| *Mellivora capensis* | L | L | S | - | R | I | E | D | Q | G | - | Q | P | I |
| *Mesocricetus*<br>*auratus* | L | L | S | L | G | I | E | D | Q | G | L | Q | P | I |
| *Mesoplodon bidens* | M | L | S | - | - | I | * | E | E | G | L | Q | P | - |
| *Microcebus murinus* | M | L | S | V | R | I | E | D | Q | G | L | Q | P | I |
| *Micronycteris*<br>*hirsuta* | M | L | S | V | S | I | E | D | P | G | L | H | - | I |
| *Miniopterus*<br>*schreibersii* | M | L | S | G | R | I | D | D | Q | G | - | Q | P | I |
| *Mirounga*<br>*angustirostris* | L | L | - | * | R | I | E | D | Q | G | - | Q | P | I |
| *Mirza coquereli* | M | - | - | - | R | I | E | D | Q | G | L | Q | - | I |
| *Monachus*<br>*schauinslandi* | L | L | S | V | R | I | E | D | Q | R | L | Q | P | I |
| *Mormoops*<br>*blainvillei* | M | L | S | V | R | I | E | D | P | G | L | H | P | I |
| *Moschus*<br>*moschiferus* | L | L | S | V | R | I | E | D | Q | G | L | - | - | I |
| *Mungos mungo* | L | L | S | V | R | V | E | D | Q | G | L | Q | - | I |
| *Mus musculus* | L | L | S | V | G | I | E | D | Q | G | L | Q | P | I |
| *Muscardinus*<br>*avellanarius* | * | L | S | * | * | I | E | A | Q | G | L | Q | P | V |
| *Mustela putorius* | L | L | S | V | R | I | E | D | Q | G | L | Q | P | I |
| *Noctilio leporinus* | M | L | S | V | R | I | E | D | P | G | L | Q | P | I |
| *Ondatra zibethicus* | * | L | S | * | G | I | E | D | Q | G | * | Q | P | I |
| *Orycteropus afer* | L | L | S | V | R | S | E | D | Q | G | L | Q | P | I |
| *Otolemur garnettii* | L | L | S | V | R | I | E | D | H | G | L | Q | P | I |
| *Ovis canadensis*<br>*cremnobates* | L | L | S | V | R | I | E | D | Q | G | L | - | - | I |
| *Panthera onca* | L | L | - | V | R | I | - | D | Q | G | - | Q | P | I |
| *Panthera tigris* | L | L | S | V | R | I | E | D | Q | G | L | Q | - | I |
| *Paradoxurus*<br>*hermaphroditus* | L | * | S | * | - | I | E | D | Q | G | - | Q | - | I |
| *Pedetes capensis* | L | L | S | * | R | I | E | D | Q | G | - | Q | P | I |
| *Perognathus*<br>*longimembris* | L | L | - | M | P | I | E | D | Q | G | L | - | - | V |
| *Peromyscus*<br>*maniculatus* | L | L | G | L | G | I | E | D | Q | G | L | Q | P | I |
| *Petromus typicus* | L | L | S | - | R | I | - | D | Q | G | L | - | - | I |
| *Pithecia pithecia* | L | L | S | - | R | I | E | D | Q | G | L | Q | P | - |
| *Pteronura*<br>*brasiliensis* | L | L | S | V | R | I | E | D | Q | G | L | Q | - | I |
| *Pygathrix nemaeus* | L | L | S | V | * | I | E | D | Q | G | L | * | - | I |
| *Rangifer tarandus* | L | L | S | V | R | I | E | D | Q | G | - | - | P | I |
| *Rattus norvegicus* | L | L | S | V | G | I | E | D | Q | G | L | Q | P | I |
| *Rhinolophus*<br>*ferrumequinum* | M | F | S | V | R | I | E | D | Q | G | L | Q | P | V |
| *Saguinus imperator* | L | L | S | * | R | I | E | D | Q | G | * | Q | P | I |
| *Scalopus aquaticus* | L | L | S | - | R | I | E | D | Q | G | V | Q | - | I |
| *Sigmodon hispidus* | L | L | S | * | G | I | E | D | Q | G | - | Q | P | I |
| *Solenodon*<br>*paradoxus* | M | L | S | V | S | I | E | D | Q | G | L | Q | P | I |
| *Sorex araneus* | L | L | G | V | - | L | E | D | P | G | L | Q | - | I |
| *Spilogale gracilis* | L | L | S | - | R | I | E | P | Q | G | L | Q | P | I |
| *Suricata suricatta* | L | * | S | V | R | V | E | D | Q | G | - | Q | P | V |
| Species |  |  |  |  |  |  |  |  |  |  |  |  |  |  |
| *Sus scrofa* | L | L | S | V | R | T | E | D | Q | G | L | Q | P | I |
| *Tapirus indicus* | L | L | S | V | R | I | E | D | Q | G | L | Q | - | I |
| *Tapirus terrestris* | L | L | S | V | R | I | E | - | Q | G | L | R | P | I |
| *Tonatia saurophila* | M | L | S | V | S | V | * | D | P | G | L | H | - | V |
| *Tragulus javanicus* | L | L | S | V | R | I | E | D | Q | G | L | L | - | I |
| *Uropsilus gracilis* | M | L | S | V | R | I | E | D | Q | G | L | Q | S | V |
| *Vulpes lagopus* | L | - | S | V | R | I | E | D | Q | G | L | Q | - | I |
| *Xerus inauris* | L | L | S | * | R | I | E | D | Q | G | - | Q | P | I |
| *Zalophus californianus* | L | L | S | V | R | I | E | D | H | G | * | Q | - | I |

"\*"indicates that there is no one-to-one orthologous protein-coding gene sequence
"-"indicates that there is no amino acids in that location

